# Different learning algorithms achieve shared optimal outcomes in humans, rats, and mice

**DOI:** 10.1101/2023.01.30.526119

**Authors:** Elena Menichini, Quentin Pajot-Moric, Ryan Low, Victor Pedrosa, Amirali Pourdehghan, Peter Vincent, Liang Zhou, Lillianne Teachen, Athena Akrami

**Author notes:** Corresponding author: Athena Akrami. These authors contributed equally, and are listed alphabetically (based on first name) (Elena Menichini, Quentin Pajot-Moric, Ryan Low).

## Abstract

Animals must exploit environmental regularities to make adaptive decisions, yet the learning algorithms that enabels this flexibility remain unclear. A central question across neuroscience, cognitive science, and machine learning, is whether learning relies on generative or discriminative strategies. Generative learners build internal models the sensory world itself, capturing its statistical structure; discriminative learners map stimuli directly onto choices, ignoring input statistics. These strategies rely on fundamentally different internal representations and entail distinct computational trade-offs: generative learning supports flexible generalisation and transfer, whereas discriminative learning is efficient but task-specific. We compared humans, rats, and mice performing the same auditory categorisation task, where category boundaries and rewards were fixed but sensory statistics varied. All species adapted their behaviour near-optimally, consistent with a normative observer constrained by sensory and decision noise. Yet their underlying algorithms diverged: humans predominantly relied on generative representations, mice on discriminative boundary-tracking, and rats spanned both regimes. Crucially, end-point performance concealed these differences—only learning trajectories and trial-to-trial updates revealed the divergence. These results show that similar near-optimal behaviour can mask fundamentally different internal representations, establishing a comparative framework for uncovering the hidden strategies that support statistical learning.

## Introduction

Intelligent behaviour depends on discovering and exploiting structure in a noisy world (Fiser et al. 2010; Knill and Pouget 2004). Across species, animals continuously extract statistical regularities from sensory inputs to predict outcomes and guide decisions (Shettleworth and Shettleworth 2010; Schapiro and N. Turk-Browne 2015; N. B. Turk-Browne et al. 2010; T. L. Griffiths and Tenenbaum 2006). Yet despite extensive behavioural, theoretical, and neural work, it remains unknown whether different brains achieve this flexibility through shared or distinct learning algorithms.

Categorisation, the process of grouping variable sensory events into meaningful classes, captures the core computational challenge of learning structure from sensory experience. It transforms noisy, continuous inputs into discrete representations that support generalisation and adaptive decision-making (Britten et al. 1992; Anderson 1991). Such mappings are inherently context-dependent: identical cues can acquire different meanings depending on prior experience or environmental statistics. A fruit’s colour may indicate ripeness in one culture but immaturity in another, and the same wavelength of light may be labelled “blue” or “green” across languages (Kay and Regier 2006; Regier, Kay, and Khetarpal 2007). Likewise, in rats, ultrasonic vocalisations are interpreted differently depending on behavioural context and recent experience (Wöhr et al. 2017; Seffer, Schwarting, and Wöhr 2014). Categorisation thus depends on learning and exploiting the statistical structure of sensory inputs, an ability known as statistical learning that allows organisms to internalise the probabilistic structure of the environment, forming priors that disambiguate uncertain inputs, anticipate upcoming events, and optimise behaviour in a changing world (Fiser et al. 2010; Aslin and Newport 2012; Schapiro and N. Turk-Browne 2015).

A central question is how this statistical knowledge is represented. Competing theories propose that learners either construct generative models of the sensory world, encoding the distribution of inputs within each category, or adopt discriminative mappings that directly link sensory evidence to category labels or actions (Hsu and T. E. Griffiths 2009; Love et al. 2015; Peters et al. 2024). These strategies can yield similar performance, yet rely on fundamentally different internal representations and entail distinct computational trade-offs. To illustrate, a generative learner who distinguishes cats from dogs would acquire knowledge about the sensory structure of each category; what features define cats, how variable they are, and how they differ from dogs. Such a learner could infer unobserved properties (“does this new animal have claws?”) or imagine new exemplars consistent with past experience. A discriminative learner, by contrast, would only learn how to separate the two categories; able to classify an animal as cat or dog, but unable to answer any other questions about either. Generative learning thus supports generalisation and transfer but requires maintaining richer internal structure; discriminative learning is simpler and efficient when only boundary information is needed (Ng and Jordan 2001; Bouchard and Triggs 2004). Despite extensive theoretical work, direct behavioural evidence distinguishing these algorithms has been lacking.

While normative theories describe what behaviour should be in order to maximise reward and reduce uncertainty (Green and Swets 1966; Simon 1986; Whiteley and Sahani 2008), they do not specify how that behaviour is achieved. Algorithmic theories address this gap by defining how internal representations are formed and updated (Marr 2010). Generative and discriminative learning provide two such algorithmic solutions to the same normative goal: both can approximate optimal behaviour but through fundamentally different computations. To dissociate these possibilities, we designed a task that fixes category boundaries, reinforcement contingencies, and decision rules while manipulating only the sensory distributions within each category. This manipulation isolates the contribution of sensory priors i.e beliefs about stimulus statistics, independent of decision priors such as choice biases (Akrami et al. 2018; Gao, Tortell, and McClelland 2011; Ashwood et al. 2022).

Decision priors are choice-level biases, such as a preference for particular actions that arise from asymmetric rewards or choice probabilities (Gao, Tortell, and McClelland 2011; Rorie et al. 2010; Hanks et al. 2011; Ashwood et al. 2022). Sensory priors, in contrast, reflect knowledge about the distribution of sensory inputs themselves, such as which tones are more likely to occur within a category (Sohn et al. 2019; Akrami et al. 2018; Körding and Wolpert 2004; L Griffiths, Kemp, and B Tenenbaum 2008). Both can bias behaviour (Gupta et al. 2024; Findling et al. 2025; Ashwood et al. 2022; Boboeva et al. 2022; Akrami et al. 2018; Fassihi et al. 2014; Ashourian and Loewenstein 2011), yet rely on fundamentally different representations: action mappings versus models of the sensory environment. Previous studies primarily manipulated decision priors by adjusting reward asymmetries (Gao, Tortell, and McClelland 2011; Rorie et al. 2010; Feng et al. 2009), decision boundaries (Zhong et al. 2019; Hachen et al. 2020; Sainburg et al. 2025), response mapping (Ghosh and Zador 2021), or block-wise choice biases (Findling et al. 2025; Ashwood et al. 2022; Noel et al. 2025), leaving the sensory structure fixed. As a result, history-dependent biases often correlated more strongly with past actions than past stimuli (Lau and Glimcher 2005; Bolkan et al. 2022; Findling et al. 2025), obscuring whether they arose from decision-level reinforcement or true sensory-statistical learning (Gupta et al. 2024; Abrahamyan et al. 2016; Busse et al. 2011).

By varying the statistical structure of sensory inputs while holding reinforcement and decision rules constant, our design uniquely separates sensory from decision-level learning. Using this framework, we compared humans, rats, and mice performing the same auditory categorisation task. Cross-species comparison offers a principled way to ask whether similar adaptive behaviour arises from shared computational principles or distinct algorithmic implementations (Fiser et al. 2010; Santolin and Saffran 2018). Despite differences in architecture and scale, all three species must extract and exploit statistical structure to optimise performance, providing a common ground for testing general principles of learning.

In our two-alternative forced-choice (2-AFC) sound categorisation task (Otazu et al. 2009; Jaramillo and Zador 2011; Raposo et al. 2012; Kelly and O’Connell 2013; Znamenskiy and Zador 2013), subjects judged the category of auditory stimuli drawn from two distributions separated by a fixed boundary. By fixing the stimulus range and category boundary but varying the underlying statistical distribution of stimuli within each category, we created distinct statistical contexts that selectively engaged sensory priors. This manipulation changed the sensory statistics without altering the decision rule or reinforcement structure, allowing us to isolate how learners internalise and exploit environmental regularities to bias decisions under uncertainty.

We show that, under a normative model, the optimal decision-maker should bias responses toward the more frequent category near the boundary, thereby improving reward under uncertainty (Whiteley and Sahani 2008; Rorie et al. 2010; Gao, Tortell, and McClelland 2011). All three species adopted such near-optimal, context-specific biases and flexibly adapted to uncued changes in stimulus statistics. Yet behind this shared outcome, individuals diverged in how they represented and updated information: some relied on distributional (generative) models, others on boundary-tracking (discriminative) strategies. Crucially, these differences were invisible in end-point performance: only the trajectory of learning and trial-to-trial choice updates exposed their divergence. This reveals a general behavioural diagnostic principle—distinct internal representations leave identifiable signatures in learning dynamics even when behaviour appears optimal. By linking normative goals to algorithmic implementation within a shared cross-species task, this framework exposes the diversity of computational strategies that support adaptive behaviour across brains.

## Results

### A comparative paradigm to study learning and utilising sensory statistics in humans, rats and mice

We designed an auditory 2-AFC categorisation task to probe how humans, rats and mice learn and use sensory statistics (Figure 1A). On each trial, subjects were required to categorise a brief auditory stimulus as falling below a boundary (category A) or above it (category B). Human participants categorised pure tones according to frequency by selecting A or B on a screen, and received visual feedback indicating correctness of the response (Figures 1A and S1C; see Methods). Rats and mice categorised white noise stimuli according to amplitude (but see Figure S2 for frequency version). Freely-moving rats responded by choosing between two water ports located on the left and right of an operant chamber, and head-fixed mice responded by licking a left or right spout (Figures 1A and S1A,B). Rewards were delivered for correct choices, from the port or spout corresponding to the correct category: lower-range sounds to port/spout A, upper-range sounds to port/spout B. The mapping between ports/spouts and category labels (A/B) was counterbalanced across rats and mice (see Methods). Incorrect responses in both rodent tasks were followed by a brief timeout (see Methods and Figure S1 for details of trial structure). The range of sound amplitudes or frequencies was specific to each species. For ease of comparison, we describe stimuli in common units (‘distance from boundary’). Here, log physical stimulus values are remapped to place the minimum at −1, the maximum at 1, and the boundary at 0 (see Methods).

**Figure 1.**
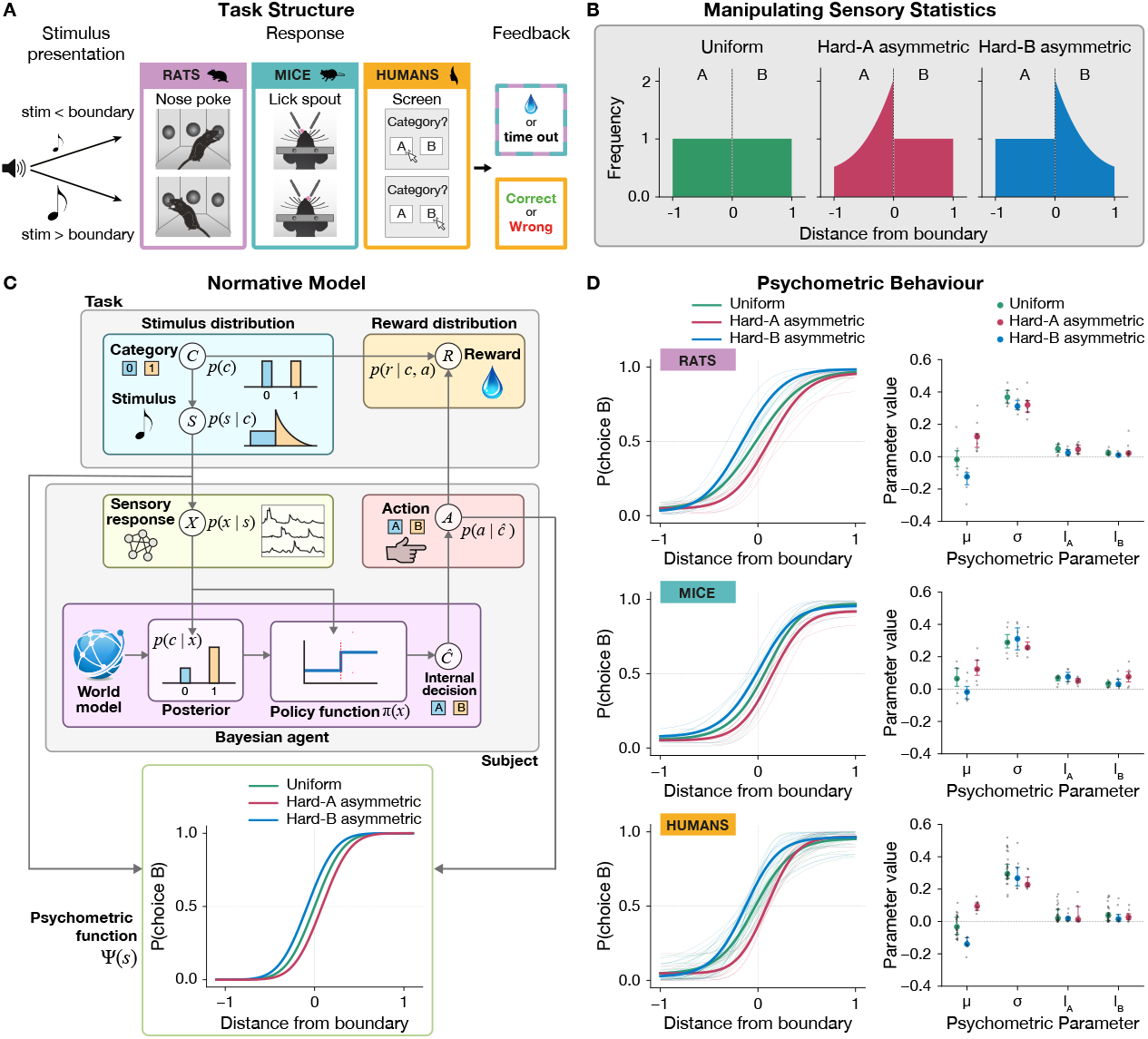
Cross-species sound categorisation paradigm for humans, rats, and mice. **A**: Experimental setup for each species. Rats were freely moving and indicated their choice by poking one of two nose ports (left/right). Mice were head-fixed and reported their choice by licking one of two water spouts. Humans performed experiments online and indicated their choice by clicking on a category button on the screen. Rodents received a water reward for correct choices and a timeout for errors; humans received visual feedback. **B**: Stimulus distributions used in the task. Three sampling distributions were tested: uniform, hard-A asymmetric, and hard-B asymmetric. In all cases, the total probability mass was balanced below and above the decision boundary, ensuring equal overall category probabilities. **C**: Schematic of the normative model. The model assumes noisy sensory encoding and maps posterior category estimates to decisions via a reward-maximising policy. **D**: Left panels show psychometric curves for all individuals (thin lines) and the average across individuals (thick line), separately for each species and stimulus distributions shown in (B). Right panels show individual parameter estimates (dots) from psychometric fits—mean (*µ*), width (*σ*), and lapse rates (*l*_*A*_ and *l*_*B*_)—alongside the group median and interquartile range (rats: n = 14; humans: n = 29; mice: n = 7).

To test whether the underlying sensory statistics influence categorisation behaviour, we exposed subjects to one of three stimulus distributions: uniform, hard-A asymmetric, or hard-B asymmetric (Figure 1B). In the uniform condition, stimuli were evenly sampled across the entire stimulus range (Figure 1B, green). In the hard-A condition, category A stimuli were over-represented near the category boundary, making them harder to classify, while category B stimuli were uniformly distributed (red). Conversely, in the hard-B condition, category B stimuli were concentrated near the boundary, and category A stimuli were sampled uniformly (blue). Importantly, although the shape of stimulus distributions varied, the category boundary was always fixed at the midpoint of the stimulus range, and the overall probabilities of drawing a category A or B stimulus remained equal (*P*(*A*) = *P*(*B*) = 0.5). This design isolates the effect of sensory statistics while keeping reinforcement contingencies and overall category probabilities constant. This allowed us to disentangle sensory statistical learning from action history or decision prior effects (Lau and Glimcher 2005; Bolkan et al. 2022; Abrahamyan et al. 2016; Busse et al. 2011), providing a unique behavioural assay of the internal representations that support statistical learning.

### A normative model of categorisation under uncertainty

We first asked: why should a decision maker care about the stimulus distribution, and how should decisions reflect it? To address these questions, we built a normative model (Figure 1C) that produces optimal behaviour. It describes an ideal decision maker with knowledge of the task structure and statistics, but constrained by sensory and decision noise. In this model, a Bayesian agent perceives a noisy sensory representation of the stimulus, and combines it with prior knowledge of the stimulus, category, and sensory noise distributions to compute the posterior probability of each category. Posterior probabilities are combined with knowledge about reward structure to produce a choice that maximises expected reward, despite possible corruption by downstream decision noise (see Normative Model). From an external experimenter’s perspective, these internal computations produce behaviour summarised by a psychometric function linking the presented stimulus to the probability of the observed categorical response.

The normative model shows that behaviour should adapt to the stimulus distribution because doing so increases reward. The strategy it employs compensates for sensory noise, reducing errors on difficult stimuli. More specifically, under the reward structure of our task, expected reward is maximised by choosing the category with greater posterior probability, given the sensory response. In our task where categories have equal probability of occurring, this corresponds to choosing the category under which the sensory response is most probable. A noisy sensory response near the boundary is more likely to have arisen from the category where stimuli are more concentrated in this region. Therefore, the normative model qualitatively predicts that when the stimulus distributions are asymmetric (hard-A and hard-B asymmetric distributions, Figure 1B, blue and red), choices in our task should be biased toward the category with greater density near the boundary. The predicted behavioural signature is a systematic shift in psychometric midpoint (Figure 1C; which shows psychometrics in the regime of stimulus-independent sensory noise and zero decision noise).

Consistent with this, psychometric fits to humans, rats and mice revealed similar effects (Figure 1D). We fitted psychometric functions, characterised by four parameters (see Psychometric Curve Fitting): 1. The mean, which reflects biases in the point of subjective equality between the two categories, 2. The width (slope) which captures perceptual sensitivity, 3. and two lapse values, which quantifies the error rate at the easiest category A and B stimuli. Across all species, the primary effect of changing the stimulus distribution was a change in the mean of the psychometric function Figure 1D; Kruskal–Wallis test for humans and rats where each individual was tested on only one distribution; humans: p-value < 0.001; rats: p-value < 0.001; Friedman test for mice where each individual was tested on all conditions: p-value < 0.01; see Statistical Analysis). Specifically, the psychometric curves for hard-A and hard-B were shifted rightward and leftward, respectively, compared to the uniform, creating significantly different means between the two conditions (in humans and rats, each subject was exposed to only one distribution; Mann–Whitney *U* test for independent groups, Bonferroni corrected; humans: p-value = 0.0001 for uniform vs hard-A, p-value = 0.0002 for uniform vs hard B, p-value = 0.0012 for hard-A vs hard-B; rats: p-value = 0.0340 for uniform vs hard-A, p-value = 0.0244 for uniform vs hard B, p-value = 0.0041 for hard-A vs hard-B; in mice group, the same subjects were tested across multiple distributions, so the groups were paired; Wilcoxon signed-rank test, Bonferroni corrected for mice: p-value = 0.0469 for hard-A vs hard-B, p-value = 0.0469 for hard-A vs hard-B; see Methods). These shifts indicate a systematic bias in choice behaviour toward the more frequent category near the boundary, as predicted by the normative model (Figure 1C). These shifts confirm that subjects adjusted their decisions in line with normative prediction.

#### Humans, rats, and mice exhibit near-optimal adaptation to sensory statistics

The normative model specifies how an ideal Bayesian agent should behave given two factors: the environmental structure and statistics (including the prior sensory and reward distributions) and the agent’s internal sensory and decision noise. Because all subjects experienced the same environment, any differences in predicted psychometric shifts arise solely from individual noise characteristics, assuming that each has learned the correct priors, as expected for an optimal Bayesian agent. Estimating this noise is therefore critical to test whether behaviour matches normative predictions.

We fit each subject’s performance in the uniform and one asymmetric distribution to estimate sensory and decision noise parameters (Figure 2A; see Fitting Noise Distributions). Stimuli *S* were modelled as noisy percepts *X* = *S* + *ε*, with *ε* drawn from a flexible distribution that captured both constant and stimulus-dependent variability. These fits yielded an individualised likelihood function for sensory perception. Combining this likelihood with the known prior then allowed us to simulate how an optimal Bayesian agent with the same internal noise would behave under the other asymmetric distribution. For example, parameters estimated from hard-A (plus uniform, if available) were used to predict performance in hard-B, and vice versa. For this analysis, all species were tested within-subject across both asymmetric distributions, with switches occurring uncued.

**Figure 2.**
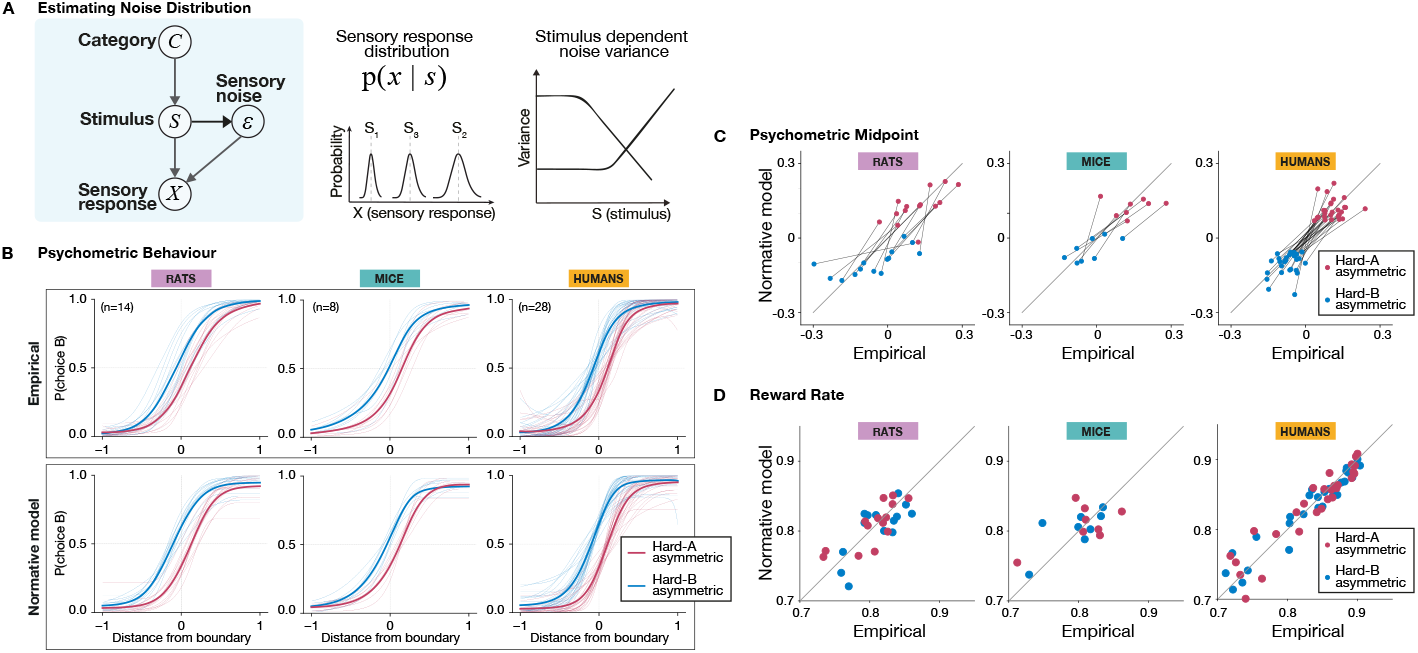
Humans, rats and mice adapt to sensory statistics as predicted by an optimal agent. **A**: Schematic of the internal sensory encoding model. Each stimulus *S* is transformed into an internal variable *X* by adding Gaussian noise: *X* = *S* + *ε*, where *ε* | *S* ∼ 𝒩 (0, *σ* ^2^(*S*)). The noise variance *σ* ^2^(*S*) is modelled with a non-linear stimulus dependent function, allowing for flexible encoding precision across the stimulus range (showing two possible examples in the right panel). **B**: Top panels show empirical psychometric curves for rats, mice, and humans under each stimulus distribution (hard-A in red and hard-B in blue, see Spline psychometric). Thick lines show the average across individuals; thin lines show individual subject fits. Columns correspond to species. Bottom panels show normative model predictions for each species, using subject-specific noise parameters estimated from uniform condition data. The model captures the direction and magnitude of the psychometric shift across contexts. **C**: Comparison of psychometric midpoints predicted by the normative model (y-axis) and observed empirically (x-axis), for each species. **D**: Comparison of reward rate predicted by the normative model (y-axis) and observed empirically (x-axis), for each species. The close alignment to the identity line in **C-D** indicates that subjects adjusted their behaviour in line with model predictions.

The model accurately predicted both the direction and magnitude of psychometric shifts, as well as overall reward rates (Figure 2B–D). Slopes of the normative–empirical midpoint relationship were reliably positive across individuals, with 95% BCa (bias-corrected and accelerated) bootstrapped confidence intervals that did not include native values (rats: [1.22 3.27], mice: [0.87 1.42], humans: [1.15 1.99]). By contrast, differences between empirical and normative reward rates had 95% BCa bootstrapped confidence intervals that included zero (rats: [-0.005 0.002], mice: [-0.006 0.016], humans: [-0.004 0.003]), showing that subjects across species adjusted their behaviour in a near-optimal, reward-maximising way.

#### Priors shape choices most when evidence is uncertain

Intuitively, priors should matter most when sensory evidence is weak. For stimuli far from the boundary, the evidence is clear, uncertainty is low, and choices are determined almost entirely by the stimulus itself. In these cases, changing the stimulus distribution should have little to no effect on behaviour. Near the boundary, however, the evidence is ambiguous, and here the balance of categories plays a decisive role: what matters is the relative category probability at the boundary, not the global shape of the distribution. Moreover, if sensory evidence becomes noisier, such as when stimulus duration is shortened, priors should carry more weight; if the evidence becomes more reliable, the influence of priors should fade.

Our experiments confirmed these predictions. When probability mass was shifted away from the boundary—so that local stimulus probabilities near the boundary remained close to uniform (‘off-boundary asymmetry’)—behaviour showed little or no measurable change in rats (Figure S3B). Easy-asymmetric contexts, which altered global distribution shape but preserved local imbalance near the boundary, produced biases equivalent to their “hard” counterparts in humans and rats (Figure S3A). Similarly, unimodal and bimodal manipulations that symmetrically over or under-sampled hard stimuli had no detectable effect in rats (Figure S3C).

Finally, in mice, reducing stimulus duration increased perceptual noise (as seen in shallower psychometric slopes; Figure S3E–F) and amplified the behavioural shift in hard-A and hard-B conditions, while longer durations reduced the effect (Figure S3D–F).

Together, these results suggest that priors are always incorporated, but their detectable impact is confined to conditions of high perceptual uncertainty, whether due to stimulus position near the boundary or to increased sensory noise.

### Different computational routes to the same optimal behaviour

Our normative model specifies the optimal bias for maximising reward under each stimulus distribution, but does not explain how this behaviour is learned. Importantly, from a normative perspective, generative and discriminative strategies are indistinguishable, because in both cases behaviour is ultimately determined by the posterior probability of category given stimulus, *P*(*c* | *s*). This posterior can be derived from full generative knowledge of the environment, i.e. the joint distribution *P*(*c, s*), or obtained directly through a discriminative mapping. In either case, the same posterior governs choice. To uncover which representations animals actually use, we introduced two complementary learning models: a boundary-estimation (BE) model, which tracks the decision boundary directly, and a stimulus-category (SC) model, which builds distributional representations of each category. Although humans and rodents converged on similarly near-optimal psychometric performance, such outcomes can arise from fundamentally different learning strategies, captured by these two models. Each model embodies a distinct hypothesis about the type of information subjects learn and use when performing the task.

In the BE model, the agent maintains a probability distribution over boundary location, initially uniform over the stimulus range (Figure 3A). Choices are made by comparing the noisy perceived stimulus (see Learning models) to this distribution: the probability of choosing category B is given by the fraction of the boundary distribution lying below the perceived stimulus. Feedback updates the boundary distribution, softened by parameters controlling update precision, learning rate, and a forgetting term that relaxes beliefs toward uniformity i.e. maximum uncertainty about the boundary location (see Boundary-Estimation model).

**Figure 3.**
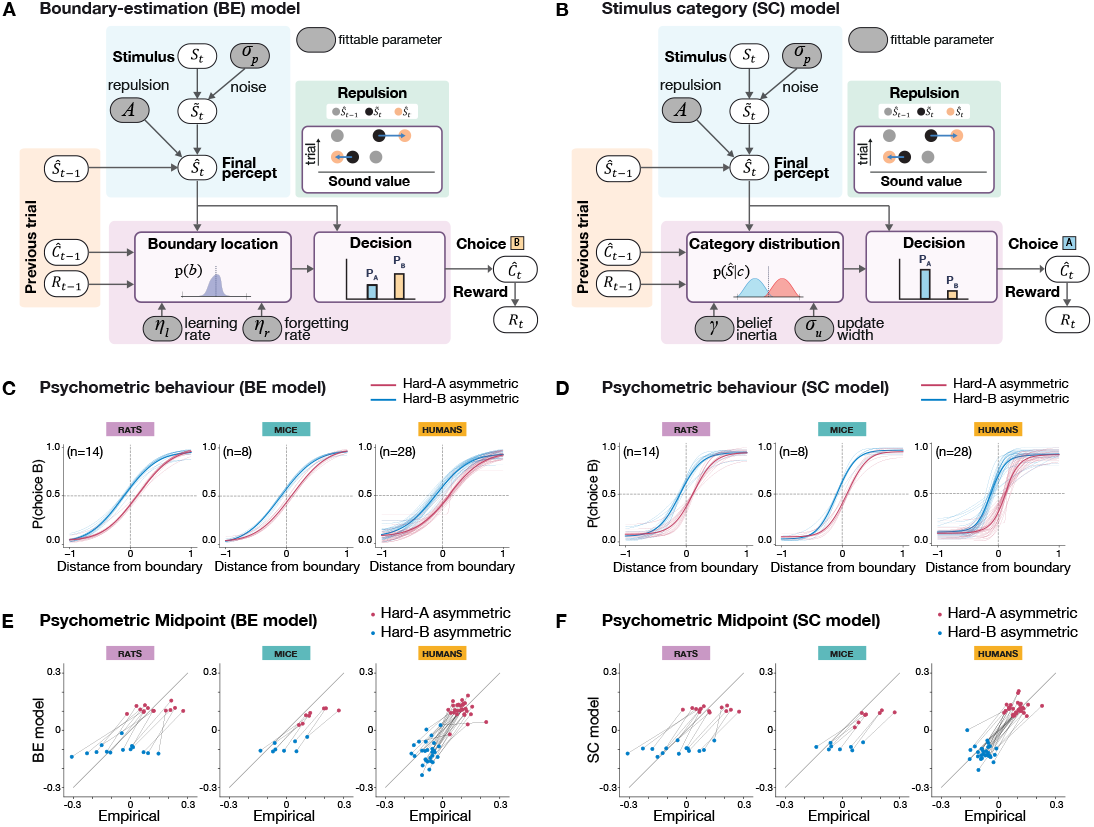
Boundary-estimation and Stimulus-category models account for learning of the optimal behaviour. **A**: Schematic of the Boundary-estimation (BE) model. A noisy perceived stimulus is compared to the agent’s belief about boundary location to generate a choice (below vs. above boundary). Following feedback, together with information about the previous trial (stimulus and reward), the belief over the boundary location is updated. **B**: Schematic of the Stimulus-category (SC) model. As in the BE model, choices are based on the perceived stimulus, but here decisions depend on beliefs about full category distributions, *P*(*S*|*C*). Feedback and trial history are used to update these category beliefs. **C**: Psychometric behaviour predicted by the BE model for hard-B contexts, simulated using parameters fit to hard-A (and vice versa), shown separately for rats, mice, and humans. **D**: Equivalent predictions for the SC model. **E**: Comparison of model-predicted and empirical psychometric midpoints (Point of Subjective Equality, see Methods) for BE across hard-A and hard-B conditions, shown for all individuals. **F**: Same as in (E) for the SC model. Together **C-F** show that both models were able to successfully reproduce the psychometric shifts observed, empirically, in response to changes in stimulus statistics.

In the SC model, the agent instead learns two category-specific stimulus distributions, *P*(*s*|*A*) and *P*(*s*|*B*), representing sensory priors for each category (Figure 3B). The probability of choosing a category is proportional to the amplitude of its distribution at the perceived stimulus. Following feedback, the chosen category distribution is updated locally around the perceived stimulus, while the opposite distribution remains unchanged (see Stimulus-category model).

We fitted both models to conditional psychometric performance (conditioned on the previous-trial stimulus, see Model Fitting) using data from the hard-A (and uniform, when available) distributions, and predicted psychometric behaviour under the hard-B distribution (and vice versa). Both models reproduced the observed psychometric shifts (as in Figure 3C–D): predicted midpoints (Point of Subjective Equality, PSEs) for hard-A were significantly larger than for hard-B (Figure 3E–F), consistent with empirical data. Across individuals, the slope of the relationship between BE-(or SC-) predicted and empirical midpoint values was reliably positive, with 95% BCa (bias-corrected and accelerated) bootstrapped confidence intervals that excluded zero (BE: rats: [1.14 2.79], mice: [0.87 1.26], humans: [1.11 1.51]; SC: rats: [1.13 2.29], mice: [0.84 1.19], humans: [1.24 1.60]). This confirms that both models captured systematic, context-dependent shifts in boundary placement across species.

### BE and SC models predict divergent updating strategies for humans and rodents

While both models captured the distribution-dependent psychometric shift, they rely on fundamentally different internal representations. The boundary-estimation (BE) model learns a belief over boundary location, whereas the stimulus-category (SC) model learns the shape of each category’s stimulus distribution. Therefore, the *trial-to-trial learning rules* differ in these two models. In BE, each trial’s perceived stimulus and feedback update the posterior over boundary location. Easy trials (far from the boundary) induce minimal updates; difficult trials (near the boundary) induce larger shifts, strongly influencing subsequent choice probabilities (Figure 4A, left). In SC, the agent instead updates the full category distribution locally around the perceived stimulus. Because mass is conserved, adding probability near one stimulus reduces it elsewhere. Thus, an easy-A trial (far from the boundary) reduces mass near the boundary, making subsequent difficult A trials less likely to be categorised as A; conversely, a difficult-A trial adds mass near the boundary, making subsequent difficult A stimuli more likely to be categorised as A (Figure 4A, right). BE and SC therefore make opposite predictions about how easy vs. difficult trials influence subsequent boundary-adjacent choices.

**Figure 4.**
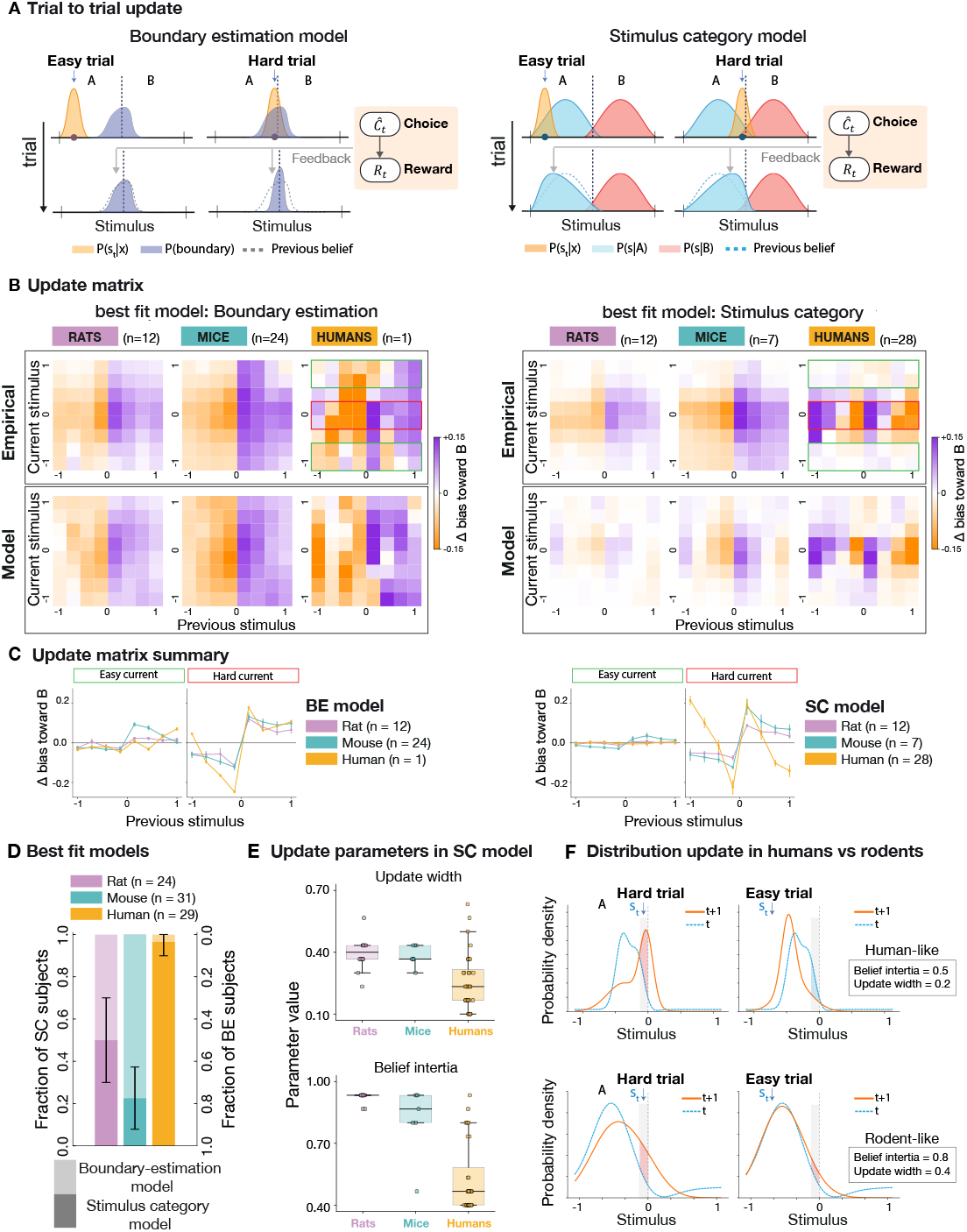
Boundary-estimation and stimulus-category models predict distinct trial-to-trial updating patterns. **A**: Schematic of trial-to-trial updates in stimulus-category (SC, left) and boundary-estimation (BE, right) model. In SC, an easy A trial (far from the boundary) increases probability mass near that stimulus, but reduces mass elsewhere (particularly near the boundary), thereby decreasing the likelihood of classifying the next hard A as A. Conversely, a difficult A trial (near the boundary) increases mass near the boundary, raising the likelihood of subsequent difficult A trials being categorised as A. In BE, easy trials induce minimal updates, whereas difficult trials produce large shifts in the boundary estimate, strongly biasing the next choice (shown here with weak forgetting). **B**: Average update matrices for each species in which the BE (right) or SC model provided the best fit. Top: empirical data; bottom: model predictions. **C**: Summary of the empirical update matrices for individuals best fit by SC (left) or BE (right). For each subject, the update matrix was averaged over easy current trials (green box in B) and hard current trials (red box in B). Similar to **B**, first the best fit model for each individual is identified, then the update matrix was averaged around current easy vs hard trials. Lines show the Δ bias toward category B as a function of previous stimulus, for rats (purple), mice (cyan), and humans (yellow); errorbars show SEM. For SC, previous-trial effects vanish on easy current trials across species, but diverge sharply on hard trials: in humans, the bias depends on both the value and category of the previous stimulus. After an easy A trial, the bias shifts away from A, whereas after a hard A trial it shifts toward repeating A—producing opposite-sign effects. Rodents instead show a simple win–stay pattern, increasing the probability of repeating the previous choice regardless of difficulty. BE agents show qualitatively different update signatures without these sign reversals. **D**: Fraction of individuals in each species best fit by SC or BE. Error bars: 95% binomial proportion confidence intervals for the proportion of individuals in which each model provided the best fit. **E**: Parameter estimates for individuals best fit by SC. Rats and mice had broader update widths (less precise updates) and higher belief inertia (slower updating) than humans. Thus, humans updated their category beliefs faster and more locally. **F**: Example post-correct updates of category A distributions in the SC model using human-like parameters (top; narrow update width = 0.2, low inertia = 0.5) or rodent-like parameters (bottom; broad width = 0.4, high inertia = 0.8). With human-like parameters, hard vs. easy trials have opposite effects near the boundary (hard trials increase, easy trials decrease mass). With rodent-like parameters, both easy and hard trials increase mass near the boundary. These results are consistent with weaker stripe structure in rodents update matrices.

Because these two models differ in their internal representations (boundary vs. category distributions), the outcome of model comparison is directly informative about what subjects have learned. To adjudicate between these alternatives, we fitted both models to conditional psychometric behaviour (post-correct; conditioned on the previous-trial stimulus), and performed an individual-level model comparison using repeated cross-validation with prediction on held-out data (see Model Fitting; see Figure S13 for model validation). Each subject was assigned to the winning model (BE or SC), and their predicted update matrices were generated from simulations with best-fitting parameters. The update matrix, defined as the difference between the post-correct conditional psychometric function (conditioned on the previous stimulus) and the overall post-correct function (Lak et al. 2020), provides a first-order summary of how the previous stimulus influences current-trial performance (see Figure S6 for empirical update matrices on different distributions).

Group-level averages of update matrices revealed species-specific signatures (see Figures S10 and S11) for individual update matrices). BE agents reproduced the canonical win–stay signature: previous correct choices, especially near the boundary, increased same-category responding (Figures 4B, left; 4C, left; S7 for hard-A vs hard-B distributions). SC agents, in contrast, generated a distinctive “stripe” pattern, reflecting fine-grained, stimulus-dependent updates across trials. This pattern was prominent in humans (Figures 4B, right; 4C, right; S7 for hard-A vs hard-B distributions), whose fitted parameters showed faster belief updating (lower inertia *γ*) and more localised updates (smaller *σ*_update_), relative to rodents (Figure 4E). In rodents, SC fits showed weaker and more diffuse stripe structure, consistent with greater belief inertia and broader updating kernels (Figure 4E). This is further illustrated by simulations for category A updates, based on the human-like vs rodent-like parameters (Figure 4F). Narrow and large updates in humans result in opposite effects of hard vs easy on stimuli close to the boundary (Figure 4F, top), in contrast to broader and similar updates in rodents (Figure 4F, bottom). These differences explain why human update matrices exhibit sharp, detailed stripe patterns, whereas rodents—even when best fit by SC—show weaker, more diffuse updates (see Figure S11 for individual update matrices).

A further general property of the SC model was evident across all species: when the current trial was easy (far from the boundary), the influence of the previous trial was minimal. This follows because the alternative category carries near-zero probability mass at easy stimuli, making choices invariant to prior redistributions of probability (Figure 4C, left). In rodents fit by the BE model, the inferred boundary distribution retained substantial uncertainty even for easy stimuli i.e. the tails of the distribution at easy A and easy B remained significantly above zero (Figure 4C, right). This boundary uncertainty explains why rodent BE subjects were more influenced by trial history even for easy stimuli, and why their psychometric curves were shallower compared to SC subjects (see Figure S9).

Overall, the winning models’ update matrices closely reproduced the empirical ones (Figures 4B, S10 and S11), whereas the losing models diverged Figure S5). Importantly, while the update matrix captures only one-trial-back history effects and does not reflect deeper temporal dependencies, it reveals clear mechanistic differences between the models. Moreover, while BE appears similar to a win–stay heuristic at this level, a pure win–stay rule cannot explain the systematic hard-A vs. hard-B biases we observed.

In summary, humans predominantly conformed to the SC model, maintaining rich generative representations of category distributions that produced fine-grained, stimulus-specific updating. Rodents, and especially mice, were better described by the BE model, reflecting simpler boundary-tracking strategies. Rats exhibited greater inter-individual diversity, with a substantial fraction fitted best by the SC model, albeit with weaker and less detailed stripe patterns than in humans (Figures 4B and S11). These results demonstrate that while both BE and SC models achieve near-optimal adaptation, they do so through distinct internal representations and trial-level dynamics, with species differences in the precision and granularity of learning.

### Humans, rats and mice have similar convergence of their psychometric behaviour toward optimal

Earlier, we showed that humans, rats and mice flexibly switch their psychometric behaviour when the underlying statistical context changes without explicit cues. But how quickly do they achieve this shift? In the human experiment, participants completed 24 blocks of 50 trials, with each block randomly assigned either a hard-A or hard-B asymmetric stimulus distribution (Figure 5A). This random switching prevented participants from inferring any higher-level task structure and forced them to adapt based solely on accumulated sensory evidence. Fitting block-specific psychometric curves, focusing on blocks that followed a distribution switch (i.e., hard-A preceded by hard-B and vice versa), revealed consistent, distribution-dependent biases—mirroring those observed in fixed-distribution sessions (Figure 1D), demonstrating humans’ ability to rapidly and reversibly adapt to changes in sensory statistics (Figure 5B, two-sided paired t-test; n=28, p-value < 0.000001 for hard-A vs hard-B).

**Figure 5.**
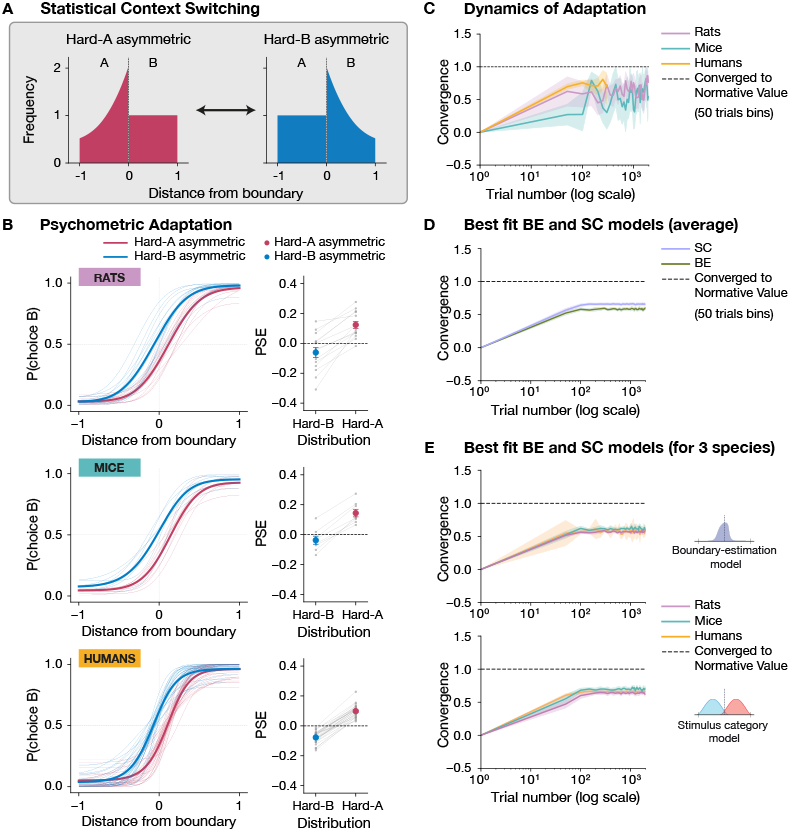
Similar convergence toward the optimal behaviour. **A**: Schematic of distribution switches between hard-A and hard-B asymmetric contexts. **B**: Psychometric functions (4-parameter cumulative Gaussian fits) for rats, mice, and humans following a switch. Left panels show psychometric curves for all individuals (thin lines) and the average across individuals (thick line), separately for each species. Right panels show paired Point of Subjective Equality (PSE) values, extracted from psychometric fits, for each individual in hard-A vs. hard-B conditions (circles show mean *±* SEM, gray lines connect within-subject data). **C**: Time course of convergence toward normative predictions across species, quantified using PSE values estimated from psychometric fits in contiguous 50-trial bins. Solid line: across-subject mean per bin. Shaded region: pointwise bootstrap confidence intervals. **D**: Convergence of simulated BE and SC agents using best-fit parameters from each subject’s winning model under the same switch conditions. **E**: Convergence trajectories for simulated BE (top) and SC (bottom) agents, shown separately for each species.

Switching experiments in rats and mice were analogous. For both species, switching the stimulus distribution produced a reversal in psychometric bias: exposure to hard-A produced a bias toward category A near the boundary, while hard-B induced bias toward category B, independent of training history (two-sided paired t-test; rats (n=14): p-value = 0.000003 for hard-A vs hard-B; mice (n=8): p-value = 0.000006 for hard-A vs hard-B). Thus, rodents also flexibly adapted to the prevailing stimulus statistics (Figure 5B). In a within-session variant with probabilistic switches (similar to humans) from hard-A to hard-B (or vice versa) every 100 trials, rats likewise adapted flexibly to the underlying statistical context (Figure S4).

To quantify the time course of adaptation, we measured the convergence of the psychometric function toward the optimal performance predicted by our normative model discussed earlier (see Methods). Specifically, we computed the Point of Subjective Equality (PSE) in 50-trial windows after each switch and normalised these PSEs to the mean PSE before the switch:

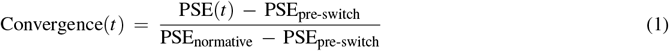

Across species, near-optimal behaviour, as expressed by the plateau in convergence, emerged within 200 trials (Figure 5C), with humans reaching higher convergence than mice by trial 50 post-switch (Mann–Whitney *U* test, Bonferroni corrected; rats vs. mice: p-value = 0.2137; mice vs. humans: p-value = 0.0008; rats vs. humans: p-value = 1.0000; rats: n=14; mice: n=6; humans n=28).

By trial 50 post-switch, humans had already reached stable convergence values that did not differ from later performance (Wilcoxon signed-rank, convergence at 50 trials post-switch vs mean of subsequent trials, p-value = 0.246, n=28). Rats likewise showed stable convergence after 50 trials (Wilcoxon signed-rank, p-value = 1.000, n=14). In contrast, mice showed lower convergence values at trial 50 than across subsequent trials (Wilcoxon signed-rank, p-value = p=0.031, n=6). By 200 trials post-switch, however, convergence in mice no longer differed from subsequent trial windows (Wilcoxon signed-rank, p=0.5625, n=6), indicating that adaptation had stabilised.

This convergence of psychometric performance toward the normative prediction maximises reward (see Figure S12). Consistent with this, accuracy (equivalent to reward rate) dipped immediately after the switch but recovered as convergence increased (Figure S12). This pattern was visible in all species and was particularly pronounced in mice, whose slower updating produced a more marked initial drop before gradual improvement.

A key question is whether the nature of the learned representation—category distributions or a decision boundary—affects how quickly behaviour convergences to the normative solution. In principle, the two models can adapt at different rates depending on their parameter regimes. Yet, when we simulated choices from each subject’s best-fit model under switch conditions matched to the experiments, both SC and BE models reached the normative solution with comparable speed (Figure 5D-E), under switching conditions matched to the experiments. Thus, despite species differences in update dynamics (e.g. richer, stimulus-dependent SC updates in humans, and some rodents, versus simpler boundary-based updates in others) and despite the contrasting representations assumed by the two models (generative versus discriminative), both pathways mapped onto final choice behaviour that converged toward the normative solution with similar efficiency. This convergence is not trivial: it reflects that, in the regimes fitted to behaviour, both SC and BE strategies are capable of supporting near-optimal adaptation.

Taken together, these findings show that humans, rats and mice share the ability to flexibly adapt to changes in sensory statistics, and that this flexibility can emerge from distinct underlying computational strategies.

## Discussion

### Distinct learning algorithms can yield shared optimal behaviour

Animals often behave as if they exploit environmental regularities optimally (Fiser et al. 2010), but optimal outcomes do not uniquely determine the computations that generate them. To disentangle normative performance from algorithmic process, we developed a cross-species paradigm that holds category boundaries, reinforcement contingencies, and decision rules fixed while manipulating only the statistical distribution of sensory inputs within each category. This design isolates how learners represent and update sensory structure, independent of reward or explicit rule changes, a distinction that previous categorisation studies could not make.

Using this approach, we show that humans, rats, and mice all adapt their decisions to changes in sensory statistics in a near-normative way i.e. biasing choices toward the more frequent category near the decision boundary where uncertainty is greatest (Whiteley and Sahani 2008). Yet behind this shared behavioural solution lie fundamentally different learning algorithms. Generative learners maintain and update models of the sensory distributions defining each category, whereas discriminative learners adjust only the decision boundary mapping stimuli to choices. Both achieve comparable performance but rely on different information and learning rules.

Crucially, these algorithmic differences were invisible in end-point psychometric performance: only the temporal structure of learning i.e. the trial-to-trial updates following feedback, revealed the underlying computation. This behavioural diagnostic provides a new, falsifiable tool for inferring how different brains represent and update sensory regularities, bridging normative theory and algorithmic implementation.

### Adaptive integration of sensory and reward information

Learning the statistics of the environment supports adaptive behaviour in uncertain and changing contexts. Using our two-alternative forced-choice (2-AFC) sound categorisation task with precisely controlled stimulus distributions, we asked how humans, rats, and mice adapt when the sensory structure of the task changes but the decision rule and reward contingencies remain constant. Previous paradigms typically manipulated rewards, boundaries, or block-wise choice biases to probe decision priors (Gao, Tortell, and McClelland 2011; Rorie et al. 2010; Feng et al. 2009; Findling et al. 2025; Ashwood et al. 2022), making it impossible to separate sensory from action-level biases. By decoupling the reinforcement structure from the stimulus statistics, our design provides a uniquely clean assay of how sensory regularities themselves shape learning and choice.

Under perceptual noise (Neri 2010), the optimal decision-maker should bias responses toward the locally over-represented category near the boundary (Simon 1986; Whiteley and Sahani 2008), thereby improving expected reward without altering the decision rule. Consistent with this prediction, all species exhibited context-specific biases that enhanced performance and reward rate (Whiteley and Sahani 2008). Adaptation was selective and efficient—restricted to ambiguous, boundary-adjacent stimuli—and scaled with sensory reliability. Mice, for instance, relied more on priors when stimuli were shorter and noisier. In contrast, global changes in the overall distribution that did not affect local uncertainty had no measurable effect. Thus, animals exploited sensory statistics only when doing so improved expected reward, providing a clear behavioural signature of efficient statistical integration.

Although the normative observer captures this adaptive bias, it does not specify how sensory information is represented or updated. Our experimental design, by stimulus statistics from reinforcement contingencies, shows that similar normative outcomes can arise from distinct algorithmic routes, revealing a diversity of computational mechanisms underlying adaptive behaviour.

### Distinct algorithmic regimes revealed by learning dynamics

The trial-by-trial dynamics of learning revealed two distinct computational regimes. In generative learning, subjects maintain structured internal models of category distributions, capturing how stimuli are distributed within each class and how they might vary across contexts (Behbahani and Faisal 2012). Such learners can infer unobserved features and generalise to novel examples, reflecting a representation of the underlying sensory world itself. In discriminative learning, by contrast, subjects encode only the mapping from sensory evidence to category labels (Hsu and T. E. Griffiths 2009; Love et al. 2015; Ng and Jordan 2001), sufficient for accurate classification but blind to the structure of sensory space. These regimes embody complementary trade-offs: generative learning supports generalisation and transfer from limited data but requires richer internal structure, while discriminative learning is more efficient and robust under stable mappings (Bouchard and Triggs 2004).

We formalised these regimes with two computational models. The Stimulus–Category (SC) model represents the full distribution of stimuli within each category and updates these beliefs on each trial, renormalising the internal probability structure. The Boundary–Estimation (BE) model tracks only the categorical boundary and updates it from feedback, independent of within-category statistics (Jebara 2012). while both captured overall psychometric shifts, their trial-to-trial update dynamics diverged sharply.

SC learning redistributed probability mass within categories, producing graded, stimulus-specific “stripe-like” signatures in the update matrix, reflecting probability normalisation and the propagation of sensory and category information across trials. BE learning, by contrast, produced monotonic, win–stay–like updates reflecting categorical reinforcement. These distinctive temporal signatures provide a mechanistic diagnostic of the underlying algorithm. Even when augmented with sensory repulsion, BE could not reproduce the structured update patterns observed in human data, underscoring the mechanistic distinctiveness of SC.

Cross-validated model comparison confirmed systematic differences across species and individuals. Humans were best described by SC, with sharply localised updates and low belief inertia; rodents showed broader, slower updates, reflecting coarser internal representations. Rats spanned both regimes, whereas mice were predominantly BE-like. Thus, different learners—both across and within species—achieved similar normative outcomes through distinct representational strategies.

### Variability across species and individuals

The coexistence of multiple strategies across species and individuals suggests that the brain flexibly balances generative and discriminative computations depending on ecological constraints, task structure, and experience. Generative strategies may be favoured when sensory inputs are uncertain or rapidly changing, while discriminative shortcuts may suffice when sensory environment is stable. The broad distribution of strategies in rats, including evidence for strategy switching, indicates that individuals can dynamically recruit different algorithms depending on context or internal state.

Some apparent species differences may reflect procedural factors—such as freely moving versus head-fixed conditions, motor demands, or shaping histories that influence engagement or belief-updating timescales. Yet the convergence of near-optimal adaptation across distinct neural architectures is itself a striking result: different brains can reach equivalent behavioural solutions via distinct computational routes. Understanding how the brain selects, combines, or transitions between these algorithms remains a central challenge for future cross-species behavioural and neural studies.

### Neural implications

Identifying the learning algorithm is essential for interpreting neural data. Without knowing whether behaviour reflects a generative or discriminative computation, it is impossible to determine what any given brain area represents or how neural dynamics support learning. The same behavioural pattern can arise from entirely different circuit mechanisms—whether updating sensory representations, reweighting action mappings, or integrating feedback—and these distinctions fundamentally shape how we interpret neural function.

Generative (SC-like) learning implies adjustments to sensory representations, such as shifts in category-selective tuning in auditory cortex (Xin et al. 2019), reflecting reshaping of priors over sensory space. Discriminative (BE-like) learning, by contrast, may arise from reweighting of sensory–motor mappings in corticostriatal circuits (Cox and Witten 2019), biasing decision variables without altering sensory representations. Both can yield similar psychometric behaviour but rely on different loci and forms of plasticity.

Once the algorithmic regime is known, it defines specific neural hypotheses. For a generative learner, trial-to-trial fluctuations in neural activity can be regressed against model-inferred updates in sensory and category-level probabilities, revealing how top-down and bottom-up information jointly shape learning. SC-like behaviour likely requires coordination between auditory and association cortices and higher-order regions such as frontal or orbitofrontal cortex, which integrate feedback and maintain probabilistic beliefs. By contrast, BE-like learning should manifest primarily in decision or action circuits, such as striatum or premotor cortex, where reinforcement directly biases boundary estimates.

Recent large-scale recordings show that sequential choice biases in rodents often reflect action kernels, reinforcement-weighted tendencies to repeat previous actions, implemented in striatal networks (Findling et al. 2025). Our results extend this framework by identifying boundary estimation as a distinct discriminative mechanism and contrasting it with distributional learning, which possibly depends on cortical updating of sensory priors. These distinctions yield clear predictions: discriminative computations should localise to decision circuits, whereas generative ones require coordinated plasticity across sensory and associative cortices.

Taken together, these findings caution against assuming that behavioural biases reflect a single neural mechanism. The same pattern of history dependence could arise from action-level reinforcement signals (possibly in striatum Samejima et al. 2005; Daw et al. 2011; Greenstreet et al. 2025), from boundary-tracking computations in decision circuits (Freedman, Riesenhuber, et al. 2001; Freedman and Assad 2006; Reinert et al. 2021), or from distributional learning (possibly in sensory cortex Berkes et al. 2011; Walker et al. 2020). Our comparative framework makes these alternatives explicit and provides behavioural signatures that can guide circuit-level tests. Disentangling these mechanisms—by orthogonalising sensory and decision priors and tracking their neural correlates—will be critical for correctly interpreting the neural basis of statistical learning.

Finally, the diversity of strategies across species and individuals points to an adaptive computational repertoire rather than a single universal algorithm. Humans predominantly relied on generative learning, mice on discriminative, while rats exhibited intermediate or mixed strategies—sometimes switching between them. This variability suggests that different brains can flexibly recruit distinct algorithms depending on uncertainty, task demands, or internal state. Even when rodents employed generative strategies, their category representations were coarser than those of humans, sufficient for near-optimal behaviour in our task but potentially limiting in richer or dynamically changing environments. Understanding how these algorithms are selected, combined, and adapted—at both the behavioural and neural level—will be key for explaining how diverse brains achieve flexible intelligence.

## Methods

### Human subjects

The study was approved by the University College London (UCL, London, UK) Research Ethics Committee [16159/001]. Electronic consent was obtained in accordance with the approved procedures. In all experiments, participants (*n* = 98) were recruited using the Prolific online platform (www.prolific.co). The experiments were carried out online using Psychopy (Peirce et al. 2019) or Gorilla (www.gorilla.sc), using their APIs or custom code in JavaScript. Participants were paid a fixed amount for their time and a bonus payment based on their performance to motivate their continuous attention. Once a participant had completed an experiment, they were excluded from any further experiments, and only participants with a high completion rate on Prolific were allowed to start the experiment. The demographics of the participants were matched with those of Prolific’s available set of participants.

### Rat subjects

All experiments were conducted in accordance with UK Home Office regulations (Animal Welfare Act 2006) and approved by the local Animal Welfare and Ethical Review Body (AWERB). Adult male Lister Hooded rats (Charles River, UK; in total: *n* = 44; intensity categorisation: *n* = 41, frequency categorisation: *n* = 3) were used in all experiments; training and testing started at ≥ 8 weeks of age. Rats were housed in large clear plastic cages and maintained on a 12h reverse light/dark cycle in a temperature and humidity-controlled room. All training and testing took place during the dark phase. Six rats received Neuropixels implants and were singly-housed post-surgery to protect the implant; all other rats were group (2 or 3) housed. Food was available *ad libitum*, but animals were placed under water restriction to motivate performance in the behavioural task. Outside of daily training sessions (1.5–3 h), rats received 5–15 min *ad libitum* access to water. Body weights were monitored regularly and maintained at a minimum of 90% of pre-restriction baseline.

### Mouse subjects

All experimental procedures were conducted in accordance with the UK Animals (Scientific Procedures) Act 1986, under Home Office regulations, and approved by the Animal Welfare and Ethical Review Body (AWERB). Two mouse lines were used: CA/C57BL/6J and TL/VGAT-ChR2-EYFP. Only male mice were included (in total: *n* = 34; intensity categorisation: *n* = 31; frequency categorisation: *n* = 3). Animals arrived in the experimental holding room at 6 weeks of age and underwent a 5-day acclimation period, during which they were handled daily to habituate them to the experimenter. Mice were house under a 12-hour reverse light/dark cycle (dark-phase: 8am to 8pm, with one hour of “twilight” between 7am/pm and 8am/pm) throughout the study. All experiments were conducted during the “dark” phase. Unless otherwise stated, food (standard chow) and water were provided ad libitum. Environmental enrichment was provided to reduce boredom and aggression and to offer shelter. During behavioural testing, mice were placed under controlled water restriction to maintain motivation, receiving 1.25–1.5 mL water per day, either during task performance or as supplementary water from the experimenter. Body weight was monitored daily, and animals were maintained at no less than 80% of their baseline weight. If a mouse’s weight dropped below 80%, or if signs of dehydration or discomfort were observed, supplemental water was provided until weight and condition returned to acceptable levels.

#### Surgery

Mice were acclimated to the experimental unit for at least one week prior to surgery. Surgical procedures were never combined with other manipulations (e.g., water restriction). Animals were required to be in good health and weigh at least 18 g before surgery. All procedures were performed under aseptic conditions in accordance with the Project Licence and UK Home Office regulations.

Anaesthesia was induced with isoflurane (4%) and maintained at 1–2% in oxygen, enabling reliable monitoring of anaesthetic depth. Pre-operative analgesia (Metacam, 2.5 µL/kg, s.c.) was administered, followed by oral dosing post-operatively. Eyes were protected with ophthalmic ointment, and body temperature was maintained using a heat pad and monitored with a rectal probe (target: 36.5–37^°^C). Breathing rate was kept between 60–100 Hz throughout.

Mice were secured in a stereotaxic frame using ear bars. The scalp was shaved, cleaned with antibacterial solution, and incised to expose the skull, which was cleaned of blood. The skull was levelled so that the difference between bregma and lambda was less than 20 µm. A custom stainless-steel head-bar was positioned at AP –2.95 using a 3D-printed holder and affixed with dental cement, sealing the interface between exposed skull and skin.

Following surgery, animals were weighed and transferred to a heated recovery cage with access to food, water, and wet mash. Recovery was typically complete within 30 minutes. Mice were monitored daily for three days, with weight and signs of discomfort or distress recorded. Only after full recovery were further procedures (e.g., water restriction for behavioural training) initiated.

### Human behaviour

The human version of the two-alternative forced-choice (2-AFC) categorisation task was implemented online. On each trial, a fixation cross was presented at the centre of the screen for 500 ms, followed by a 300 ms auditory stimulus (pure tone, see Figure S1D). The choice options were displayed as two boxes labelled A and B on the screen. Participants responded by clicking on one of the boxes within a 5 s response window. Upon response (or after the time limit), visual feedback indicated whether the choice was correct or incorrect (Figure S1C). In an earlier version of the task (uniform distribution, n = 30) participants responded using the “s” or “k” keys to select option A or B, and advanced to the next trial, in a self-paced manner, by clicking a “Next trial” button. In all versions, low-frequency sounds were mapped to category A and high-frequency sounds to category B. The frequency range spanned 380–420 Hz and was split into two equal, non-overlapping sets, with the boundary set at 400 Hz (Figure S1D). To parallel the rodent experiments, participants were not given explicit instructions on the categorisation rule; instead, they were told to attend to the sound, make a choice and learn from the feedback. Participants discovered the task rule by trial-and-error and were asked post-hoc to describe their decision strategy in writing. The first 30 trials served as learning trials and were excluded from analysis. These trials were used to verify that participants had understood the task; only those passing this stage continued to the main task, comprising 1200 trials, organised into 24 blocks of 50 trials. Ten subjects in the uniform setting completed only 8 blocks (400 trials in total). Within each block, stimuli were drawn a fixed distribution. The next block was randomly assigned to a different distribution. Participants were included in the analysis if they achieved a minimum performance of 70% accuracy in their session.

### Rat behaviour

Rats performed a two-alternative forced-choice (2-AFC) sound-intensity categorisation task in custom-built behavioural boxes equipped with three nose-poke ports (left, centre, and right) arranged along a curved wall. Two speakers were mounted above the left and right side ports. Each port contained a visible white LED, and the side ports were equipped with sipper tubes for water delivery controlled by solenoid valves. The rigs were constructed from 3D-printed components and acrylic panels.

Behaviour was controlled and monitored in real-time using the Bpod system (Finite State Machine Gen 2, Sanworks) in combination with the BControl software interface (Sanders and Kepecs 2014; Brunton, Botvinick, and Brody 2013). BControl provided flexible protocol design, stimulus delivery, and timestamping of all events with *<* 1 ms accuracy.

The Bpod hardware interfaced with LEDs for light cues, infrared sensors for nose-poke detection, solenoid valves for water delivery, and an analogue output module for sound control. Sounds were played through custom-built speakers (Sonitron amplifier, piezoelectric miniature speaker; 86 dB, 1–20 kHz). Hardware details are documented in the Bpod Wiki.

#### Training protocol

Rats were water-restricted to motivate performance. Each daily training session typically for 90 min, occasionally 120–150 min, with rats completing 200–500 trials. Light in the centre port indicated that a trial could be self-initiated by nose-poking. Rats were pre-trained to habituate to the behavioural rig and get accustomed to water delivery in the rig from the nose ports.

Training progressed through multiple shaping stages:

#### 1. Stage 1: Habituation and reward association

Naive rats were first shaped on a classical conditioning paradigm, where they learned to poke in a lit side port to receive water, accompanied by a reward click and LED illumination.

#### 2. Stage 2: Trial initiation and fixation

Rats initiated trials by poking in the centre port when the LED was lit, and learned to maintain fixation until a go cue indicated the permitted withdrawal. A visual cue (LED light) in the correct side port guided the animal’s response. The fixation period was gradually increased from 10 ms to 2 s.

#### 3. Stage 3: Stimulus introduction

Auditory stimuli were introduced during fixation, but the correct side port was still lit after the go cue. Therefore, rats were not explicitly required to learn the association between stimulus and side of the response yet.

#### 4. Stage 4: Full task

No visual cue was provided after the go cue; rats were required to infer the correct side port from the auditory stimulus alone.

Rats typically reached criterion performance (70% accuracy) within 5-7 sessions.

#### Task structure

In the full task, rats fixated in the centre port until an auditory go cue (3 kHz pure tone, 100–200 ms) signalled the end of the fixation period. Three hundred ms after trial initiation, a broadband noise stimulus (2–20 kHz) was presented for 400 ms (in some sessions, 150–250 ms), followed by a 100–300 ms delay. Rats then indicated their choice by poking into one of the two side ports to obtain a water reward. Correct responses were signalled by LED illumination and water delivery (reward click sound of 1.5 s was used for the first cohort and later removed). Incorrect responses resulted in a 5 s timeout (Figure S1A). Early withdrawal (“violation” trials) triggered a 9 kHz pure tone of 0.5 s duration and a 1 s timeout, and were excluded from analysis.

#### Stimulus parameters and mapping

The auditory stimuli were broadband noise (2000-20,000 Hz) produced as varying sound pressure level (SPL) values sampled from a zero-mean normal distribution. The intensity of the stimuli varied between 66 and 78 dB (Figure S1D). Mean amplitude levels across experimental rigs ranged from 66.00 *±* 1.69 dB at the lower end to 77.57 *±* 3.03 dB at the upper end, with some variability due to differences in speaker hardware. The category boundary was set at the mean intensity level across rigs, corresponding to 70.50 *±* 2.25 dB. The task required rats to learn to categorise the intensity of the sounds into two arbitrary categories. The decision boundary separating the two categories was set to be the mean of the overall range of sound intensities. For a subset of rats, if the sound intensity was higher than the decision boundary, the correct action was to poke into the left-hand nose port, whereas if the intensity was lower than the decision boundary, the correct action was to poke into the right-hand port. A separate cohort of rats was trained to use the opposite rule to map stimulus intensity and choice of side-port for counterbalancing purposes (high intensity sounds associated with poking on the right port and low intensity sounds with poking on the left port).

For the experiments described in Figure 1, rats performed an auditory task with stimuli drawn from one of the three statistical contexts. Some animals had prior exposure to other paradigms; accordingly, the figure shows data from the first of the three statistical contexts each animal experienced. For the experiments described in Figure 5 rats were first exposed to one of the two statistical context and upon reaching stable performance, the sampling distribution was switched once to the distribution with opposite asymmetry. For the experiments described in Figure S5A, rats were trained from naive status on auditory stimuli sampled from one of the two on-boundary asymmetric distributions. For the experiments described in Figure S3B, task-expert rats were exposed to the two off-boundary asymmetric distributions in separate sessions, one distribution per session. For the experiments described in Figure S3C, task-expert rats were exposed, in separate sessions, to one of the three statistical distributions (one distribution per session).

Sessions were included in the analysis if rats achieved ≥ 70% accuracy on valid trials (non violation).

### Statistical context manipulation

Stimulus values were normalised to the range [−1, 1], with 0 marking the decision boundary between categories A (negative values) and B (positive values). Across all statistical contexts, categories A and B were presented with equal overall probability (*p* = 0.5), but the distribution of stimuli within each category varied according to the experimental condition.

### Non-symmetric distributions

#### Asymmetric distributions in rats

For asymmetric contexts, one category was uniformly distributed while the other was over-represented either near the decision boundary (“hard” stimuli) or far from it (“easy” stimuli). In the hard-A context, category A stimuli [−1, 0) were sampled from a distribution skewed towards the boundary at 0, whereas category B stimuli (0, 1] were sampled uniformly. The hard-B context was defined analogously, with the category roles reversed. Stimulus sampling was performed in logarithmic space using MATLAB; exact code is provided in the accompanying repository.

#### Asymmetric distributions in humans

Human stimulus sampling followed the same logic but was implemented using a predefined lookup table of MP3 files for online presentation. Asymmetric contexts were generated from truncated exponential distributions. In the hard-A context, category A stimuli ([−1, 0)) were drawn from a truncated exponential skewed toward the boundary, while category B stimuli ((0, 1]) were drawn uniformly. The hard-B context was defined analogously, with the roles of the categories reversed.

#### Easy-asymmetric manipulations

In both species, we tested whether psychometric biases arose from the relative representation of categories near the boundary, or from the exact shape of the asymmetric distribution. To do this, we introduced easy-A and easy-B contexts, in which the non-uniform category over-represented easy stimuli (far from the boundary) and under-represented hard stimuli (near the boundary). For the experiments in rats, sampling was implemented in logarithmic space using MATLAB to generate probability weights that increased with distance from the boundary on the manipulated side. In these manipulations the paradigm used discrete stimulus values (*n* = 8 and the probability mass functions for the easy-A and easy-B contexts were:

- Easy-A: prob = [0.2770, 0.1749, 0.0438, 0.0043, 0.1250, 0.1250, 0.1250, 0.1250]
- Easy-B: prob = [0.1250, 0.1250, 0.1250, 0.1250, 0.0043, 0.0438, 0.1749, 0.2770]

All other task parameters and procedures were identical to those described for the main experimental conditions.

#### Off-boundary asymmetric manipulation

In a subset of sessions, we tested whether rats would develop a bias when the asymmetric sampling of stimuli occurred away from the decision boundary. In these “off-boundary” contexts, one category was sampled more densely at stimulus values far from the boundary (i.e., in the mid to easy range), while the other category was sampled uniformly across its range. This manipulation preserved equal category probability and avoided introducing asymmetry in the hard-stimulus region. Sampling was implemented in logarithmic space using MATLAB to generate probability weights that increased with distance from the boundary on the manipulated side. All other task parameters and procedures were identical to those described for the main asymmetric conditions.

### Symmetric distributions

To test whether altering the density of stimuli near the boundary affected boundary representation, we used symmetric distributions in which both categories were over- or under-represented near the decision boundary at 0.

#### Unimodal symmetric

A unimodal distribution over-sampled hard stimuli (close to the boundary) while under-sampling easy stimuli. If over-exposure improved boundary estimation, we would expect sharper psychometric slopes. However, rats showed no change in performance compared to uniform sampling (Figure S3C).

#### Bimodal symmetric

A bimodal distribution under-sampled hard stimuli and over-sampled easy stimuli. In this context, rats showed similar performance to the uniform condition.

In these manipulations the paradigm used discrete stimulus values (*n* = 8 and the probability mass functions for unimodal and bimodal contexts were:

1. Unimodal: prob = [0.0191, 0.0690, 0.1625, 0.2494, 0.2494, 0.1625, 0.0690, 0.0191]
2. Bimodal: prob = [0.0907, 0.1593, 0.1593, 0.0907, 0.0907, 0.1593, 0.1593, 0.0907]

### Mouse behavior

### Behavioural Apparatus

Behaviour was controlled using a combination of custom-built hardware and software. A HARP Sound Card (OEPS-7265) delivered auditory stimuli through in-house–designed and soldered speakers, capable of producing both pure tones and white noise. A HARP Behaviour Board (OEPS-7240) controlled solenoid valves for water delivery, while a capacitive board (Adafruit 485-136) connected to the behaviour board monitored licking activity. The head-fixation unit was adapted from the International Brain Laboratory (IBL) design, with the key modification of replacing the running wheel with a custom 3D-printed paw support platform. Water spout holders were also custom-designed and 3D-printed for precise positioning. Video data were acquired using CMOS infrared cameras placed inside the experimental boxes. All behavioural protocols were implemented in Bonsai-RX (Lopes et al. 2015), a visual reactive programming language enabling precise real-time control of task parameters and rapid protocol adjustments.

### Behaviour

The behavioural task was a two-alternative forced-choice auditory categorisation task. Mice were head-fixed and presented with a stimulus—either a pure tone or white noise—located on either side of a category boundary set at the midpoint of the stimulus range. Mice reported whether the stimulus was higher or lower than the boundary by licking the right or left water spout. The mapping between spouts and category labels (A/B) was counterbalanced across mice. For pure tones (Figure S2), the relevant dimension was frequency (4–16 kHz); for white noise, the relevant dimension was amplitude (50–82 dB SPL). Using separate cohorts for the two modalities confirmed that observed effects were not frequency-specific but generalised across auditory domains. For clarity, we refer to the lowest stimulus value as −1, the highest as 1, and the category boundary as 0. Training was conducted once daily, with each session lasting 30–60 minutes.

A shaping protocol was used to train the animals through the following stages:

#### 1. Stage 1

Passive association of extreme stimuli (−1 or 1, every 10-15 s) with the corresponding water spout, with automatic delivery of 5 *µ*L water after the sound (takes 1 day).

#### 2. Stage 2

Active side selection, where a lick on a given spout triggered the associated extreme stimulus and subsequent water reward of 5 *µ*L (takes 1 day).

#### 3. Stage 3

Side persistence requirement, where animals had to lick the same spout at least three times before switching, to reduce early side bias (around 2 days).

#### 4. Stage 4

Forced-choice with only two stimuli (−1 and 1) followed by 300 ms gap of silence (go-cue), preceding a 10 s response window. Correct responses were rewarded, incorrect responses punished with a 3 s timeout. Trials were separated with a 2-second inter-trial-interval (ITI). Therefore, incorrect licks effectively doubled the duration of the ITI. Within 6-10 days, animals usually reached >80% performance on this stage.

#### 5. Stage 5

Full task with the continuous range of stimuli (−1 to 1) and the same reward/punishment contingencies. Once animals reached *>*80% correct, experimental manipulations of stimulus distribution commenced.

On the final stage, mice performed on average 352 valid trials per session.

#### Stimulus distributions

The uniform distribution was generated by random continuous sampling across the entire range (−1 to 1). Asymmetric distributions were generated using truncated exponential functions, either *p*(*x*; *λ*) = *λe*^*λx*^ + *k* between −1 and 0 for the hard-A asymmetric distribution, or *p*(*x*; *λ*) = *λe*^−*λx*^ + *k* between 0 and 1 for the hard-B asymmetric distribution, where *k* renormalises the truncated distribution. The parameter *λ* was chosen so that the probability of sampling at the boundary was twice that of the uniform distribution. Categories were presented with equal overall probability.

#### Stimulus duration manipulation

To manipulate the reliability of sensory evidence, we varied the duration of the auditory stimulus in a subset of sessions. The standard stimulus duration (400 ms) was compared with shorter durations (150–200 ms), which increase perceptual noise by reducing the integration time of sensory input. All other task features, including stimulus range, distribution, and reward contingencies, were held constant. This manipulation allowed us to test the prediction of the normative model that higher sensory noise should increase bias magnitude in asymmetric contexts.

### Psychometric curve fitting

To quantify behavioural performance, we computed the probability of choosing category B as a function of the stimulus value, parameterised by its distance from the decision boundary (Wichmann and Hill 2001).

We have used two different methods to compute the psychometric curves: (i) a simple parametric fit using a four-parameter cumulative Gaussian function, which allows direct comparison of estimated parameters (mean, slope, and lapse rates) across task conditions, and (ii) a flexible spline fitting, which makes minimal assumptions about the underlying shape and is therefore well-suited for direct comparison between empirical data and the predictions from the normative model (see Normative Model’s Psychometric Function).

### Four-parameter cumulative Gaussian psychometric function

We modelled the relationship between stimulus value (distance to boundary) and choice probability with the following psychometric function:

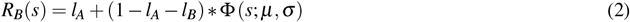

where *R*_*B*_(*s*) is the predicted probability of choosing category B for stimulus *s. l*_*A*_ and *l*_*B*_ are the lapse rates for categories *A* and *B*, respectively; and Φ is the Gaussian CDF with mean *µ* and standard deviation *σ* (inversely related to the slope of the psychometric curve at its midpoint). Throughout the figures, we show psychometric fits for individual subjects (thin lines) and a fit to the group average across individuals (thick lines).

At the individual subject level, we fitted binary choices (0 or 1) as a function of stimulus value using maximum likelihood estimation. The negative log-likelihood (NLL) was defined as:

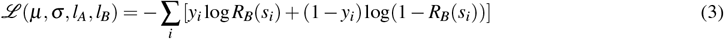

where *y*_*i*_ is the observed choice on trial *i* and *R*_*B*_(*s*) is the probability of choosing category B given stimulus *s*, given by Equation 2.

Parameters were optimised using `scipy.optimize.minimize`with the `L-BFGS-B`optimisation method, with the following bounds on the parameters: *µ* ∈ [−1.0, 1.0], *σ* ∈ [0.01, 10.0], *l*_*A*_ ∈ [0.0, 0.5], and *l*_*B*_ ∈ [0.0, 0.5].

### Spline psychometric function

To compare the predictions from our normative model, we used a spline logistic regression model to flexibly estimate “empirical” psychometric functions for each animal, on each stimulus distribution. Our approach assumes a continuous function with varying degrees of smoothness but, unlike typical parametric psychometric models, does not impose any particular shape. On each trial, the log odds of a positive vs. negative action *a* in response to stimulus *s* are given by a piecewise linear spline *f* (with *k* − 1 = 100 pieces):

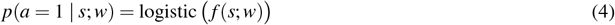

*f* is a linear combination of fixed B-spline basis functions {*ϕ*_1_, …, *ϕ*_*k*_} of order 2, with coefficient vector *w* ∈ ℝ^*k*^. The associated knot points {*t*_1_, …, *t*_*k*_} are strictly increasing and cover the entire stimulus range, with even spacing Δ = *t* _*j*+1_ −*t*_*j*_.

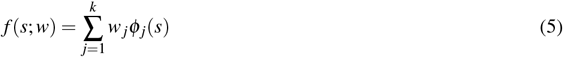

Let (*s*_1_, *a*_1_), …, (*s*_*n*_, *a*_*n*_) be a training set containing observed stimuli and actions over *n* trials, which are assumed i.i.d.. Coefficients are fitted by minimising the negative log likelihood, plus a smoothness penalty *R* (with penalty strength *λ* ≥ 0):

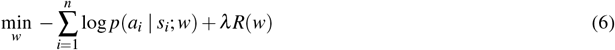

The smoothness penalty measures the sum of squared differences in slope between adjacent pieces of the spline (whose derivative is *f* ^*′*^). Thus, higher penalties encourage smoother log odds, and the spline logistic regression model approaches linear logistic regression as *λ* → ∞.

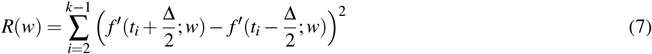

Coefficients were optimised using a trust-region algorithm (Powell’s dogleg method). For each dataset, the smoothness penalty was fitted using 10-fold cross validation. We selected the penalty strength *λ* that maximised test set likelihood, from among 19 logarithmically spaced values from 10^−1^ to 10^8^. Final coefficients were then fitted to the entire dataset using the optimal smoothness penalty.

### Point of Subjective Equality (PSE)

We defined Point of Subjective Equality (PSE) as the stimulus value at which the psychometric curve crosses 0.5 on the y-axis, i.e. the model is equally likely to choose either category. Given the four-parameter cumulative Gaussian (Equation 2), we defined PSE as the stimulus value where *R*_*B*_(*s*) = 0.5; we rescaled the 0.5 level on the underlying Gaussian CDF Φ:

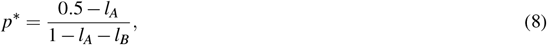

then invert the Gaussian CDF with location *µ* and scale *σ*:

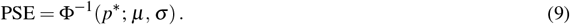

Computation was implemented with SciPy’s `scipy.stats.norm.ppf`; PSEs falling outside the stimulus range [−1, 1] were discarded.

### Time-resolved adaptation and convergence metric

After each asymmetric-to-asymmetric switch in statistical context (hard-A ↔ hard-B), in humans, rats, and mice, we estimated psychometric functions in contiguous, non-overlapping 50-trial bins aligned to the switch (bin 0 = trials before the switch, bin 1 = trials 1–50 after the switch, etc.). Within each bin, choices were fit with the four-parameter cumulative Gaussian (Equation 2), and the point of subjective equality (PSE) was computed as described above (see Point of Subjective Equality). For each switch, we then normalised the post-switch PSE trajectory by the pre-switch baseline and the normative prediction for the new statistical context, as defined in Equation 1:

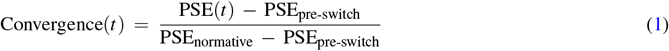

Bins yielding undefined PSEs (e.g., due to lapse constraints) were omitted. The pre-switch baseline PSE_pre-switch_ was estimated from the trials immediately preceding each switch: the last 50 trials for humans (one 50-trial bin) and the last 250 trials for rats and mice. In the human task, trials were organised in 50-trial blocks; at each block boundary the statistical context could switch between hard-A and hard-B, so participants experienced multiple uncued switches. In rodents, each rat underwent a single asymmetric-to-asymmetric switch in a single direction (either from hard-A to hard-B or from hard-B to hard-A), and mice underwent one or, for some subjects, two switches. For group summaries and statistical tests, we computed *subject-level* means, so each subject contributed a single trajectory/estimate regardless of the number of switches.

### Update matrices

To examine how stimuli from previous trials influenced current choices, we computed *update matrices* following the approach of Lak et al. 2020 and Akrami et al. 2018. Each update matrix quantified change in choice probability (probability of choosing category B) for a given current stimulus, relative to the overall psychometric function, when conditioned on the stimulus presented in the previous rewarded trial.

#### Trial selection

We first filtered the data to include only trials that met the following criteria: (i) the current trial immediately followed a rewarded trial; (ii) the subject provided responses in both the current and the previous trials; and (iii) the current trial was not the first trial of a session.

#### Stimulus intervals

The stimulus space, normalised to the decision boundary, *S* = [−1, 1], was uniformly partitioned into eight equal-width intervals *I*_1_, …, *I*_8_. The first seven intervals were half-open:

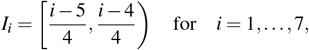

and the final interval was closed on the right:

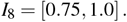

#### Overall and conditional psychometric curves

From the selected trials, we computed the *overall post-correct psychometric curve* (or grand average) defined as the proportion of category B choices as a function of current stimulus. We fitted a four-parameter cumulative Gaussian (Equation 2) to the *overall post-correct psychometric curve* as detailed above (see Psychometric curve fitting)

Similarly, eight *conditional post-correct psychometric curves* were computed, one for each possible previous stimulus interval. For the *j*-th conditional curve, only the trials for which previous stimulus *s*_*t*−1_ fell within interval *I*_*j*_ were included. Each conditional curve was fitted in the same manner as the overall curve.

#### Update matrix computation

We then constructed an 8×8 update matrix, denoted as *U*_*i j*_, where *i* indexes the current trial’s stimulus interval and *j* indexes the previous trial’s stimulus interval. Each entry *U*_*i j*_ equals the conditional post-correct psychometric value at the midpoint of the current interval for trials whose previous stimulus was in *I*_*j*_, minus the corresponding overall post-correct psychometric value at that same midpoint.

Thus, *U*_*i j*_ captures the deviation in choice probability for current interval *I*_*i*_ attributable to the previous stimulus being in interval *I*_*j*_. Positive values indicate a higher-than-average probability of choosing category B under that condition, and negative values indicate a lower-than-average probability. Each update matrix was visualised as a heat map, with colour representing the magnitude and sign of these deviations, revealing systematic influences of past stimuli and outcomes on current decision-making. Matrices are organised such that *U*_11_ is on bottom left corner and *U*_88_ is on top right corner.

The conditional matrix is constructed in the same way as the update matrix, except that the overall post-correct psychometric value is not subtracted from the conditional post-correct values.

## Computational models

### Normative model

The normative model formalises how an ideal decision maker should behave in our task, given both the external structure of the environment (category and stimulus distributions, reward contingencies) and internal sensory and decision noise. The model specifies how stimuli are encoded with noise, how beliefs about category membership are computed, and how choices are generated to maximise expected reward. It also describes how behaviour would appear to an external experimenter, as a psychometric function probabilistically linking choice behaviour to presented stimuli. Similar to (Whiteley and Sahani 2008), the model establishes a principled framework to quantify expected behaviour if agents have already learned the correct priors and integrate them optimally with noisy sensory evidence.

### Data generating process

On each trial within a statistical context, the experimenter generates a real-valued stimulus *s* from a binary category *c*, according to the Bernoulli category distribution *p*(*c*) and stimulus distribution *p*(*s* | *c*). In our experiments, *negative* (*c* = 0) and *positive* (*c* = 1) category labels correspond to stimulus categories *A* and *B*, respectively.

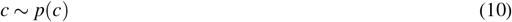

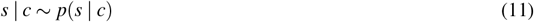

The stimulus elicits a stochastic neural response *x* in the sensory system:

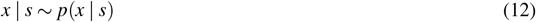

Assume the sensory system smoothly encodes the stimulus, so neural activity falls near a 1d manifold isomorphic to the stimulus space. Position along the manifold can therefore be parameterised using the same coordinates as the stimulus itself, and neural activity can be summarised with a one-dimensional variable, given by the stimulus plus stimulus-dependent noise *ε*:

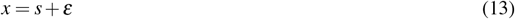

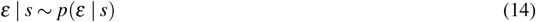

Given the observed the sensory response, the agent generates an internal binary choice *ĉ* using policy function *π*: ℝ → {0, 1}. In our experiments, *negative* (*ĉ* = 0) and *positive* (*ĉ* = 1) choice labels represent the intention to make response *A* or *B*, respectively.

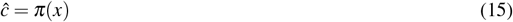

Behavioural output is subject to decision noise, which may flip the internal choice with some probability. This yields a binary action *a*:

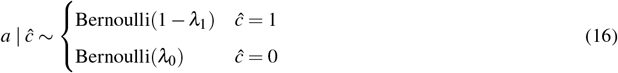

Positive lapse rate *λ*_1_ and negative lapse rate *λ*_0_ give the probability of flipping a positive or negative choice, respectively. Lapse rates are assumed not to be extreme (*λ*_1_ + *λ*_0_ *<* 1).

After responding, the agent receives a stochastic, real-valued reward *r*, depending on the stimulus category and action:

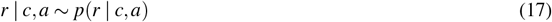

Let 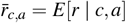 denote the mean reward for each category and action. Mean reward is assumed to be greater for category-matching actions (*a* = *c*) than non-matching actions (*a* ≠ *c*) (if category/action labels cannot be defined this way, then the task can trivially be solved by always taking the same action).

All variables are assumed to be independent across trials, and identically distributed within each statistical context. For each trial, the joint distribution factorises as:

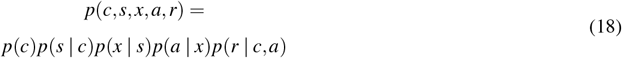

### Behavioural policy

The agent seeks to maximise expected reward, given the information available to it on each trial (i.e. the neural response) and its knowledge of how the world generally works. The latter takes the form of a probabilistic world model that describes the agent’s beliefs about how the relevant variables are related. The optimal policy 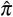 produces the internal choice that maximises this expectation:

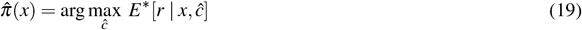

In this setting, a deterministic policy is optimal; choosing stochastically cannot increase expected reward. We use the superscript ∗ to denote subjective quantities and distributions pertaining to the agent’s world model, which may differ from the true data generating process above, but is assumed to share the same general dependence structure. The world model is also assumed to be static across trials, such that no learning occurs; we consider inference and decision making in an agent already familiar with the structure of the world.

Expected reward (according to the world model) is obtained by marginalising out the stimulus category and action to obtain:

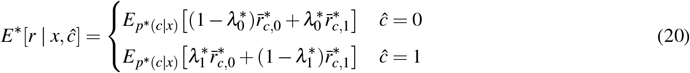

The optimal policy (Equation 19) then admits the following solution:

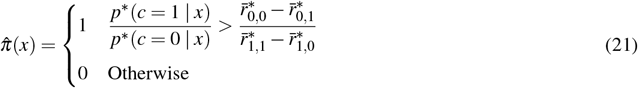

Thus, the optimal choice is based on comparing the posterior odds (of the positive vs. negative stimulus category, given the observed sensory response) to a threshold. The threshold considers the difference in mean reward for an action that matches vs. differs from the stimulus category, and is given by the ratio of this quantity for the negative vs. positive stimulus category.

### Decision boundary

The policy function partitions sensory response space into regions that map to a positive or negative choice. Let {𝒳_1_, …, 𝒳_*k*_} be disjoint sub-intervals that map to a positive choice under 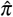, and *α*_*i*_, *β*_*i*_ be left/right endpoints of each 𝒳_*i*_. The decision boundary contains all finite endpoints, which are obtained by numerically solving for sensory response values *x* with threshold-level posterior odds (working in log space for numerical stability):

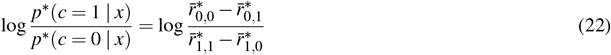

Left vs. right endpoints can be distinguished by whether the posterior odds are increasing (left) or decreasing (right) at each solution. Furthermore, we have *α*_1_ = −∞ if the posterior odds decrease at the least point in the decision boundary, and *β*_*k*_ = ∞ if they increase at the greatest point. The log posterior odds are computed as follows, with expectations calculated using numerical quadrature:

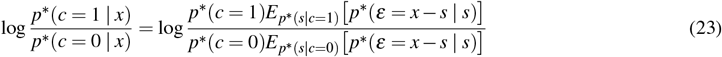

### Psychometric function

The psychometric function *ψ* summarises the agent’s behaviour from the perspective of an experimenter who observes the stimulus and action, but for whom the neural response and internal choice are treated as unknown. *ψ*(*s*) yields the probability of a positive action, given the stimulus, which is obtained by marginalizing out the unknown neural response:

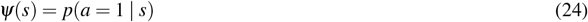

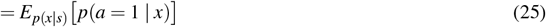

As above, let {𝒳_1_, …, 𝒳_*k*_} be disjoint sub-intervals of sensory response space that map to a positive choice under the policy function. Then the psychometric function can be expressed as:

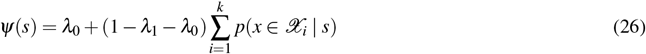

The probability of a sensory response in the *i*th sub-interval (with left/right endpoints *α*_*i*_, *β*_*i*_) is:

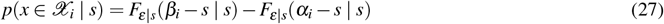

where *F*_*ε*|*s*_ is the CDF of the sensory noise, given the stimulus. Note that distributions here pertain to the true data generating process, not the agent’s subjective world model.

### Behaviour across statistical contexts

As with experimental subjects, we studied behaviour of the normative model across multiple statistical contexts. Each context was defined by a different stimulus distribution *p*(*s* | *c*), but shared common category probabilities and reward contingencies. Due to the change in stimulus distribution, the normative model yields a different behavioural policy and psychometric function for each context.

All distributions were chosen to match the behavioural task: stimulus distributions as described above for mouse/rat/human subjects, equiprobable categories, and deterministic, constant reward for category-matching actions (otherwise none). We set the subjective world model equal to the true data generating process, so the normative model behaved as an ideal observer that understands the statistical structure of the world and the context in which each decision is made, but is constrained by sensory and decision noise.

Sensory and decision noise were assumed to be i.i.d. across contexts, so all contexts shared the same lapse rates and sensory noise distribution. To flexibly model sensory responses, we used Gaussian sensory noise *ε* | *s* ∼ 𝒩 (0, *σ* ^2^(*s*)) with stimulus-dependent variance. Noise variance *σ* ^2^(*s*) was a power of a linear function of the stimulus, with the slope, intercept, and exponent controlled by free parameters. The function was parameterised to constrain the exponent to the interval (0.2, 20), and the variance to exceed 2.5 × 10^−3^ for all stimuli.

### Fitting noise distributions

The choice of noise distributions may strongly affect the model’s behaviour. Furthermore, noise may differ across species and individuals. We therefore fit sensory and decision noise distributions to the data for each subject. We employed a cross validation approach, where results calculated for each context used noise distributions fit to data from all *other* contexts (i.e. excluding the current context).

We examined hard-A asymmetric and hard-B asymmetric contexts for all species, in addition to the uniform context for mice. For each subject, we included data from all contexts with at least 200 trials and accuracy at least 70%. Analysis proceeded only for subjects that met inclusion criteria on both hard-A asymmetric and hard-B asymmetric contexts.

Sensory and decision noise parameters were fit as follows. A training set { (*s*_*n*,*t*_, *a*_*n*,*t*_) } contains pairs of recorded stimuli and actions for each trial *t* within each context *n*. Let *θ* be a parameter vector for the sensory noise variance function and *λ* = (*λ*_0_, *λ*_1_) be the lapse rates. A given set of parameters determines a psychometric function *ψ*_*n*_ for each context, which is computed using the noise distributions specified by the parameters and the stimulus distribution defined by the context.

The psychometric functions yield the probability of each observed action (given the stimulus):

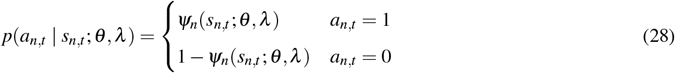

Parameters were fit by maximizing the log likelihood, which we treated as a nested optimisation problem:

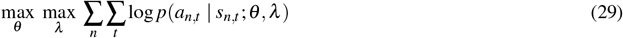

For any set of sensory noise parameters, the optimal lapse rates (inner problem) were found using the L-BFGS algorithm, with lapse rates constrained to the interval (10^−6^, 0.5 − 10^−6^). This approach was computationally efficient, since the log likelihood is differentiable with respect to the lapse rates. Sensory noise parameters (outer problem) were optimised using a derivative-free algorithm (Powell’s method). To avoid local minima, the best solution was retained after solving the problem 10 times with random initial parameters. Preliminary numerical experiments suggested that further random restarts were unlikely to improve the solution.

### Reward rate

Task performance can be quantified using the reward rate (i.e. expected reward on each trial). This is obtained by marginalizing out the stimulus, category, and action, and can be computed from the psychometric function, together with the true stimulus, category and reward distributions defined by the behavioural task:

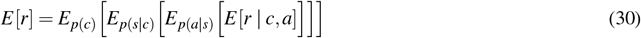

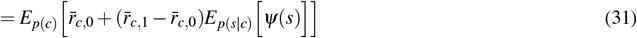

For the reward structure used in our experiments (deterministic unit reward if the action matches the category, otherwise zero):

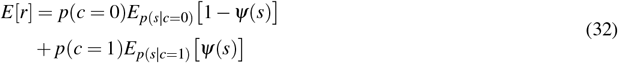

The expectations above were computed using numerical quadrature.

### Learning models

In both models, decisions are based on a perceived stimulus, which represents the internal sensory estimate after noise corruption and repulsion adjustment. On each trial *t*, we sampled 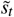 from a normal distribution centred at the true stimulus value *s*_*t*_, with variance 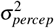. We then applied an additive repulsion term (a function of the current stimulus, and the previous trial’s perceived stimulus), defined as:

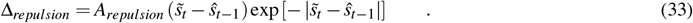

where *A*_*repulsion*_ controls the amplitude of the repulsion effect. The final perceived stimulus used for decision-making on trial *t* is then given by: 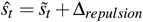.

Both models share two parameters: perceptual noise *σ*_*percep*_ ≥ 0 and repulsion amplitude *A*_*repulsion*_ ≥ 0. In the stimulus-category model, additional parameters include the belief inertia *γ* and the update kernel width *σ*_*update*_. In the boundary-estimation model, we instead have two parameters controlling updates: *η*_*learning*_ and *η*_*relax*_ (see below).

The stimulus space *S* is defined to include all possible perceived stimuli *ŝ*_*t*_. While true stimulus values fall within [−1, 1] range, added noise and repulsion extend this range causing the range of perceived stimulus, *ŝ*_*t*_ to be larger. Given that 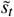 is drawn from a Gaussian distribution centered at *s*_*t*_ with standard deviation *σ*_*percep*_, almost all values (≈ 99.999998%) lie within *±*6*σ percep* of *s*_*t*_. Thus,

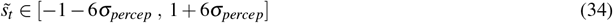

Additionally, the repulsion term affects range of perceived stimulus, as well. Since 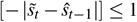, repulsion term is bounded as below:

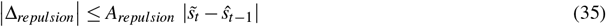

A conservative bound is obtained by assuming the extreme case when 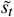 and *ŝ*_*t*−1_ are on opposite ends of the stimulus range. Therefore:

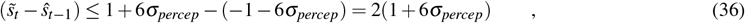

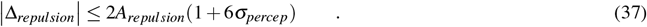

Hence, the overall stimulus space *S*, is defined as:

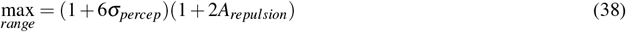

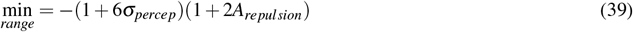

Finally, to mirror empirical data, on trials where the subject did not respond, we neither update beliefs nor register a model choice. Instead, we assign a `NaN`to represent a no-response trial in the simulated agent.

### Stimulus-category model

In the stimulus-category model, the agent maintains probabilistic belief distributions over the stimulus space for each category, denoted *p*(*S*|*A*) and *p*(*S*|*B*). On each trial *t*, the agent is presented with a stimulus *s*_*t*_ and outputs a binary decision, A or B. On each trial, after obtaining the perceived stimulus *ŝ*_*t*_, the agent computes posterior probabilities for each category; decision A with probability

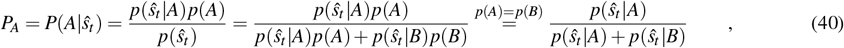

and decision B with probability

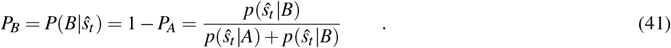

To capture stochastic nature of decisions, the agent’s binary choice *Y*_*t*_ is modelled as a Bernoulli random variable with probability *P*_*B*_;

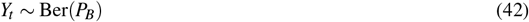

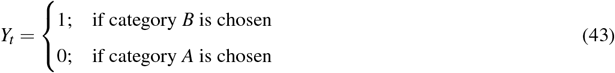

Upon receiving feedback, the belief distribution for the chosen category *C* ∈ *A, B* (either *p*(*S*|*A*) or *p*(*S*|*B*)) is updated. If the choice was correct, the distribution is updated with a positive feedback kernel; if incorrect, it is adjusted with a negative kernel:

1. **Positive feedback** (correct choice):

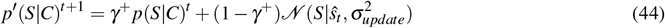
2. **Negative feedback** (incorrect choice):

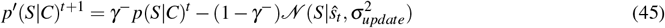

The parameters *γ*^+^ and *γ*^−^ determine the weight given to existing beliefs versus new evidence. For simplicity, we assume *γ* = *γ*^+^ = *γ*^−^. The parameter 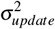 determines the variance of Gaussian kernel used during belief updating, controlling how locally or diffusely new information influences the belief distribution. Smaller values yield more focused updates around *ŝ*_*t*_, while larger values produce broader adjustments.

To ensure non-negativity, any negative values in the updated unnormalised distribution are shifted upwards:

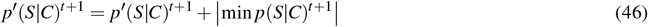

Finally, the distribution is normalised:

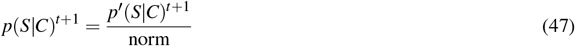

where the normalisation constant ensures the distribution (over *S*) integrates to 1.

Initial category beliefs are Gaussian: for A, mean −0.75 and SD 0.5; for B, mean 0.75 and SD 0.5:,

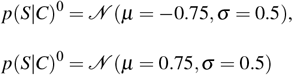

### Boundary-estimation model

In the boundary-estimation (BE) model, the agent maintains a probabilistic belief about the location of the decision boundary rather than representing full category distributions. Specifically, this belief is described by a probability distribution, *B*_*t*_(*S*), over the stimulus space *S* at trial *t*. The belief distribution is updated after each trial through two mechanisms: a feedback-dependent learning term, Δ_*learning*_, and a relaxation (forgetting) term, Δ_*relax*_. The learning term is defined as:

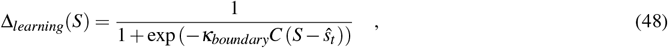

where *κ*_*boundary*_ controls the precision of the update– higher values yield more localised updates around *ŝ*_*t*_. Hence, we set *κ*_*boundary*_ as the inverse of the perceptual noise standard deviation, *σ*_*percep*_. The variable *C* indicates the category of true stimulus (*C* = 1 for category B and *C* = −1 for category A) and reflects the agent’s knowledge after feedback. After feedback, the updated (unnormalised) boundary belief is:

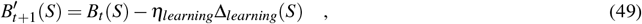

where *η*_*learning*_ is the learning rate governing the strength of the feedback update. The relaxation term gradually pulls the updated boundary belief back towards a uniform baseline, acting as a forgetting term. The amplitude of this relaxation term is given by the distance between the model’s current belief of boundary and the uniform distribution over [−1, +1]

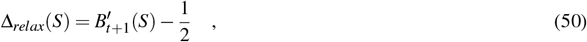

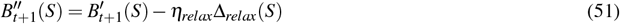

where *η*_*relax*_ controls the degree of this reversion toward baseline. If 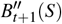 contains negative values, it is shifted upwards to maintain non-negativity:

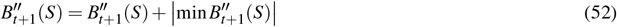

The belief is then normalised:

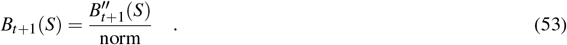

ensuring it integrates to one over *S*.

At each trial, the agent bases its decision on its current belief before updating. To determine the agent’s choice at trial *t*, we compute the probability of choosing category *B, P*(*B*|*ŝ*_*t*_), which corresponds to the probability that the boundary is located below the perceived stimulus *ŝ*_*t*_:

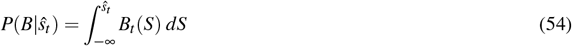

To capture the stochastic nature of decisions, the agent’s choice *Y*_*t*_ is modelled as a Bernoulli random variable with probability *P*(*B*|*ŝ*_*t*_), *Y*_*t*_ ∼ Ber(*P*(*B*|*ŝ*_*t*_)), and the mapping of the random variable to categories is as shown in Equation 43. The initial boundary belief is set to a uniform distribution, reflecting an uninformed state with no prior exposure or expertise.

### Model fitting (BE and SC model)

To ensure robust parameter estimation when fitting psychometric curves, we imposed constraints on all four parameters of the psychometric function *R*(*B*) (see Psychometric fitting), restricting them to appropriate ranges (Table 1). We also set the maximum number of function evaluations to 10,000 to prevent premature stopping during optimisation.

**Table 1:** Parameter Constraints for Psychometric Curve Fitting.

| Parameter | Description | Bounds |
| --- | --- | --- |
| $\mu$ | Threshold | $[-1, 1]$ |
| $\sigma$ | Inverse slope | $[0.01, 10]$ |
| $\gamma$ | Lower lapse rate | $[0, 0.5]$ |
| $\lambda$ | Upper lapse rate | $[0, 0.5]$ |

To quantify how closely model responses matched subject behaviour, we defined an error measure based on the Frobenius distance. Let *D* be the conditional matrix derived from subject data, *M* the model conditional matrix (see Methods), and number of columns that are fully valid, i.e., have at least one `non-NaN`element, is *n* (in update matrices, it implies all elements in a column are valid, since we can not have a single element of `NaN`). Then, the error *E* is:

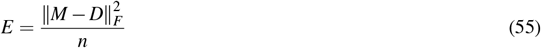

This approach ensures that columns without valid trials (i.e., no conditioned data) do not bias the error calculation, providing a fair comparison even with incomplete data.

We used a grid search to find the parameter set minimising this error, systematically exploring combinations defined in Table 2 and Table 3

**Table 2:**
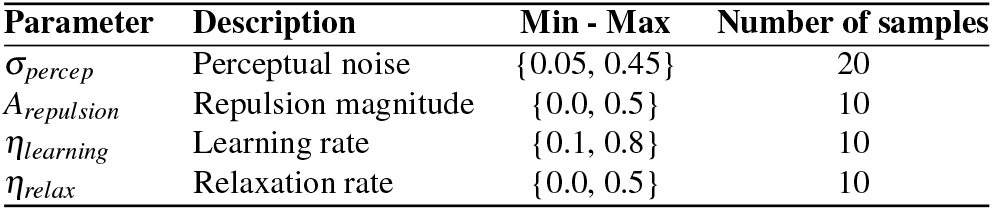
Grid Search Parameters for Boundary Estimation model.

| Parameter | Description | Min - Max | Number of samples |
| --- | --- | --- | --- |
| $\sigma_{percep}$ | Perceptual noise | {0.05, 0.45} | 20 |
| $A_{repulsion}$ | Repulsion magnitude | {0.0, 0.5} | 10 |
| $\eta_{learning}$ | Learning rate | {0.1, 0.8} | 10 |
| $\eta_{relax}$ | Relaxation rate | {0.0, 0.5} | 10 |

**Table 3:** Grid Search Parameters for Stimulus Category model.

| Parameter | Description | Min - Max | Number of samples |
| --- | --- | --- | --- |
| $\sigma_{percep}$ | Perceptual noise | {0.05, 0.45} | 20 |
| $A_{repulsion}$ | Repulsion magnitude | {0.0, 0.5} | 10 |
| $\gamma$ | Belief inertia | {0.40, 0.93} | 9 |
| $\sigma_{update}$ | Width of update | {0.1, 0.7} | 10 |

For each parameter set, we first trained the models on 1,000 simulated trials assuming complete responses, allowing them to reach expert-level performance comparable to trained human or rodent subjects. We then exposed these “expert agents” to the same sequence of stimuli used in the empirical data to generate corresponding update matrices. Therefore, when comparing the update matrices between observed and simulated data, both matrices were generated from expert agents, and both matrices were generated from the same sequence of stimuli.

#### Model selection

To identify the optimal model for each subject, we performed 2-fold cross-validation to generate an average “test score” for each participant, for each model. For each fold, we fit model parameters using the grid-search procedure described above, and evaluated performance (test error) on the held-out fold. We repeated this twice (once per fold), averaging the resulting test errors to obtain a mean cross-validated error for that model and subject. To further enhance robustness, this entire cross-validation procedure was repeated 8 times with different random seeds, yielding 8 average test errors per model per participant. We compared the distribution of test errors across the two models: stimulus-category and boundary-estimation. To formally assess differences, we performed an ANOVA on test errors. If the test revealed a significant effect (p < 0.05), we identified the model with the lowest average error as the best-fitting model for that subject. If no significant difference was found, we concluded that no single model clearly outperformed the others for that participant.

## Statistical Analysis

All statistical analyses were performed in Python. For each dataset, normality was assessed with the Shapiro–Wilk test. When the null hypothesis of normality was rejected (*p <* 0.05), non-parametric tests were used and results are reported (and plotted) as median [IQR]. When normality was not rejected, parametric tests were used and results are reported as mean *±* SEM. Multiple comparisons were adjusted as specified in each analyses. All statistical tests, corrections and p-values are reported in the main text or figure legends. For each figure, the sample size (n) and what it represents are specified either in the figure or its legend.

In Figure 1D, statistical tests differed across species depending on whether the same or different individuals contributed to the distributional conditions. For rats and humans, each subject was exposed to only one distribution condition, so the groups were independent; therefore, we compared them using a Kruskal–Wallis test followed by pairwise Mann–Whitney *U* tests with Bonferroni correction when significant (two-sided, *α* = 0.05). For mice, the same subjects were tested across multiple distributions, so the groups were paired; therefore, we used a Friedman test followed by pairwise Wilcoxon signed-rank tests with Bonferroni correction when significant (two-sided, *α* = 0.05).

In Figure 5B (rats, mice, and humans), two paired conditions were compared using a paired-samples *t*-test (two-sided, *α* = 0.05).

In Figure 5C, species were compared pairwise (rats–humans, rats–mice, mice–humans) using Mann–Whitney *U* tests with Bonferroni correction for the three comparisons (two-sided, *α* = 0.05).

In Figure S3A, rats were compared across two independent groups using a Mann–Whitney *U* test (two-sided, *α* = 0.05). For humans, we compared two paired conditions using a Wilcoxon signed-rank test (two-sided, *α* = 0.05).

In Figure S3B, two paired conditions were compared using a paired-samples *t*-test (two-sided, *α* = 0.05).

In Figure S3C, within-subject conditions were compared using paired Wilcoxon signed-rank tests with Bonferroni correction for the two comparisons (two-sided, *α* = 0.05).

In Figure S3D, within-subject conditions were compared using paired-samples *t*-tests, separately by stimulus-duration cohort.

## Supplementary Figures

**Figure S1.**
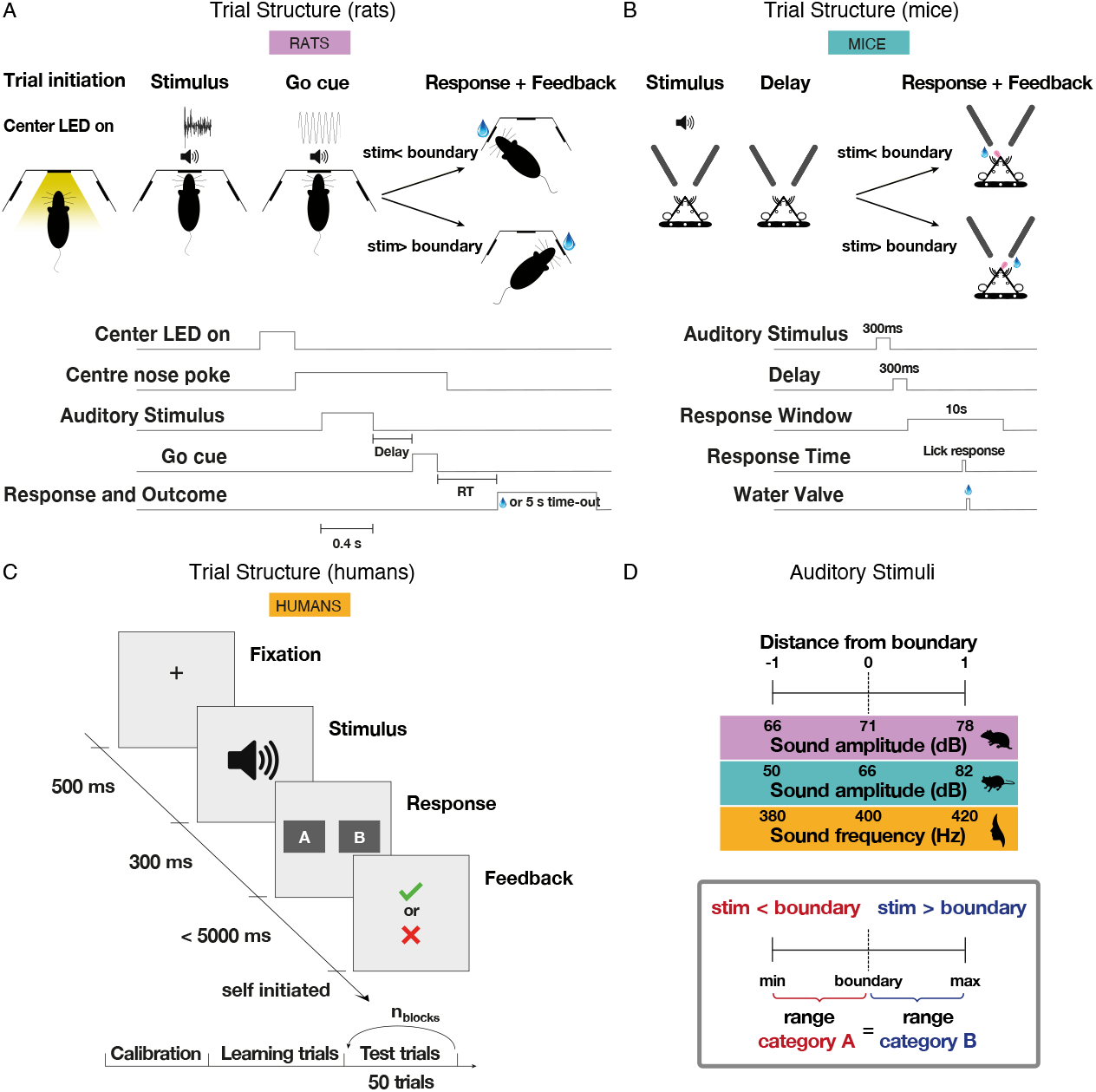
Task schematics. Full trial structure for each species (see Methods). **A**: Trial structure for freely moving rats, illustrating stimulus presentation (structured noise of varying intensity), response, and feedback. **B**: Same as (A) for head-fixed mice. **C**: Trial structure for humans, presented with pure tones of varying frequency; session outline includes calibration, training, and test phases **D**: Mapping of “distance from boundary” onto the physical stimulus dimension: sound intensity (rats and mice) and sound frequency (humans). The decision boundary was fixed at the midpoint of the stimulus range (in logarithmic scale), which was sampled continuously.

**Figure S2.**
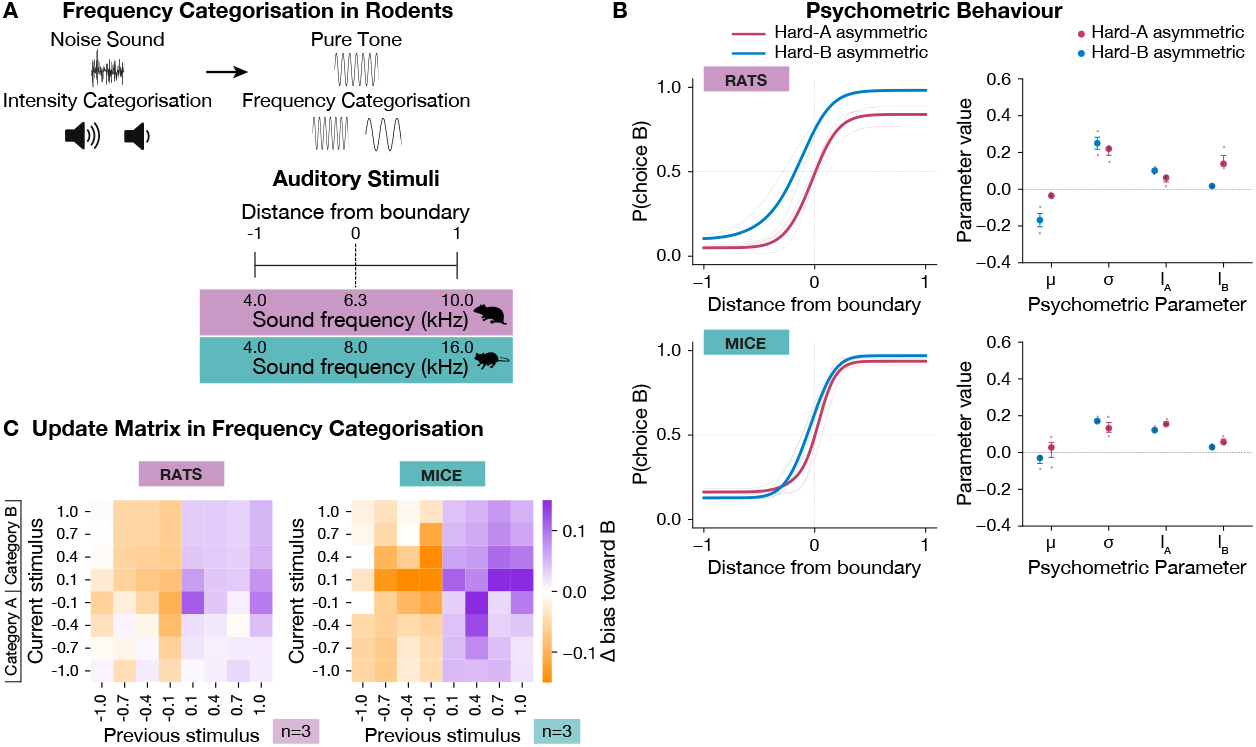
Frequency version of the sound categorisation task for rodents. Control experiment designed to match the stimulus dimension used in human subjects, who performed frequency rather than intensity categorisation. **A**: Task schematics and mapping of “distance from boundary” onto the physical stimulus dimension (sound frequency) for rats and mice. The decision boundary was fixed at the midpoint of the frequency range, which was sampled continuously. **B**: left panels show psychometric curves for all individuals (thin lines) and the average across-subject fit (thick line), shown separately for rats (top) and mice (bottom), for hard-A and hard-B contexts in frequency categorisation task, showing modulation similar to the intensity experiments. Right panels show individual parameter estimates (dots) from four-parameter psychometric fits (mean, slope, lapse rates) for hard-A and hard-B, alongside the group median and interquartile range. **C**: Empirical update matrices for rats (left) and mice (right) in the uniform context, showing choice updating patterns similar to the intensity experiments for each species.

**Figure S3.**
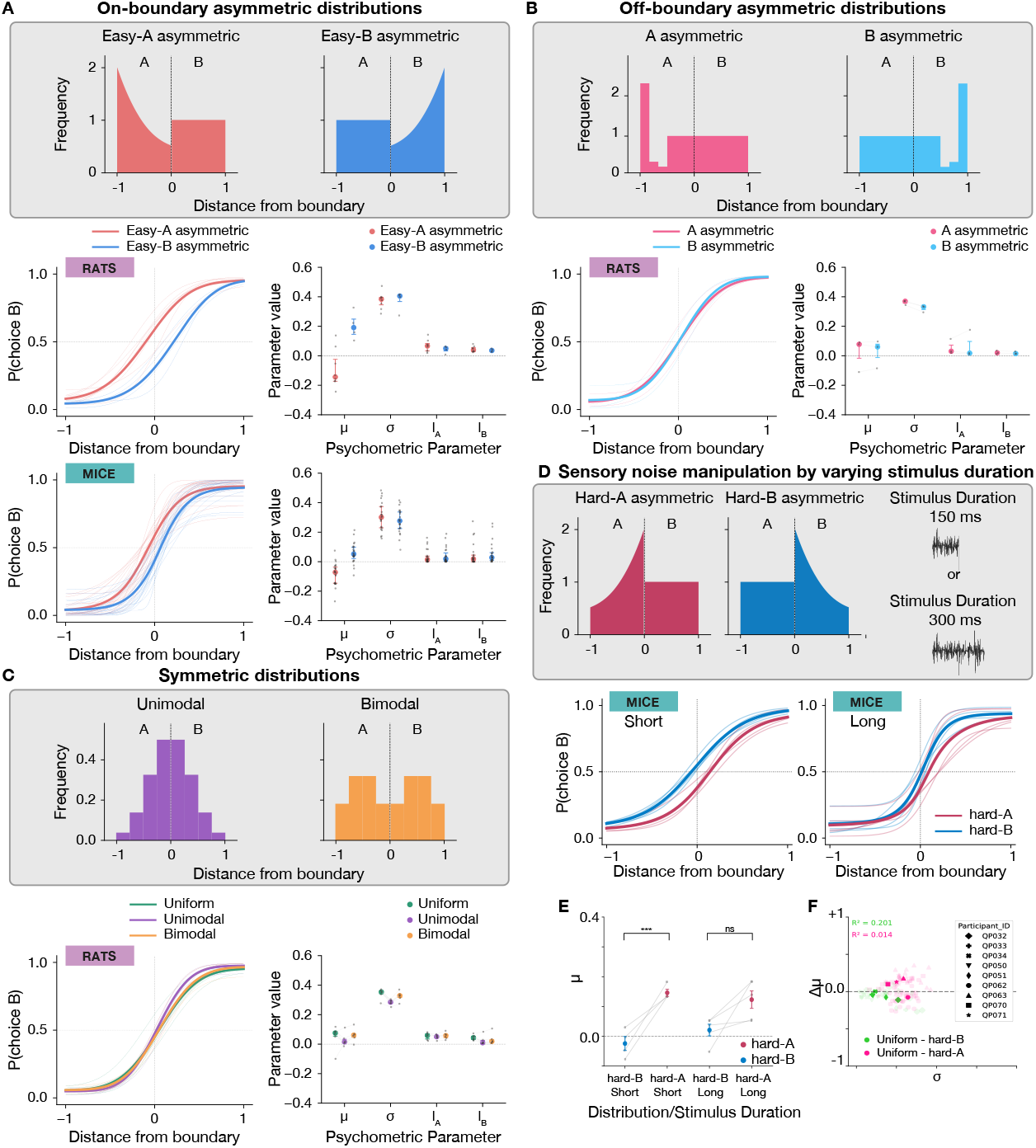
Asymmetry in sensory distribution is exploited only if needed. Biases observed in humans and rodents are solely due to the relative difference in category representation around the boundary irrespective of the shape of the non-uniform category. **A**: easy-A and easy-B contexts, in which the non-uniform category over-represented easy stimuli (far from the boundary) and under-represented hard stimuli (near the boundary). The corresponding psychometric curves for rats and humans are shown below: left panels show psychometric curves for all individuals (thin lines) and the average across-subject fit (thick line), shown separately by species and for the stimulus distributions. Right panels show individual parameter estimates from the four-parameter fits, together with the group median and interquartile range. Both humans and rats in the easy-A condition showed a bias toward category A for boundary-adjacent stimuli, while easy-B produced the opposite bias. Mann–Whitney *U* test, mu in rats: p-value = 0.01379 for easy-A (n=8) vs easy-B(n=4); Wilcoxon signed-rank test, mu in humans (n=23): p-value = 0.0000002 for easy-A vs easy-B. **B**: Off-boundary asymmetric contexts. Statistical contexts with asymmetry away from decision boundary within category A (pink) or category B (light blue), relative to the other uniformly-sampled category. Left panel shows psychometric curves for all individuals (thin lines) and the average across-subject fit (thick line), shown for the stimulus distributions. Right panel shows individual parameter estimates from the four-parameter fits, together with the group mean *±* SEM. The psychometric curves did not show a choice bias in the off-boundary asymmetrical contexts and the mean parameter of the psychometric curve fits *µ* was around zero for both distributions, indicative of a balanced performance, similar to the performance of subjects in the uniform context. Two-sided paired t-test, mu in rats (n=3): p-value = 0.72492 for A asymmetric vs B asymmetric. **C**: Over- or under-exposure to hard stimuli did not alter human or rat performances. Contexts with symmetric distributions of stimuli lead to oversampling of either easy (bimodal) or difficult (unimodal). Left panel shows psychometric curves for all individuals (thin lines) and the average across-subject fit (thick line), shown separately for the stimulus distributions. Right panel shows individual parameter estimates from the four-parameter fits, together with the group median and interquartile range. These manipulations do not vary overall performance in rats. Wilcoxon signed-rank test, Bonferroni corrected, mu in rats (n=6): p-value = 0.87500 for uniform vs unimodal; p-value = 1.00000 for uniform vs bimodal.**D**: Varying stimulus duration changes the saliency of sensory inputs, hence the perceptual reliability and noise. Bottom panels show psychometric curves for all individuals (thin lines) and the average across-subject fit (thick line), shown separately by stimulus durations and for the stimulus distributions. **E**: paired *µ* estimates from psychometric fits for each individual, pairing hard-A with hard-B within subject; circles indicate the group mean *±* SEM, and grey lines connect within-subject pairs, shown separately for stimulus-duration cohorts. Psychometric shifts tended to be smaller for longer stimulus durations (i.e., reduced sensory noise), consistent with the prediction of the normative model (paired t-test with bonferroni correction (*α* = 0.05), p = 0.010 for short duration, n = 4; paired t-test with bonferroni correction (*α* = 0.05), p = 0.058 for long duration, n = 5). Short-duration cohort have significantly steeper slopes than long duration cohort (Short mean = 0.265, Long mean = 0.150; t(7) = 2.86, p = 0.024, Hedges’ g = 1.71) **F**: Δ*µ* (shift in the psychometric mean value) vs. *σ* (reverse of slope) is shown for those 7 mice, when changing the distribution from uniform to hard-A. This suggests the shallower the slope, the larger the shift in the psychometric mean value.

**Figure S4.**
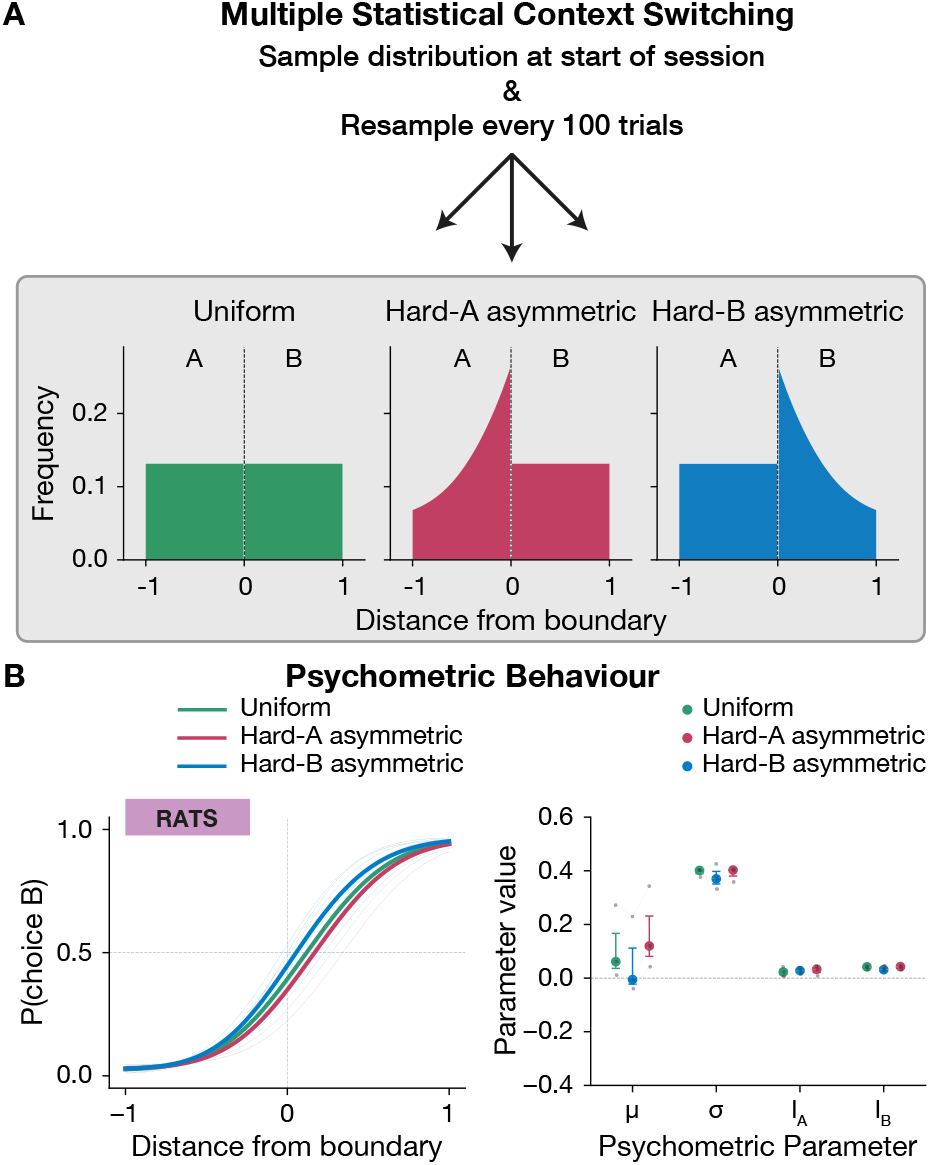
Within-session probabilistic context switching in rats. **A**: Sampling distributions used in the probabilistic switching paradigm (same as in Figure 1. **B**: Left panels show psychometric curves for all individuals (thin lines) and the average across-subject data (thick line), shown separately for each distribution in **A**. Right panels show individual parameter estimates (dots) from psychometric fits—midpoint, slope (width), and lapse rate—together with group median and interquartile range. In this within-session paradigm, analogous to that used in humans, rats adapted flexibly to the prevailing distribution, comparable to the static-context condition (see Figures 3 & 5).

**Figure S5.**
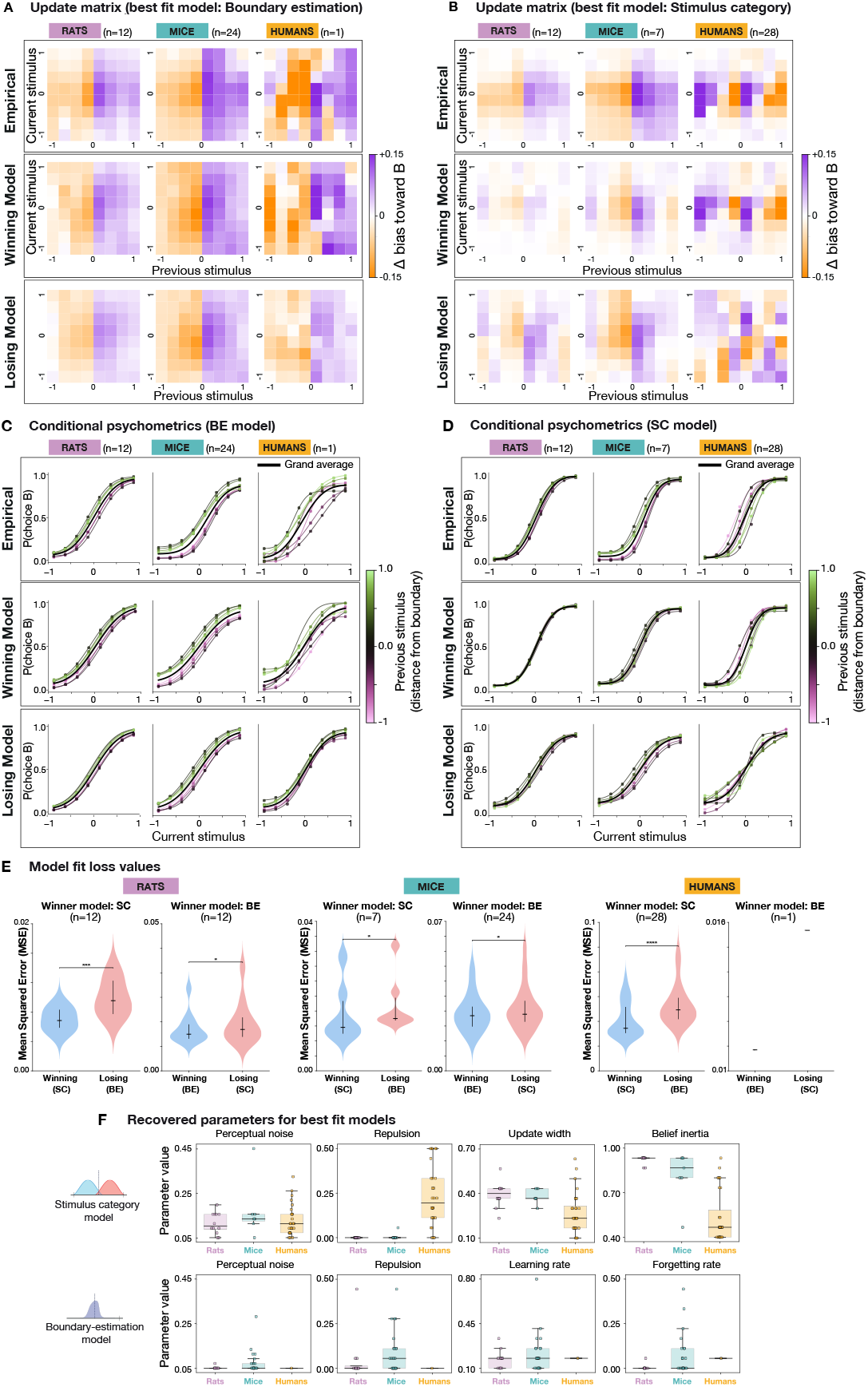
Update matrices and conditional psychometrics. **A–B**: Update matrices as in Figure 4, with an additional third row showing predictions from the *losing* model for each species (top: empirical data; middle: winning model prediction; bottom: losing model prediction). **C–D**, Full conditional psychometric functions for each species, conditioned on the previous trial’s stimulus (binned into 8 levels between –1 and 1 relative to the boundary). Top: empirical data; middle: winning model prediction. bottom: losing model prediction. **E**: Loss values (Mean Squared Error between model’s prediction and empirical conditional psychometric behaviour), for best fit models, showing the winning model in blue and losing model in red. MES values are significantly different between losing and winning models (paired t-test, all p-values <0.05) **F**: Best-fit parameter estimates for the BE and SC models, shown for each species.

**Figure S6.**
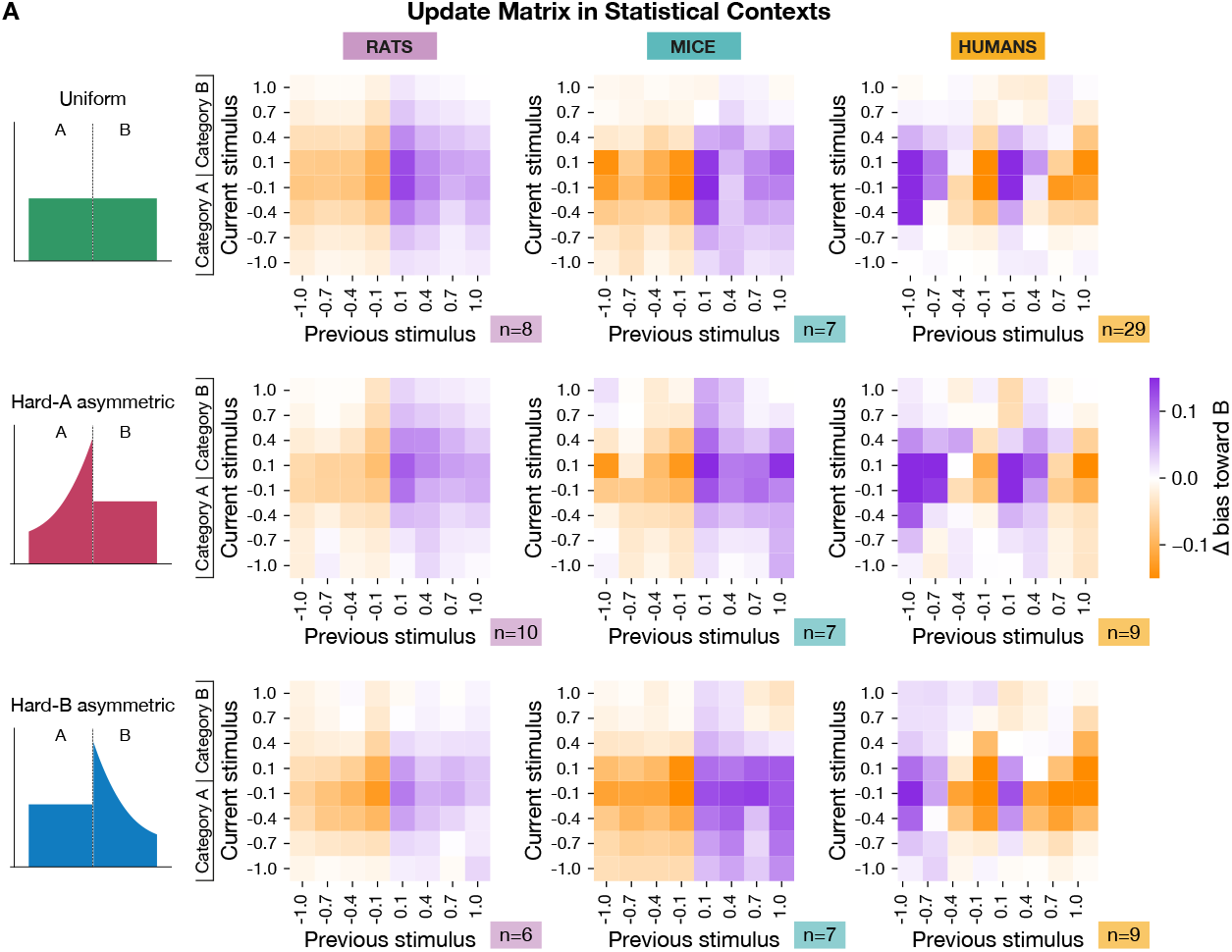
Average update matrices for uniform, hard-A, and hard-B distributions. **A**: Top panels show empirical update matrices for rats, mice, and humans in the uniform distribution condition. Middle panels show same format as in top panels, for the hard-A distribution. Bottom panels show same format as in top panels, for the hard-B distribution. These matrices capture how choice probabilities on the current trial depend on the stimulus and outcome of the previous trial, revealing species-specific patterns of trial-to-trial updating across contexts.

**Figure S7.**
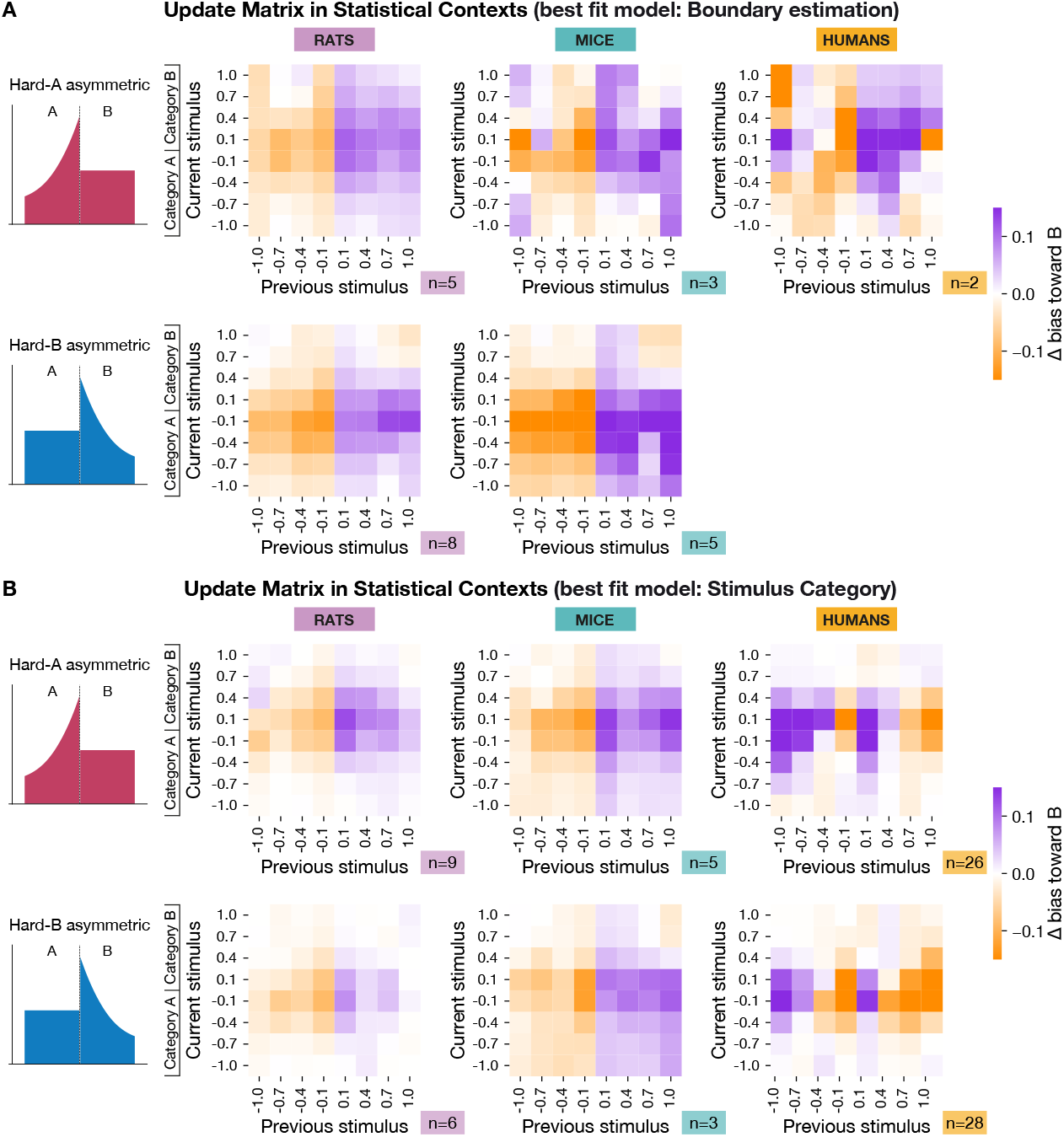
Average update matrices by best-fit model (BE vs. SC). **A**: Empirical update matrices for all individuals best fit by the Boundary-Estimation (BE) model, shown separately for hard-A (top row) and hard-B (bottom row), across rats, mice, and humans. **B**: Same format as in (A), but for individuals best fit by the Stimulus-Category (SC) model.

**Figure S8.**
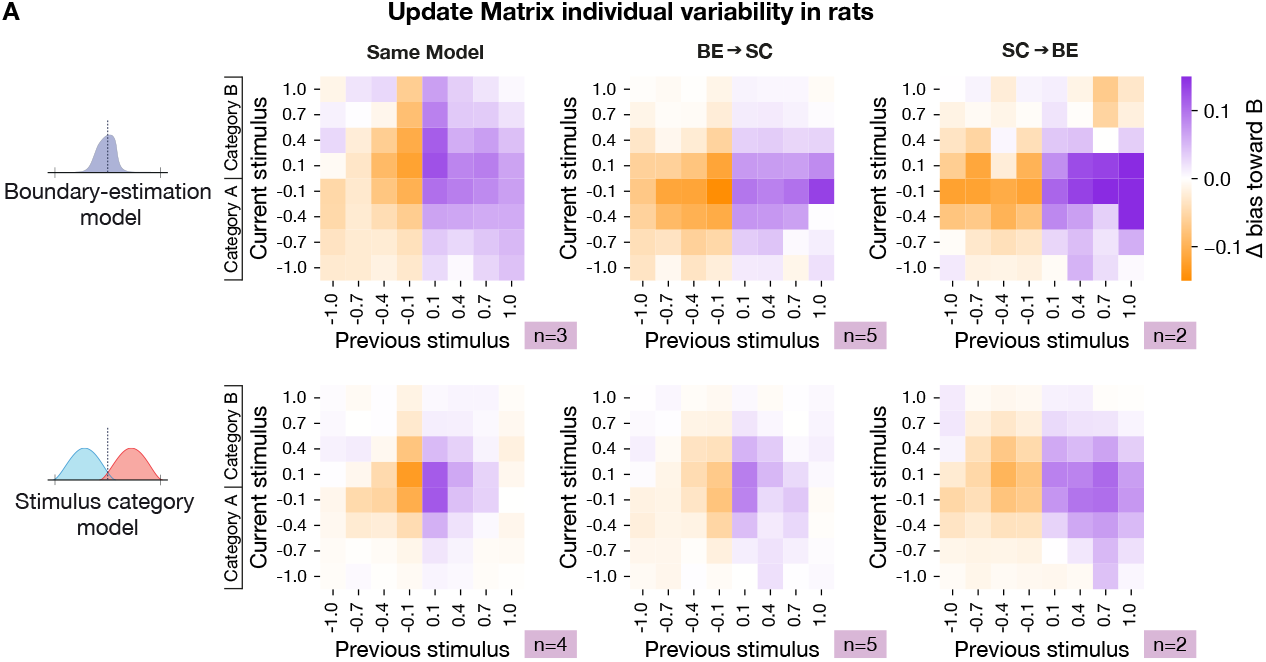
Average update matrices by best-fit model (BE vs. SC) in context-switching experiments in rats. **A**: Empirical update matrices for rats split by model-transition status across hard-A and hard-B distributions, shown separately for best-fit models Boundary-Estimation (BE) model (top) and Stimulus-Category (SC) model (bottom). Same best-fit model in both contexts (same model, left); best-fit model changes from the BE model in the first context to the SC model in the second context (BE→SC, middle); best-fit model changes from the SC model in the first context to the BE model in the second context (SC→BE, right). These results indicate substantial *between-rat* heterogeneity in updating strategies, consistent with model-specific (BE vs. SC) trial-to-trial updating patterns, and *within-rat* variability across time/experience and contexts.

**Figure S9.**
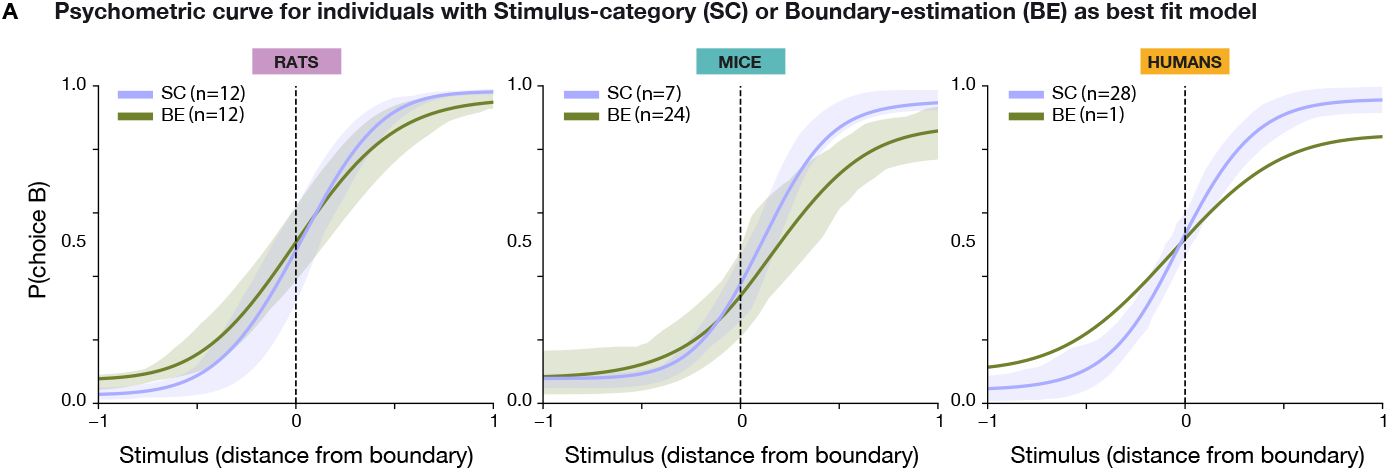
Psychometric curves for individuals with Stimulus-category (SC) vs Boundary-estimation (BE) as best fit models. Left: psychometric curves for rats with SC as best fit model (in purple) vs rats with BE as best fit model (in green). Probability of choosing category B is fit using a 4-parameters psychometric function (see Psychometric Curve fitting). Errorbars show SEM. Middle: the same for mice. Right: the same for humans.

**Figure S10.**
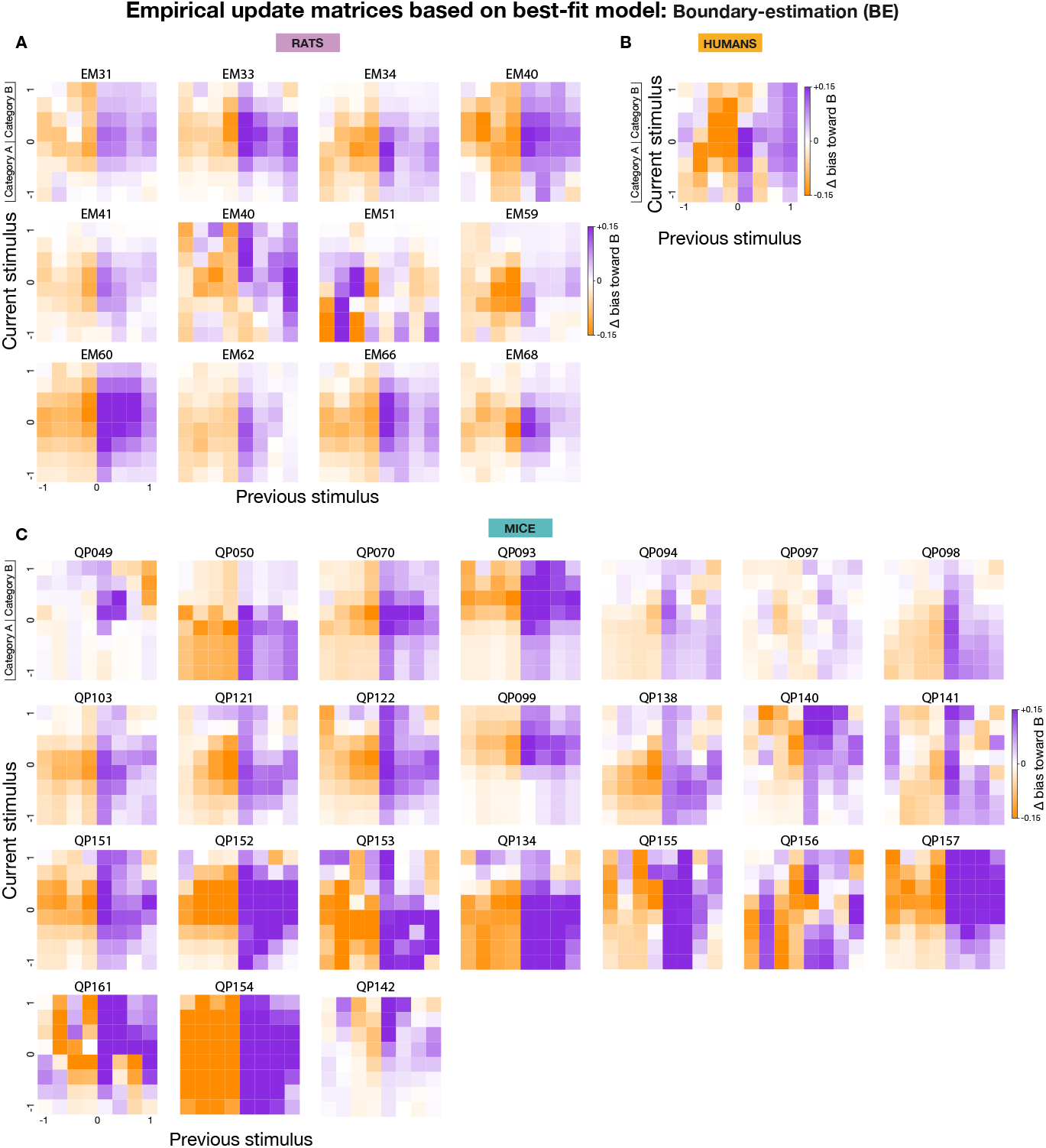
Empirical update matrices for individuals with BE as best fit model. **A**: Individual updated matrices for rats with BE as the winning model. Each rat has been exposed to one of the hard-A or hard-B asymmetric distribution (n = 12 rats out of total of 24) **B**: Similar to **A** for humans. Humans were exposed to uniform distribution only (n = 1 out of total of 29). **C**: Similar to **A** for mice. Mice were first trained on uniform distribution, followed by exposure to hard-A or hard-B or both distributions (n = 24 out of total of 31).

**Figure S11.**
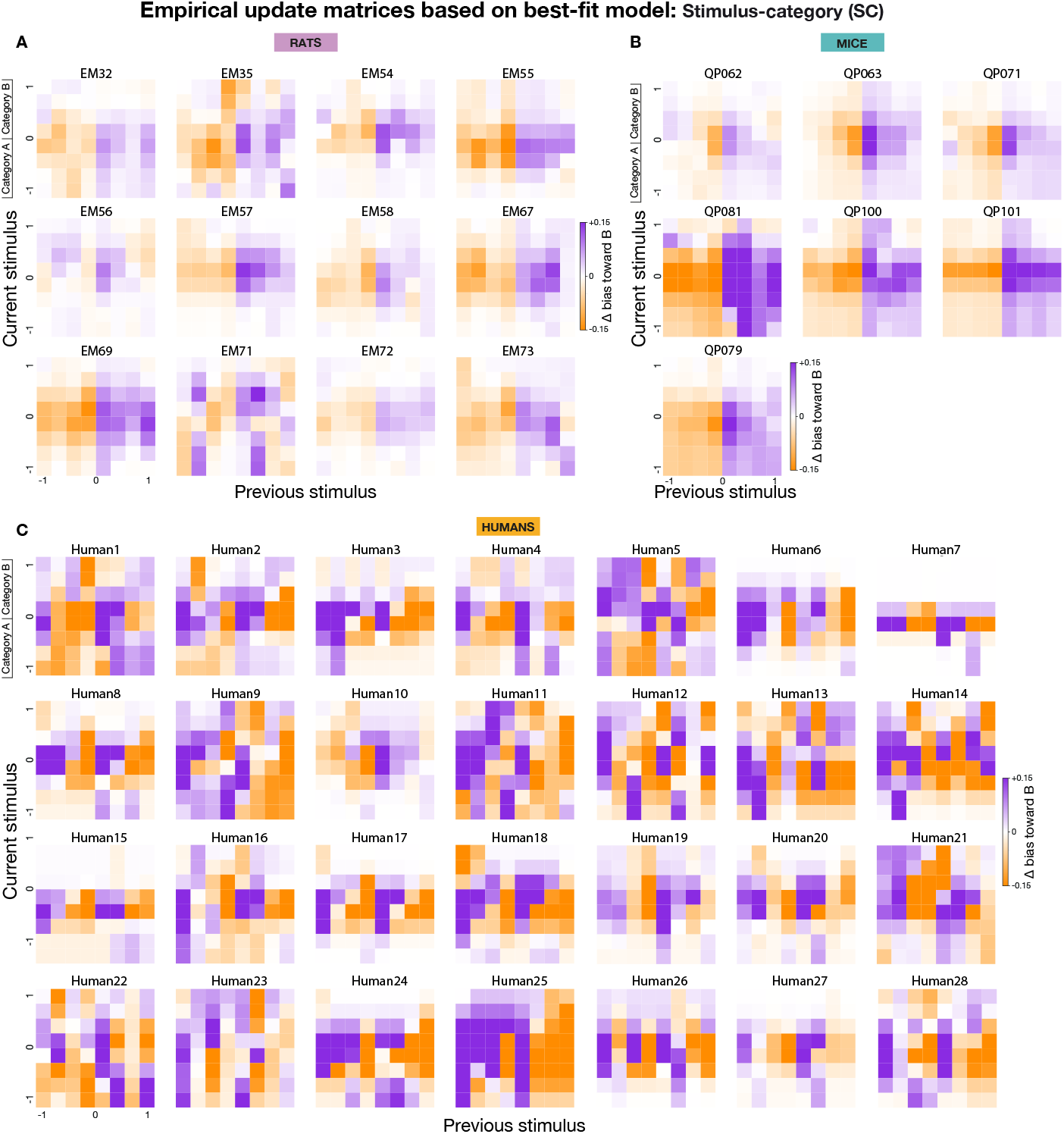
Empirical update matrices for individuals with SC as best fit model. **A**: Individual updated matrices for rats with SC as the winning model. Each rat has been exposed to one of the hard-A or hard-B asymmetric distribution (n = 12 rats out of total of 24). **B**: Similar to **A** for mice. Mice were first trained on uniform distribution, followed by exposure to hard-A or hard-B or both distributions (n = 7 out of total of 31). **C**: Similar to **A** for humans. Humans were exposed to uniform distribution only (n = 28 out of total of 29).

**Figure S12.**
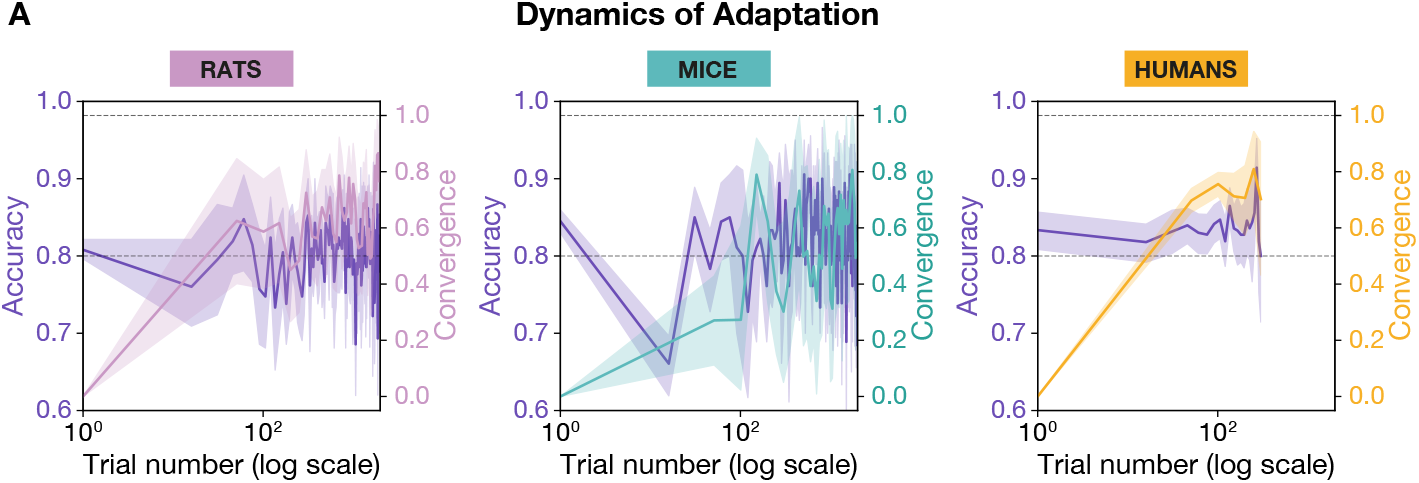
Convergence of psychometric bias toward the normative prediction maximises reward. **A**: Panels show results separately for rats (left), mice (middle), and humans (right). For each species, the left y-axis (purple line) shows empirical accuracy (reward rate) over time, while the right y-axis shows the convergence value (distance between observed and optimal PSE predicted by the normative model). Rats and mice show gradual convergence, with initial dips in accuracy followed by improvement as their PSE aligns with the optimal value. In contrast, humans converge rapidly: by 50 trials, the PSE is already near-optimal (reward rate computed in bins of 15 trials, PSE computed in bins of 50 trials). Together, these results indicate that convergence toward the normative PSE underlies reward maximisation across species, with species-specific differences in learning speed. Solid line: across-subject mean per bin for accuracy (left axis) and convergence (right axis). Shaded region: pointwise bootstrap confidence intervals.

**Figure S13.**
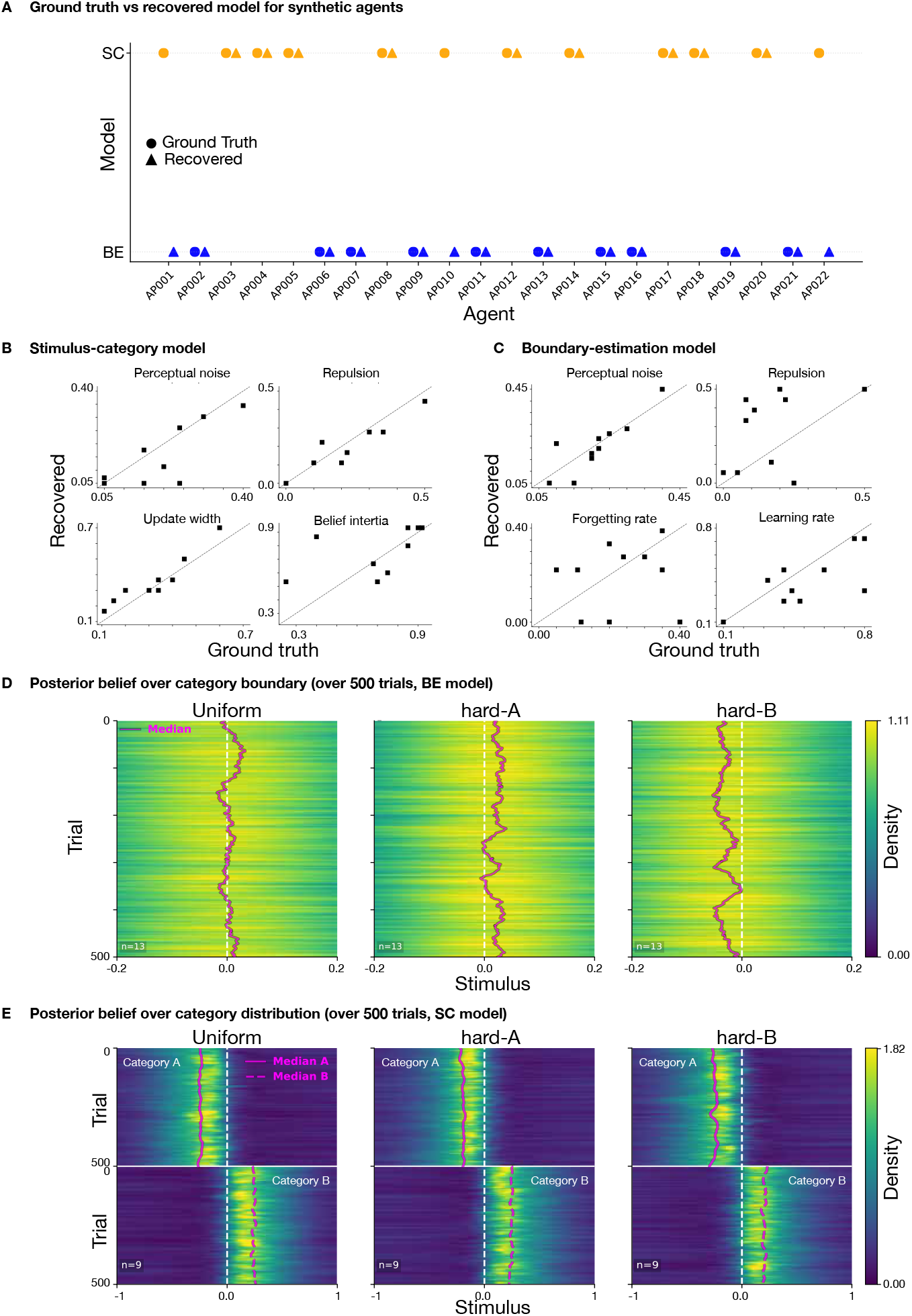
Model validation for Stimulus-category (SC) and Boundary-estimation (BE) models. **A**: True versus recovered model for each simulated agent. **B-D**: Parameter recovery for BE (left) and SC (right). Scatter plots compare recovered and ground-truth values for BE parameters (perceptual noise, repulsion magnitude, relaxation rate, learning rate) and SC parameters (perceptual noise, repulsion magnitude, update width, learning rate). D–E: Heatmaps show trial-by-trial posteriors over the first 500 trials in BE (D, n=13) and SC (E, n=9). At each trial and stimulus bin, posteriors were averaged across subjects on a common grid, so that the heatmaps display the mean posterior distributions for the group.

**Figure S14.**
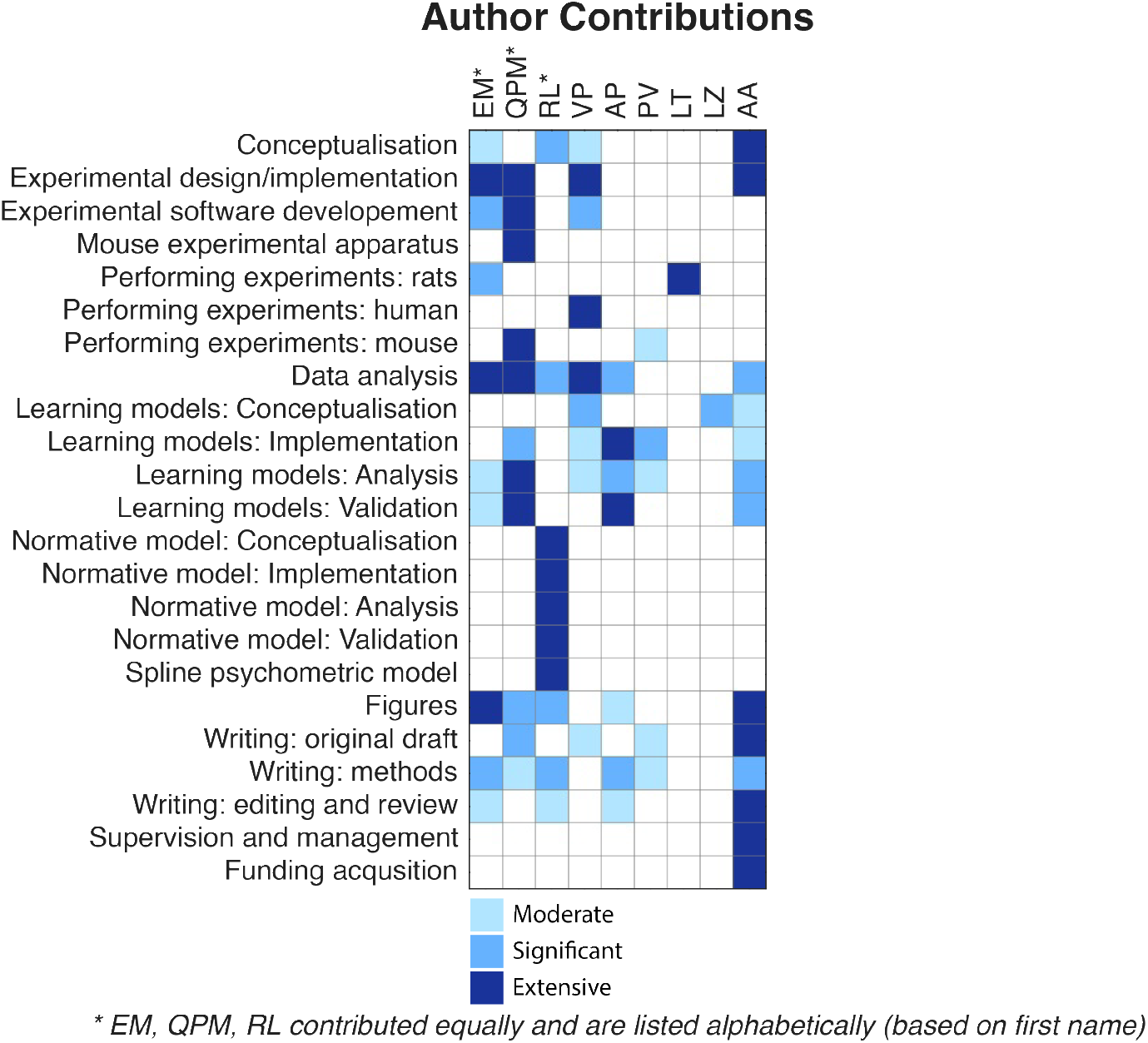
Author contribution table.

## Acknowledgment

The authors thank P. Latham, T. Behrens, E. Chong, A. Onih, A. Agarwal, S. Shentyurk, and V. Boboeva, for comments on the manuscript; the staff at the SWC Neurobiological Research Facility for animal support; the SWC Fabrication Laboratory for help with machining; K. Lee and A. Agarwal for help with training mice and rats; members of the Akrami, and other SWC laboratories for insightful discussion and advice. This work was supported by Wellcome (219880/Z/19/Z) to E.M., Wellcome (225438/Z/22/Z) to Q.P.M, Sainsbury Wellcome Centre’s core provided by Wellcome (219627/Z/19/Z), the Gatsby Charitable Foundation (GAT3755), and UK Research and Innovation grant to A.A. (EP/Z000599/1).

## Author Information

### Authors and affiliations

**Sainsbury Wellcome Centre, University College London** Elena Menichini, Quentin Pajot-Moric, Ryan Low, Victor Pedrosa, Amirali Pourdehghan, Peter Vincent, Liang Zhou, Lillianne Teachen, Athena Akrami

**Gatsby Computational Neuroscience Unit, University College London** Liang Zhou, Victor Pedrosa (current affiliation)

### Contributions

See the contribution table in Figure S14.

