## Supplemental Figures for "Different learning algorithms achieve shared optimal outcomes in humans, rats, and mice"

### Supplementary Figures

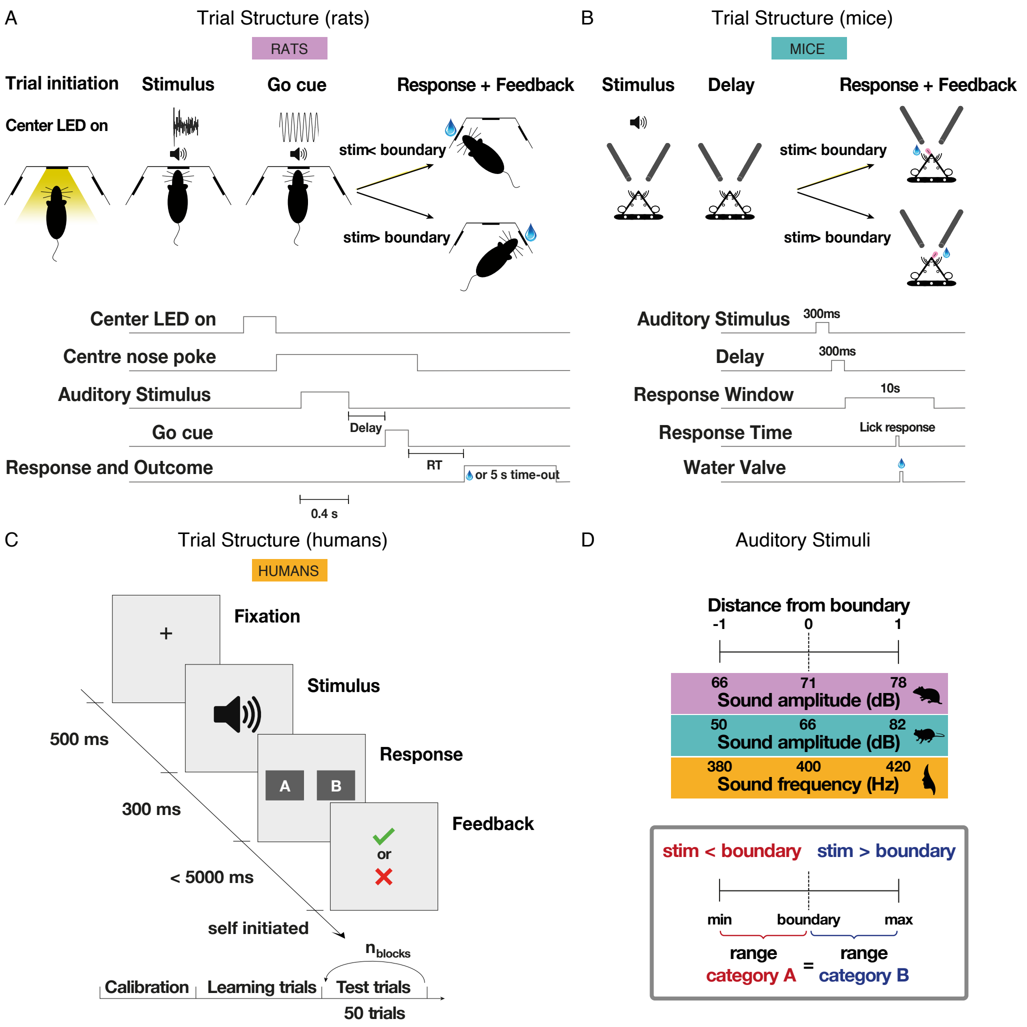


**Figure S1**: **Task schematics.** Full trial structure for each species (see Methods ). **A**: Trial structure for freely moving rats, illustrating stimulus presentation (structured noise of varying intensity), response, and feedback. **B**: Same as (A) for head-fixed mice. **C**: Trial structure for humans, presented with pure tones of varying frequency; session outline includes calibration, training, and test phases **D**: Mapping of "distance from boundary" onto the physical stimulus dimension: sound intensity (rats and mice) and sound frequency (humans). The decision boundary was fixed at the midpoint of the stimulus range (in logarithmic scale), which was sampled continuously.

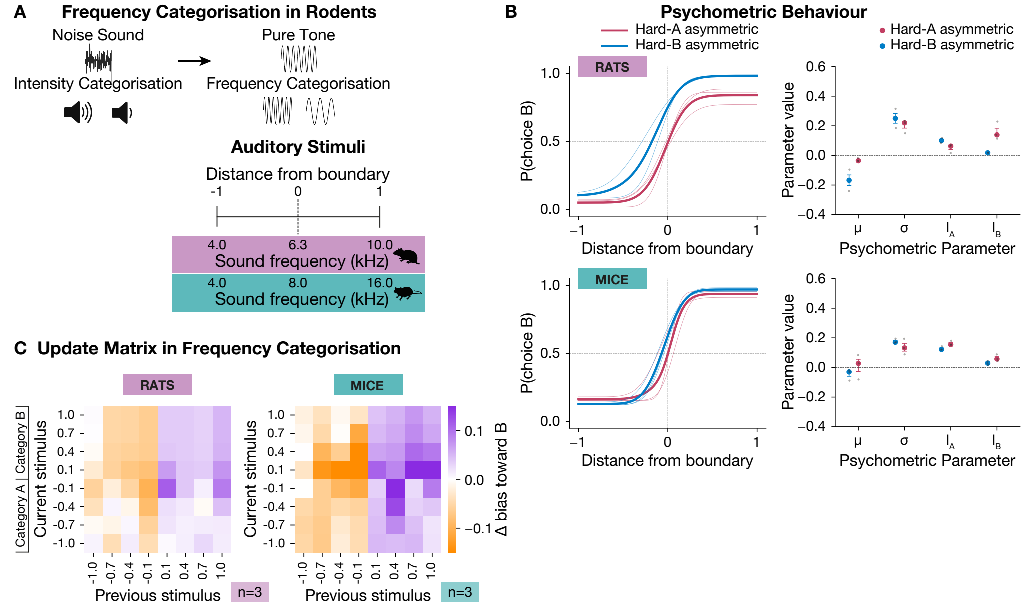


**Figure S2**: **Frequency version of the sound categorisation task for rodents**. Control experiment designed to match the stimulus dimension used in human subjects, who performed frequency rather than intensity categorisation. **A**: Task schematics and mapping of "distance from boundary" onto the physical stimulus dimension (sound frequency) for rats and mice. The decision boundary was fixed at the midpoint of the frequency range, which was sampled continuously. **B**: left panels show psychometric curves for all individuals (thin lines) and the average across-subject fit (thick line), shown separately for rats (top) and mice (bottom), for hard-A and hard-B contexts in frequency categorisation task, showing modulation similar to the intensity experiments. Right panels show individual parameter estimates (dots) from four-parameter psychometric fits (mean, slope, lapse rates) for hard-A and hard-B, alongside the group median and interquartile range. **C**: Empirical update matrices for rats (left) and mice (right) in the uniform context, showing choice updating patterns similar to the intensity experiments for each species.

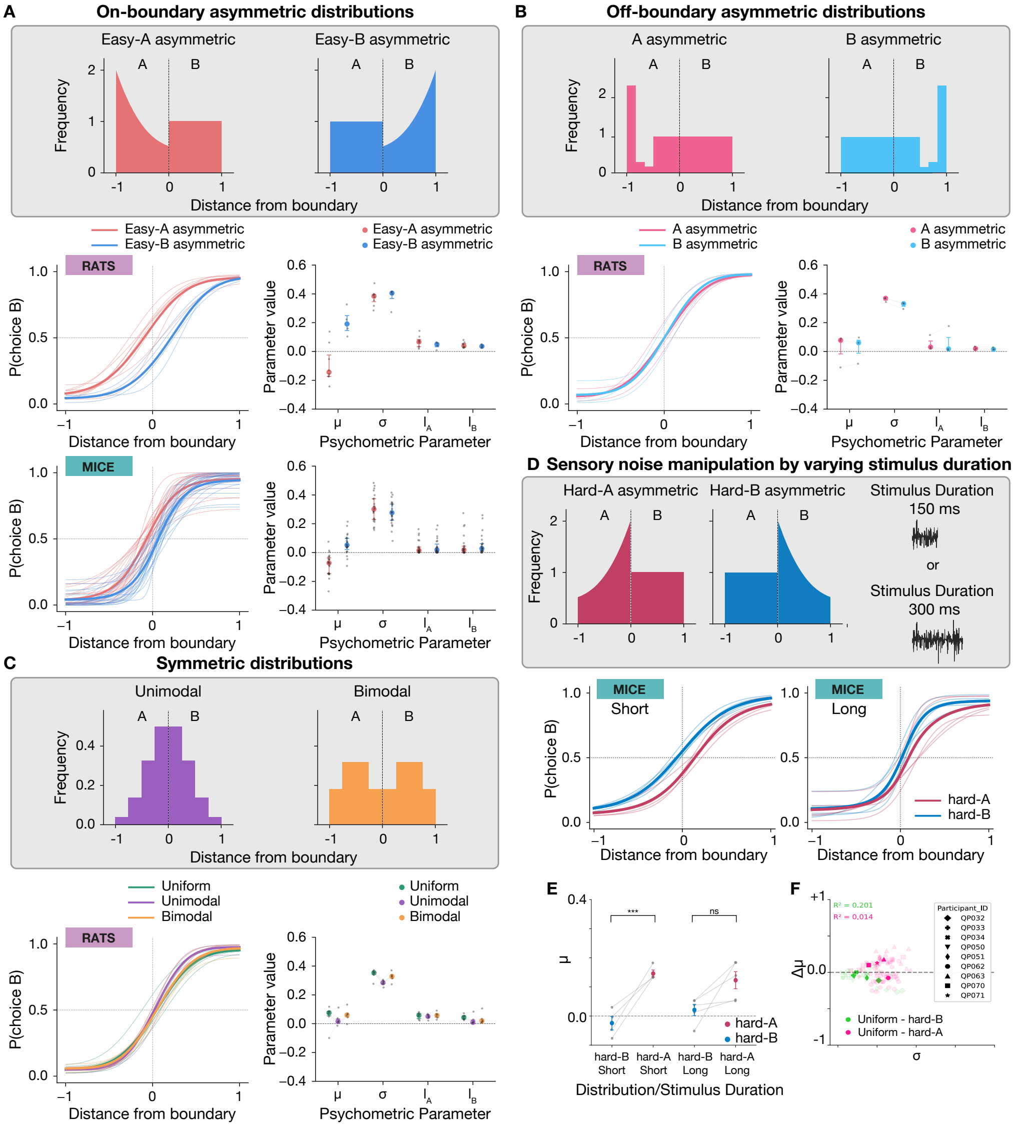


**Figure S3**: **Asymmetry in sensory distribution is exploited only if needed**. Biases observed in humans and rodents are solely due to the relative difference in category representation around the boundary irrespective of the shape of the non-uniform category. **A**: easy-A and easy-B contexts, in which the non-uniform category over-represented easy stimuli (far from the boundary) and under-represented hard stimuli (near the boundary). The corresponding psychometric curves for rats and humans are shown below: left panels show psychometric curves for all individuals (thin lines) and the average across-subject fit (thick line), shown separately by species and for the stimulus distributions. Right panels show individual parameter estimates from the four-parameter fits, together with the group median and interquartile range. Both humans and rats in the easy-A condition showed a bias toward category A for boundary-adjacent stimuli, while easy-B produced the opposite bias. Mann–Whitney $U$ test, mu in rats: p-value = 0.01379 for easy-A (n=8) vs easy-B(n=4); Wilcoxon signed-rank test, mu in humans (n=23): p-value = 0.0000002 for easy-A vs easy-B. **B**: Off-boundary asymmetric contexts. Statistical contexts with asymmetry away from decision boundary within category A (pink) or category B (light blue), relative to the other uniformly-sampled category. Left panel shows psychometric curves for all individuals (thin lines) and the average across-subject fit (thick line), shown for the stimulus distributions. Right panel shows individual parameter estimates from the four-parameter fits, together with the group mean ± SEM. The psychometric curves did not show a choice bias in the off-boundary asymmetrical contexts and the mean parameter of the psychometric curve fits $\mu$ was around zero for both distributions, indicative of a balanced performance, similar to the performance of subjects in the uniform context. Two-sided paired t-test, mu in rats (n=3): p-value = 0.72492 for A asymmetric vs B asymmetric. **C**: Over- or under-exposure to hard stimuli did not alter human or rat performances. Contexts with symmetric distributions of stimuli lead to oversampling of either easy (bimodal) or difficult (unimodal). Left panel shows psychometric curves for all individuals (thin lines) and the average across-subject fit (thick line), shown separately for the stimulus distributions. Right panel shows individual parameter estimates from the four-parameter fits, together with the group median and interquartile range. These manipulations do not vary overall performance in rats. Wilcoxon signed-rank test, Bonferroni corrected, mu in rats (n=6): p-value = 0.87500 for uniform vs unimodal; p-value = 1.00000 for uniform vs bimodal.

**Figure S3**: **Continued. D**: Varying stimulus duration changes the saliency of sensory inputs, hence the perceptual reliability and noise. Bottom panels show psychometric curves for all individuals (thin lines) and the average across-subject fit (thick line), shown separately by stimulus durations and for the stimulus distributions. **E**: paired $\mu$ estimates from psychometric fits for each individual, pairing hard-A with hard-B within subject; circles indicate the group mean $\pm$ SEM, and grey lines connect within-subject pairs, shown separately for stimulus-duration cohorts. Psychometric shifts tended to be smaller for longer stimulus durations (i.e., reduced sensory noise), consistent with the prediction of the normative model (paired t-test with bonferroni correction ($\alpha = 0.05$), p = 0.010 for short duration, n = 4; paired t-test with bonferroni correction ($\alpha = 0.05$), p = 0.058 for long duration, n = 5). Short-duration cohort have significantly steeper slopes than long duration cohort (Short mean = 0.265, Long mean = 0.150; t(7) = 2.86, p = 0.024, Hedges' g = 1.71) **F**: $\Delta\mu$ (shift in the psychometric mean value) vs. $\sigma$ (reverse of slope) is shown for those 7 mice, when changing the distribution from uniform to hard-A. This suggests the shallower the slope, the larger the shift in the psychometric mean value.

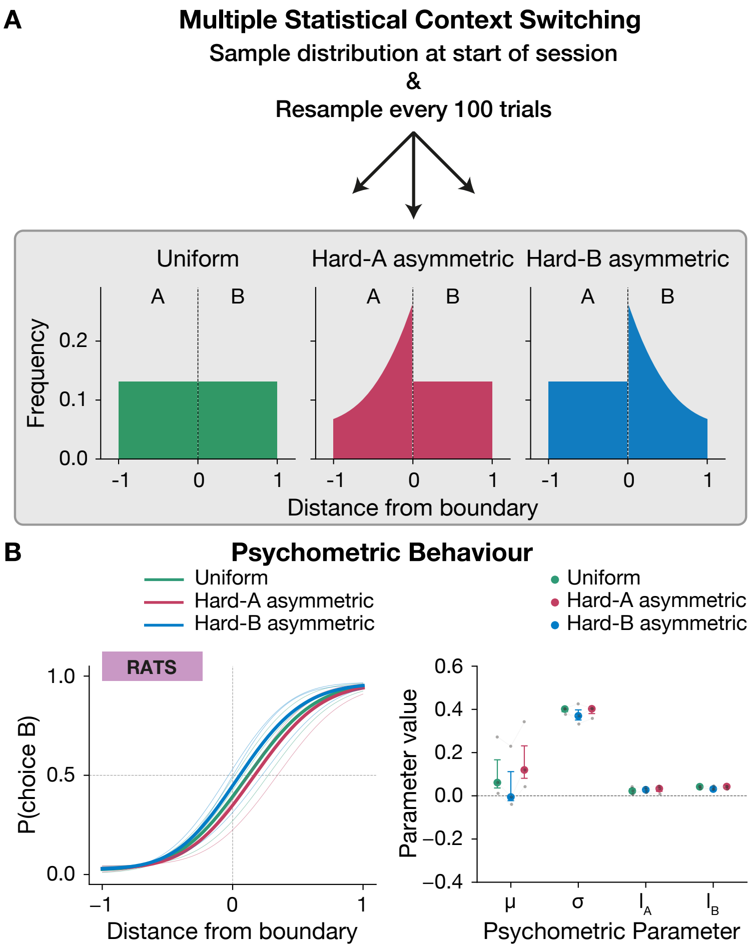


**Figure S4**: **Within-session probabilistic context switching in rats**. **A**: Sampling distributions used in the probabilistic switching paradigm (same as in Figure 1. **B**: Left panels show psychometric curves for all individuals (thin lines) and the average across-subject data (thick line), shown separately for each distribution in **A**. Right panels show individual parameter estimates (dots) from psychometric fits—midpoint, slope (width), and lapse rate—together with group median and interquartile range. In this within-session paradigm, analogous to that used in humans, rats adapted flexibly to the prevailing distribution, comparable to the static-context condition (see Figures 3 & 5).

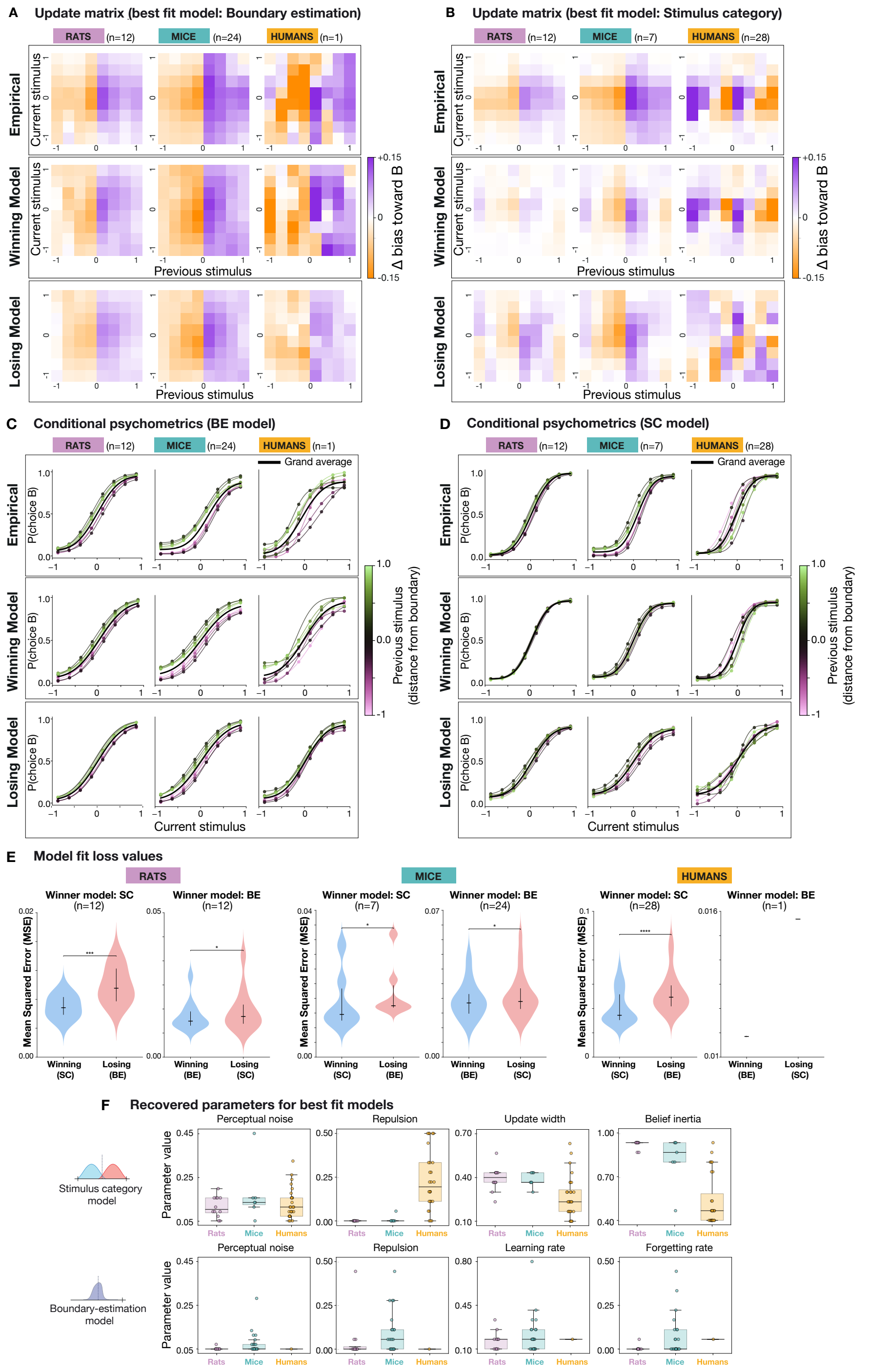


**Figure S5**: **Update matrices and conditional psychometrics**. **A–B**: Update matrices as in Figure 4, with an additional third row showing predictions from the *losing* model for each species (top: empirical data; middle: winning model prediction; bottom: losing model prediction). **C–D**, Full conditional psychometric functions for each species, conditioned on the previous trial's stimulus (binned into 8 levels between –1 and 1 relative to the boundary). Top: empirical data; middle: winning model prediction. bottom: losing model prediction. **E**: Loss values (Mean Squared Error between model's prediction and empirical conditional psychometric behaviour), for best fit models, showing the winning model in blue and losing model in red. MES values are significantly different between losing and winning models (paired t-test, all p-values <0.05) **F**: Best-fit parameter estimates for the BE and SC models, shown for each species.

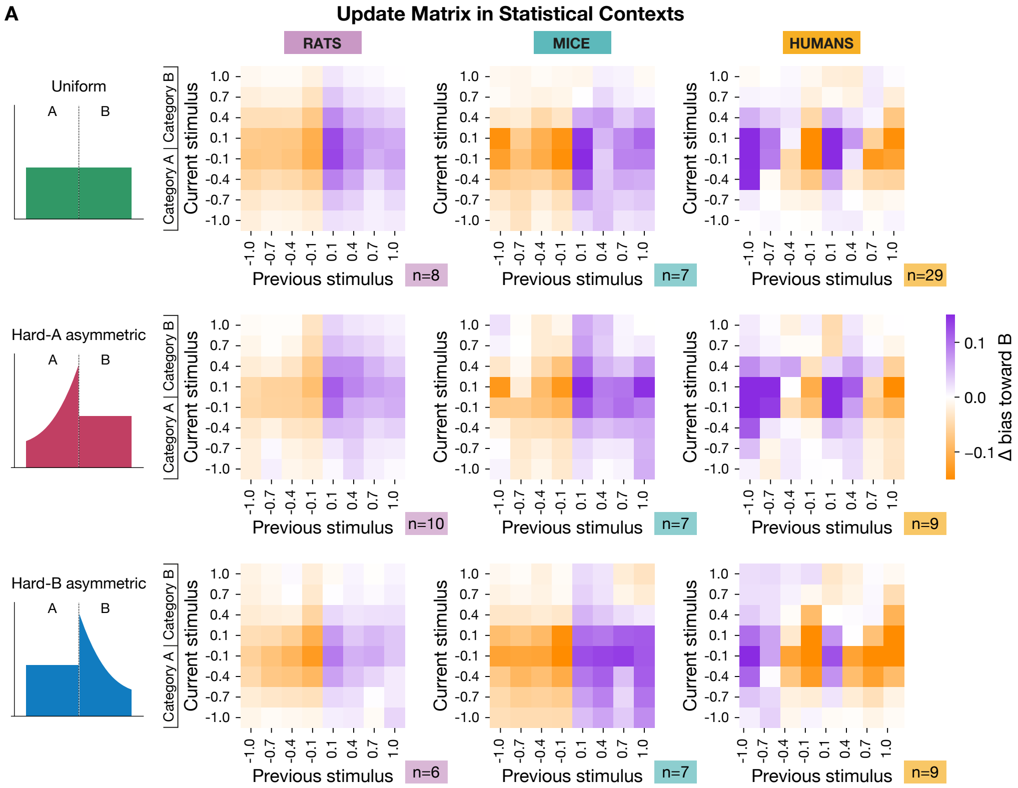


**Figure S6**: **Average update matrices for uniform, hard-A, and hard-B distributions.** **A**: Top panels show empirical update matrices for rats, mice, and humans in the uniform distribution condition. Middle panels show same format as in top panels, for the hard-A distribution. Bottom panels show same format as in top panels, for the hard-B distribution. These matrices capture how choice probabilities on the current trial depend on the stimulus and outcome of the previous trial, revealing species-specific patterns of trial-to-trial updating across contexts.

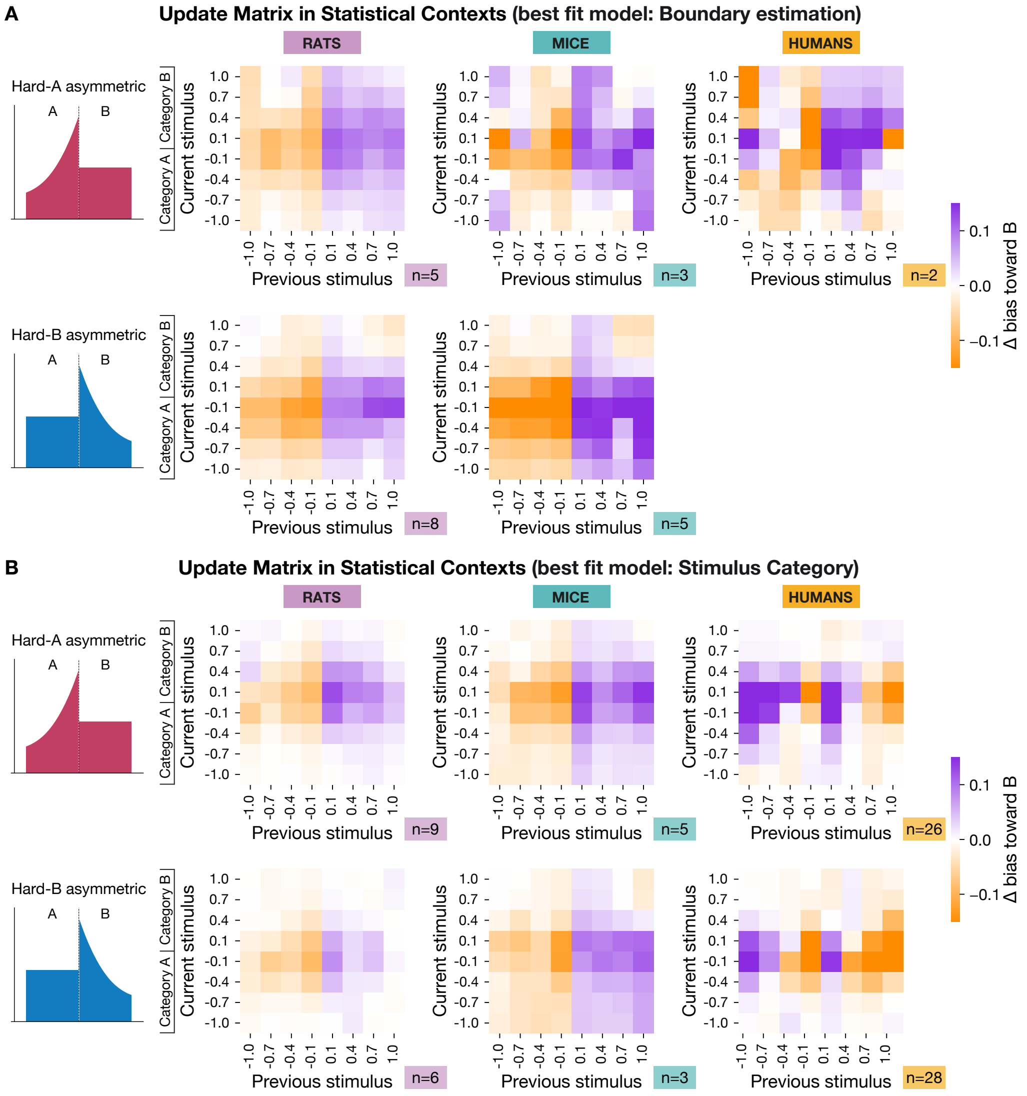


**Figure S7**: **Average update matrices by best-fit model (BE vs. SC).** **A**: Empirical update matrices for all individuals best fit by the Boundary-Estimation (BE) model, shown separately for hard-A (top row) and hard-B (bottom row), across rats, mice, and humans. **B**: Same format as in (A), but for individuals best fit by the Stimulus-Category (SC) model.

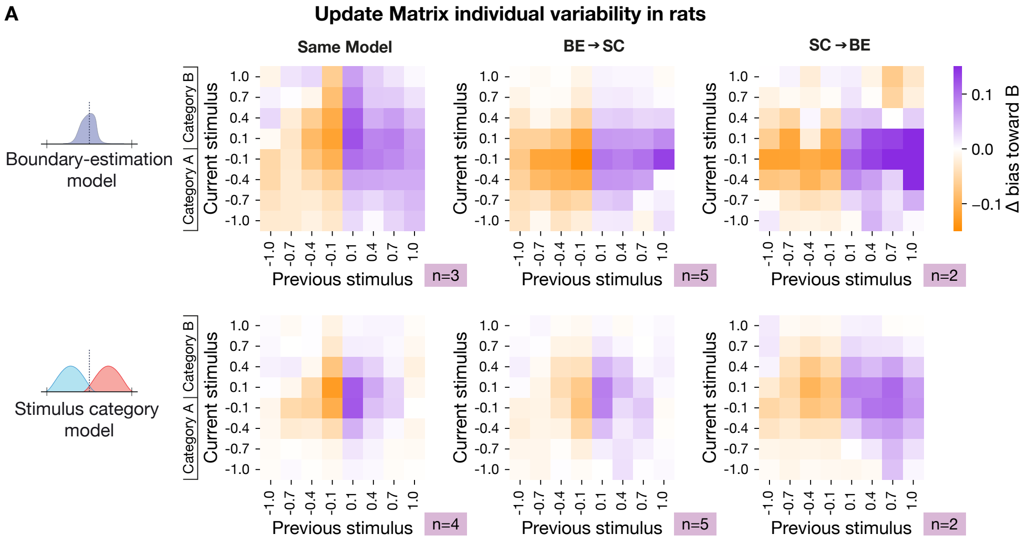


**Figure S8**: **Average update matrices by best-fit model (BE vs. SC) in context-switching experiments in rats.** **A**: Empirical update matrices for rats split by model-transition status across hard-A and hard-B distributions, shown separately for best-fit models Boundary-Estimation (BE) model (top) and Stimulus-Category (SC) model (bottom). Same best-fit model in both contexts (same model, left); best-fit model changes from the BE model in the first context to the SC model in the second context (BE→SC, middle); best-fit model changes from the SC model in the first context to the BE model in the second context (SC→BE, right). These results indicate substantial *between-rat* heterogeneity in updating strategies, consistent with model-specific (BE vs. SC) trial-to-trial updating patterns, and *within-rat* variability across time/experience and contexts.

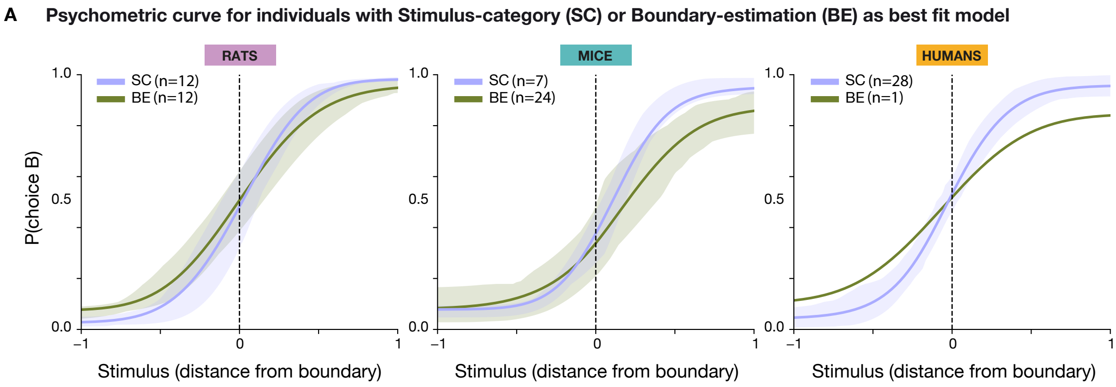


**Figure S9**: **Psychometric curves for individuals with Stimulus-category (SC) vs Boundary-estimation (BE) as best fit models.** Left: psychometric curves for rats with SC as best fit model (in purple) vs rats with BE as best fit model (in green). Probability of choosing category B is fit using a 4-parameters psychometric function (see Psychometric Curve fitting). Errorbars show SEM. Middle: the same for mice. Right: the same for humans.

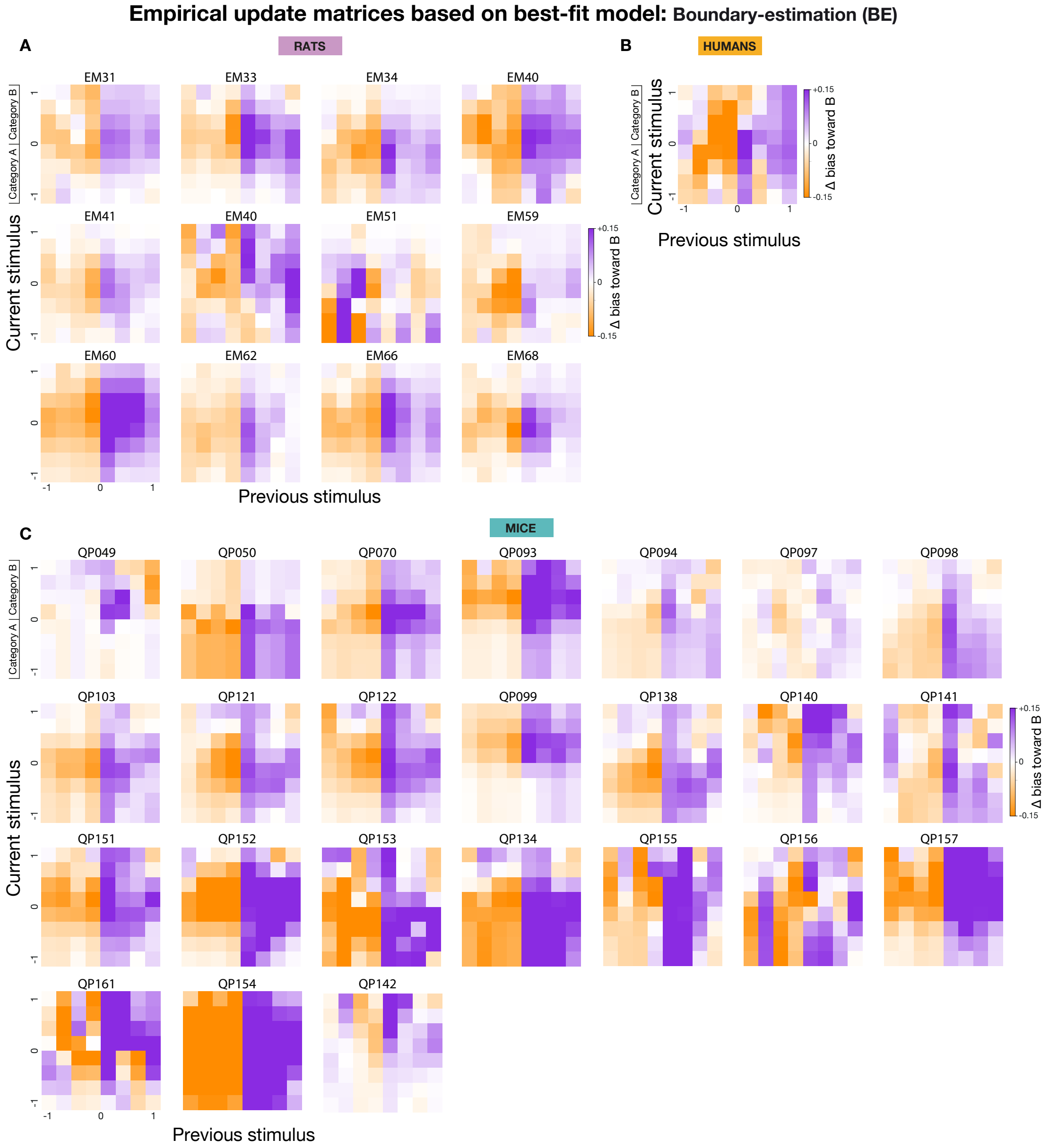


**Figure S10**: **Empirical update matrices for individuals with BE as best fit model.** **A**: Individual updated matrices for rats with BE as the winning model. Each rat has been exposed to one of the hard-A or hard-B asymmetric distribution (n = 12 rats out of total of 24) **B**: Similar to **A** for humans. Humans were exposed to uniform distribution only (n = 1 out of total of 29). **C**: Similar to **A** for mice. Mice were first trained on uniform distribution, followed by exposure to hard-A or hard-B or both distributions (n = 24 out of total of 31).

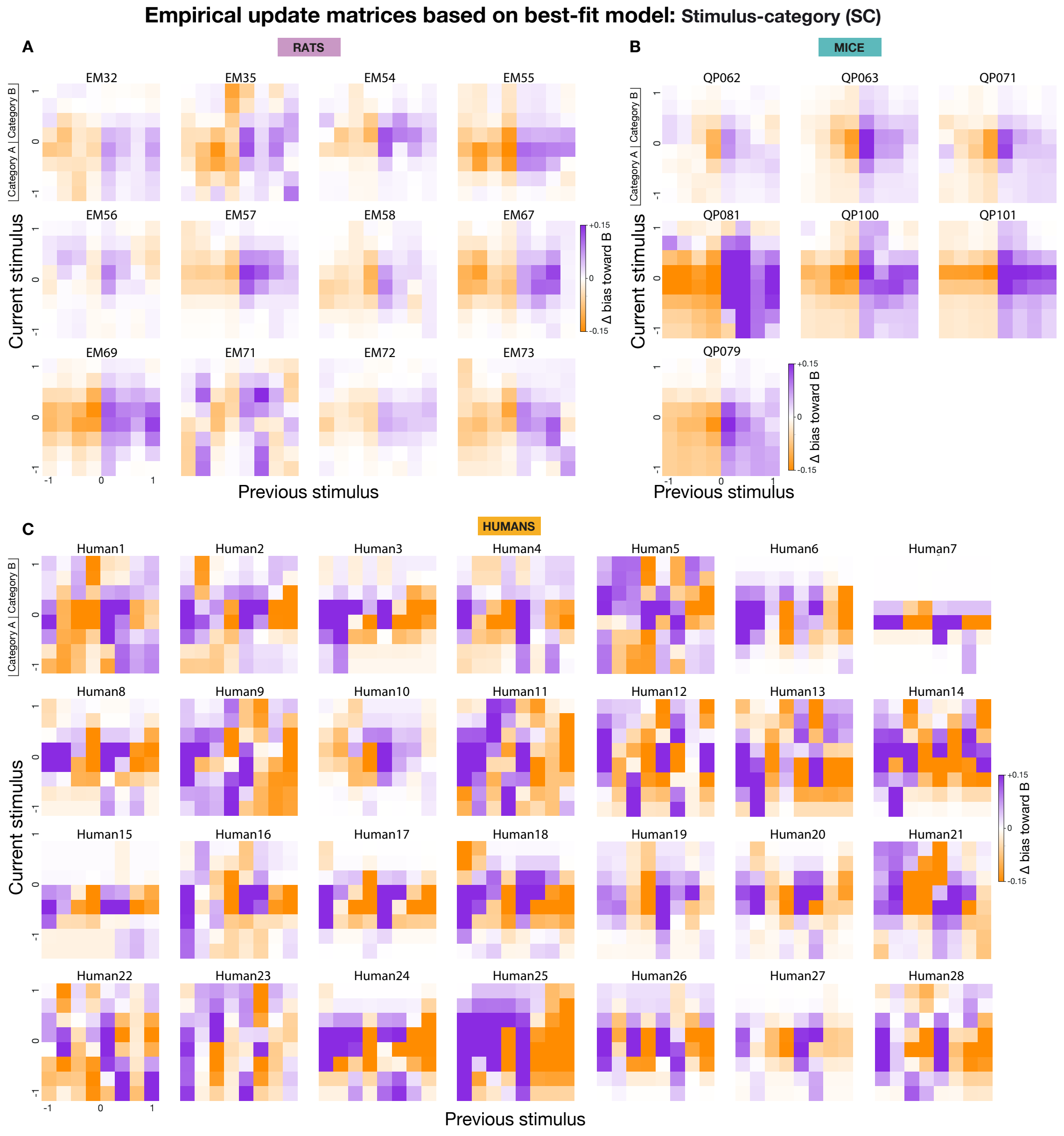


**Figure S11**: **Empirical update matrices for individuals with SC as best fit model.** **A**: Individual updated matrices for rats with SC as the winning model. Each rat has been exposed to one of the hard-A or hard-B asymmetric distribution (n = 12 rats out of total of 24). **B**: Similar to **A** for mice. Mice were first trained on uniform distribution, followed by exposure to hard-A or hard-B or both distributions (n = 7 out of total of 31). **C**: Similar to **A** for humans. Humans were exposed to uniform distribution only (n = 28 out of total of 29).

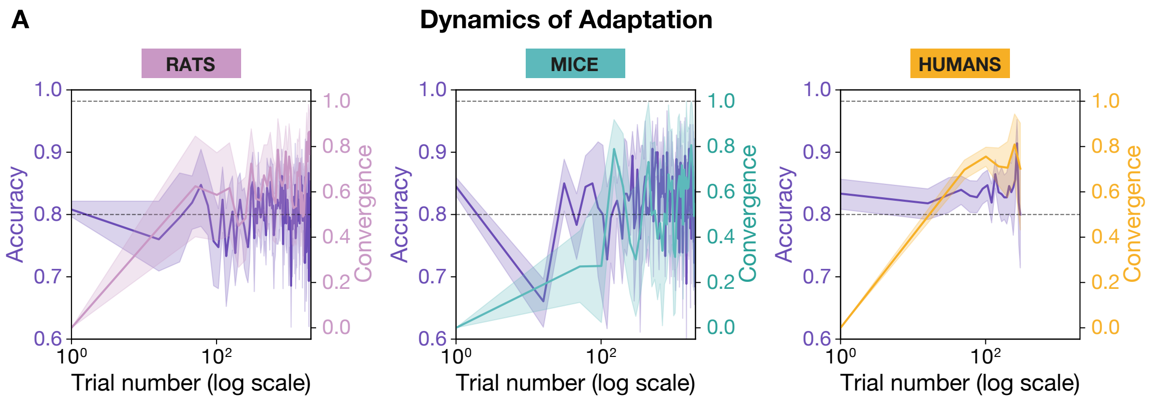


**Figure S12**: **Convergence of psychometric bias toward the normative prediction maximises reward. A**: Panels show results separately for rats (left), mice (middle), and humans (right). For each species, the left y-axis (purple line) shows empirical accuracy (reward rate) over time, while the right y-axis shows the convergence value (distance between observed and optimal PSE predicted by the normative model). Rats and mice show gradual convergence, with initial dips in accuracy followed by improvement as their PSE aligns with the optimal value. In contrast, humans converge rapidly: by 50 trials, the PSE is already near-optimal (reward rate computed in bins of 15 trials, PSE computed in bins of 50 trials). Together, these results indicate that convergence toward the normative PSE underlies reward maximisation across species, with species-specific differences in learning speed. Solid line: across-subject mean per bin for accuracy (left axis) and convergence (right axis). Shaded region: pointwise bootstrap confidence intervals.

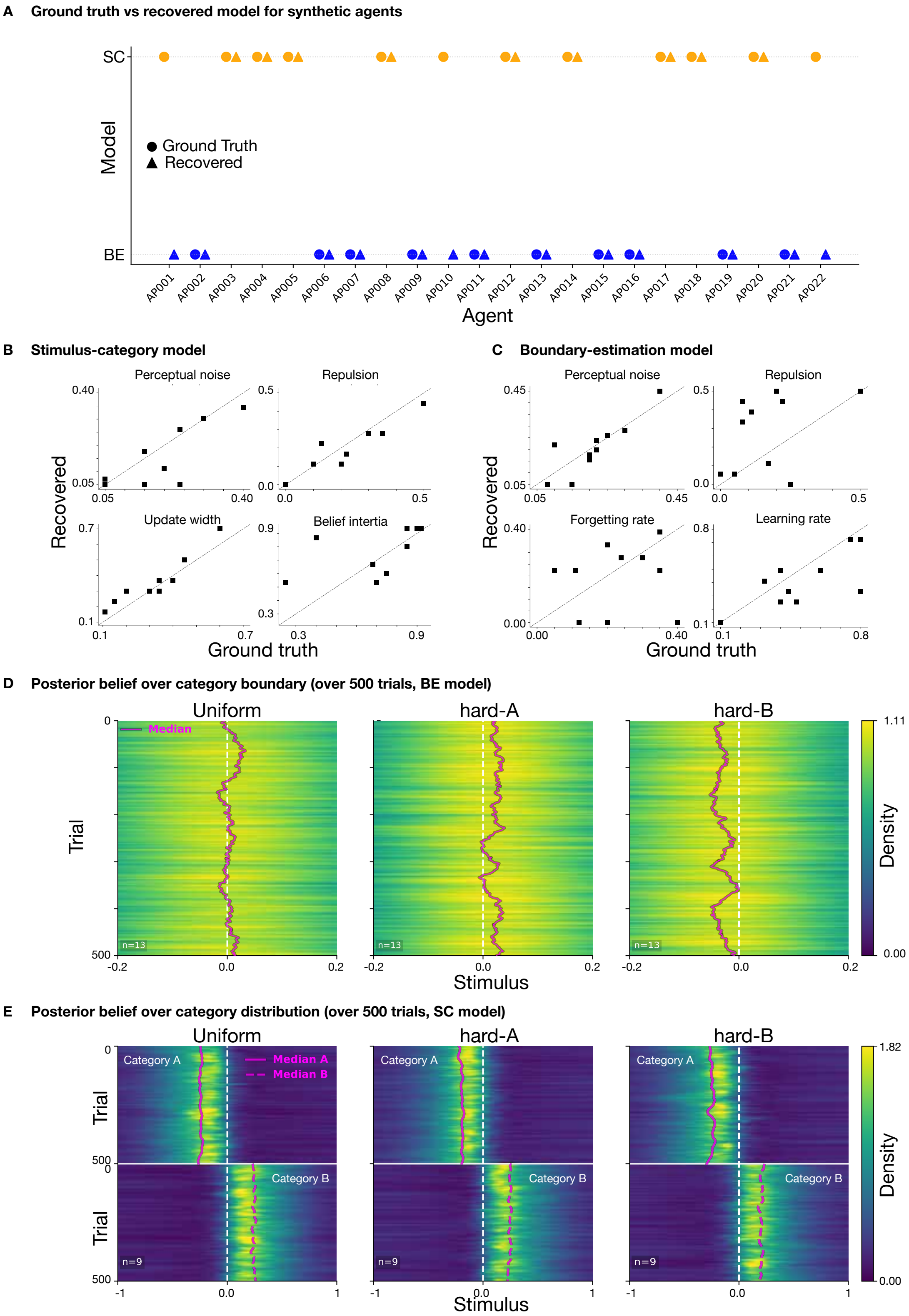


**Figure S13**: **Model validation for Stimulus-category (SC) and Boundary-estimation (BE) models. A**: True versus recovered model for each simulated agent. **B-D**: Parameter recovery for BE (left) and SC (right). Scatter plots compare recovered and ground-truth values for BE parameters (perceptual noise, repulsion magnitude, relaxation rate, learning rate) and SC parameters (perceptual noise, repulsion magnitude, update width, learning rate). D–E: Heatmaps show trial-by-trial posteriors over the first 500 trials in BE (D, n=13) and SC (E, n=9). At each trial and stimulus bin, posteriors were averaged across subjects on a common grid, so that the heatmaps display the mean posterior distributions for the group.
